# Identification of JMJD1C as a key mediator of fibroblast remodelling in human AF by single-nucleus multi-omics

**DOI:** 10.64898/2026.11.12.765924

**Authors:** Yile Fu, Lirong Li, Michiel R. L. Tubeeckx, Alessia Virzì, Samaneh Ekhteraei-Tousi, Justine Troffaes, Bernard Thienpont, Gilles W. De Keulenaer, Hein Heidbuchel, Vincent F. M. Segers, H. Llewelyn Roderick

## Abstract

Atrial fibrillation (AF) is the most prevalent arrhythmia, accounting for up to 15% of arrhythmias in the elderly. Atrial enlargement associated with fibrotic and hypertrophic remodelling, termed atrial myopathy (AM), is considered a precursor to AF development. The mechanisms underlying the development of AM and its progression to AF remain incompletely resolved. To identify the unique and combined contributions of the major cellular mediators of AM and AF, we combine single-nucleus transcriptomics of human left and right atrial appendages (LAA and RAA) with paired RNA and chromatin-accessibility profiling of 20 LAA samples. We validate these findings through histological and functional studies in primary human cardiac fibroblasts (HCFs). Patients were stratified according to the presence of AF and AM (identified by left atrial volume dimensions; LAVI, Hi and Lo), generating Hi-AF, Hi-SR, Lo-AF and Lo-SR groups that separate AF-associated from structural-remodelling-associated alterations. Across 75,654 qualified nuclei, AF was associated with chamber-dependent shifts in cellular composition and gene expression, with extracellular matrix (ECM) and TGF-β programmes prominent in LAA. Multiome analysis identified a stressed atrial cardiomyocyte (aCM4) state expressing natriuretic peptide A (NPPA) and natriuretic peptide B (NPPB), with altered *cis*-regulatory connectivity, and an inflammatory fibroblast (PTGDS^+^ FB) state characterised by *IL-6* expression and matrix- and TGF-β-associated programmes. Enhancer-driven network analysis identified *GLIS3*, *TWIST2* and *TBX20* as regulators of FB state transitions. The histone demethylase *JMJD1C* was enriched in PTGDS^+^ FB and induced by TGF-β1; its depletion attenuated α-smooth-muscle-actin (α-SMA) accumulation and reprogrammed TGF-β1-responsive transcription. Ligand–receptor analysis further predicted macrophage-to-FB TGF-β signalling, showing the contribution of macrophages to the atrial inflammatory environment. These data define discrete aspects of rhythm-and structure-associated remodelling of human atrial tissue and identify a candidate macrophage–TGF-β–JMJD1C–FB interaction for potential therapeutic development.

## Introduction

Atrial fibrillation (AF) is the most common sustained cardiac arrhythmia and a major cause of stroke, heart failure (HF), cognitive decline, and premature mortality^1–4^. Its global burden continues to rise with population ageing and the increasing prevalence of cardiometabolic risk factors, despite substantial advances in anticoagulation, rhythm control, and catheter ablation^5,6^. Current treatments primarily address thromboembolic risk or the electrical abnormalities of AF, but recurrence and progression remain common since these therapies do not remove the pathophysiological substrate that sustains the arrhythmia^7^. The nature of this substrate and the mechanisms that contribute to AF evolution in humans are not fully understood, but their identification is necessary for the development of new therapies.

The concept of atrial myopathy (AM) or atrial cardiomyopathy (AtCM), first mentioned in 1972, serves as a substrate for the development of AF^8^. AM encompasses structural, chamber architectural, inflammatory, autonomic, and electrophysiological abnormalities that can produce clinically relevant manifestations of AF, impaired atrial function, and thromboembolism^9–12^. Importantly, AM and AF are interlinked but distinct entities: an abnormal atrial substrate may precede AF, whereas rapid and irregular atrial activation can accelerate electrical, contractile, and structural remodelling. Fibrosis is a central component of this process since the associated accumulation in extracellular matrix (ECM) separates cardiomyocyte (CM) bundles, increases conduction heterogeneity, and stabilizes re-entry, while inflammatory and neurohumoral signals promote fibroblast (FB) activation^7,13^. Autoimmune mechanisms may also contribute directly to AF, as demonstrated by the identification of arrhythmogenic autoantibodies against the atrial K_ir_3.4 channel^11^. Left atrial volume index (LAVI) provides a clinically accessible measure of atrial enlargement and structural remodelling associated with AM and is associated with AF recurrence, but atrial volume alone is not necessarily a sufficient proxy for the altered cellular states and electrophysiological changes that underlie disease^9,14^. Distinguishing rhythm-associated alterations from those related to overt structural remodelling is therefore key to providing a deeper understanding of the unique and overlapping mechanisms underlying AF and AM in humans.

Single-cell and single-nucleus technologies have resolved the cellular complexity of the human heart^15–18^. Atlases describing the 4 cardiac chambers have identified region-specific CM, stromal, vascular and immune populations and demonstrated that cardiac cell identity is determined by anatomical context^19^. In AF, single-nucleus transcriptomic (snRNA-seq) studies have implicated CM stress programmes and macrophage activation, whereas cell-type-resolved chromatin maps have localized much of the common AF genetic risk to CM regulatory elements^20–24^. Multiomic analysis of the left atrial appendage (LAA) further identified androgen-receptor-associated CM programmes and *NR4A1*-dependent FB activation^25^. While these studies establish the value of cell-resolved analysis of the atria, their focus has been restricted to defining the transcriptomic changes in AF versus sinus rhythm (SR) within a single chamber. However, the relationships between rhythm status, atrial enlargement, chamber identity, and the transcriptional and regulatory states of different atrial cell populations are not known, but their identification is key for intervening in AF development.

Gene expression changes during differentiation and disease are controlled by alterations in the accessibility and three-dimensional organisation of cell-type-specific *cis*-regulatory elements (CREs) located at promoters, enhancers and insulators^26^. Paired or integrated profiling of RNA and chromatin accessibility can link altered cellular states to candidate CREs, transcription factor (TF) activity and enhancer–gene relationships that cannot be recovered from transcriptomics alone. In the adult human heart, integration of single-nucleus assay for transposase-accessible chromatin using sequencing (snATAC-seq) with snRNA-seq has mapped cell-type- and chamber-specific CREs^27^. Subsequent multiomic profiling studies of human failing hearts have linked changes in chromatin accessibility with disease-associated FB states and immune-FB signalling programmes remodelling^28^. More recently, multimodal profiling across all four cardiac chambers integrating transcriptomes, chromatin accessibility, histone modifications and three-dimensional chromatin organisation revealed extensive regulatory reprogramming and altered enhancer–gene interactions in HF, particularly in CM and FB^29^. However, the *cis*-regulatory programmes coordinating CM stress, FB activation and immune–stromal communication in human atrial remodelling remain largely unknown. Moreover, the contribution of epigenetic regulators to these gene programmes in humans is poorly characterized.

Here, we combined snRNA-seq profiling of human LAA and right atrial appendage (RAA) with paired RNA and chromatin-accessibility profiling of left atrial samples, histological assessment, and functional validation in primary human cardiac FB. To distinguish rhythm-associated from structure-associated remodelling, left atrial samples were organized along crossed AF/SR and high/low-LAVI dimensions, generating Hi-AF, Hi-SR, Lo-AF and Lo-SR groups. This design enabled us to examine chamber-dependent cellular changes, map CM and FB states to candidate *cis*-regulatory networks, and to investigate regulatory candidates within a TGF-β-responsive FB programme. The resulting atlas identifies coordinated CM, FB and macrophage programmes and nominates *JMJD1C* as an epigenetic regulator of human atrial FB activation.

## Methods

An expanded Methods section is available in the Supplementary Materials.

### Study design, participants and atrial tissue collection

Human LAA and RAA specimens were collected from human patients during cardiac surgery (75-250 mg per sample; Department of Cardiology, University Hospital Antwerp, Antwerp, Belgium). Written informed consent was obtained before tissue collection. The study was approved by the institutional review committee of the University Hospital Antwerp, and the procedures followed were in accordance with institutional guidelines (EDGE001634). Samples used for snRNA-seq were snap-frozen in liquid nitrogen and stored at −80 °C, and samples used for histology were fixed in paraformaldehyde (PFA) and stored at 4 °C, respectively. 14 participants contributed both LAA and RAA sequencing specimens; therefore, chamber-specific *N* values denote unique chamber specimens.

For histology and regulome analyses, LAA participants were classified by cardiac rhythm as AF or SR from the clinical record. Structural remodelling was stratified using LAVI, with a threshold of 30 mL/m^2^. This yielded four descriptive groups: high-LAVI AF (Hi-AF), high-LAVI SR (Hi-SR), low-LAVI AF (Lo-AF) and low-LAVI SR (Lo-SR). The groups were treated as the cross-classification of rhythm and structural remodelling. Lo-SR served as the low-remodelling, non-AF reference state where appropriate; Hi-SR represented remodelling without documented AF, Lo-AF showed AF without high LAVI, and Hi-AF the joint phenotype. The integrated transcriptomic analysis comprised 24 LAA chamber samples and 36 RAA chamber samples; 20 LAA samples were included in the paired Multiome regulome analysis. The study protocol conformed to the Declaration of Helsinki and was conducted according to national and European Union regulations on the use of human tissues. Patient information is provided in Supplementary Table 1.

### Single-nucleus Multiome profiling

#### Nuclei isolation and fluorescence-activated cell sorting (FACS)

Nuclei were isolated from heart tissue by mechanical disruption, followed by purification from debris by FACS as previously described^30^. In brief, frozen atrial tissue (4 samples in each pool; approximately 30 mg per sample) was processed on a gentleMACS dissociator in 2 mL ice-cold nuclei-isolation buffer (5 mM CaCl_2_, 2 mM EDTA, 1× protease inhibitor, 3 mM MgAc, 10 mM Tris-HCl (pH 8.0), 0.6 mM DTT and 1 U/μL RNaseOUT). Homogenates were passed through a 30 μM filter and washed with 3 mL nuclei-isolation buffer. Nuclei were pelleted at 500 × *g* for 5 min at 4 °C, resuspended in 500 μL permeabilization buffer for 5 min on ice, diluted with 600-1,000 μL sorting buffer, and stained with DAPI (1:1,000). Approximately 200,000 intact nuclei were sorted using a Sony MA900 cell sorter into 50 μL collection buffer. Sorted nuclei were pelleted at 1,000 × *g* for 15 min at 4 °C, resuspended in 18–25 μL reaction buffer, and counted using a LUNA automated cell counter.

#### Library preparation and sequencing

Paired gene-expression and chromatin-accessibility libraries were prepared from the same nuclei using the 10x Genomics Chromium Single Cell Multiome ATAC + Gene Expression workflow (Single Cell Multiome ATAC + Gene Expression v1), following the manufacturer’s instructions. Approximately 20,000 nuclei were loaded for tagmentation where material permitted. Library concentration and fragment-size distributions were evaluated before pooling. Libraries were sequenced on an Illumina NovaSeq 6000 S4 flow cell with 100-bp paired-end reads. Target depths were approximately 30,000 read pairs per nucleus for gene expression and 50,000 read pairs per nucleus for ATAC.

### snRNA-seq analysis

#### Normalization, integration, and clustering

Raw counts were stored in an AnnData layer, normalized to a total of 10,000 counts per nucleus, and log-transformed as ln(1 + normalized counts)^19,31^. Highly variable genes were selected with a minimum mean of 0.0125, maximum mean of 3, and minimum dispersion of 0.5 while accounting for batch. A composite batch factor combined dataset identifier, donor, and 10x library. Batch integration was performed with single-cell variational inference (scVI) using scvi-tools (v1.1.5)^32^. The model used 128 hidden units, a 50-dimensional latent space, three neural-network layers, and gene-by-batch dispersion; total counts, mitochondrial fraction, and ribosomal fraction were supplied as continuous covariates. The model was trained using mini-batches of 256 nuclei, with default optimization and training-stopping parameters implemented in scVI^19^.

A k-nearest-neighbour (KNN) graph was constructed in the 50-dimensional scVI latent space using Scanpy defaults^33^. UMAP coordinates were computed with min_dist = 0.2. Leiden clustering was first evaluated at resolution 1 and, after removal of low-quality or unclassified nuclei, repeated at resolution 2. Major cell classes were assigned using canonical marker genes. Markers included *MYL7* and *MYL4* for aCM; *DCN* and *PDGFRA* for FB; *VWF*, *PECAM1* and *CDH5* for endothelial cells; *RGS5*, *ABCC9* and *KCNJ8* for mural cells; *MYH11*, *TAGLN* and *ACTA2* for smooth-muscle cells; *MSLN*, *WT1* and *BNC1* for mesothelial cells; *PLP1* and neurexin-family genes for neuronal cells; *GPAM*, *FASN* and *LEP* for adipocytes; *CD14*, *C1QA* and *CD68* for myeloid cells; *CD8A*, *IL7R* and *CD40LG* for lymphoid cells; *CD19*, *SDC1* and immunoglobulin genes for B-lineage cells; and *KIT* and *CPA3* for mast cells.

#### Neighbourhood differential abundance

Differential abundance (DA) (between AF and SR) was tested with Milo as implemented in pertpy (v1.0.3)^34^. Separate LAA and RAA graphs were built from X_scVI with k = 50. Representative overlapping neighbourhoods were sampled with prop = 0.10 and nuclei were counted per donor. AF and SR were encoded with SR as the reference level, and the design was ∼ donor type. Mixed-cell-type neighbourhoods were removed, and cell classes represented by fewer than three neighbourhoods were not interpreted. Milo’s graph-aware spatial false-discovery-rate (spatial FDR) correction was used; neighbourhoods were intended to be considered significant at spatial FDR < 0.05.

#### Comparison with a human heart reference atlas

LAA atlas was compared with the left atrial (LA) atlas from the referenced human heart cell atlas (HCA-LA)^19^ after exclusion of dissociated cells, lymphatic endothelial cells, and vCMs. Atlas nuclei were normalized with Seurat (v5.0.1)^35^ LogNormalize (scale factor 10,000), 2,000 variable features were selected by the variance-stabilizing transformation method, and expression was scaled. In-house LAA data and atlas HCA-LA data were merged and integrated with the SCT workflow using 3,000 integration features and the first 50 principal components. Average expression was calculated for matched major cell types in the integrated assay, and all-gene Pearson correlations were displayed as an unclustered heatmap. Donor-level cell-type proportions were calculated independently in each atlas before comparison.

### Single-nucleus chromatin-accessibility analysis

#### ArchR processing, peak calling and peak matrix construction

ATAC fragments were analysed with ArchR (v1.0.3)^36^ in R 4.1.3. Arrow files were created with minTSS = 1.5 and minFrags = 300 while adding tile and gene-score matrices. Cell identities were transferred from the paired RNA modality. Pseudobulk coverages were generated by cell type using a minimum and maximum of 3 replicates and 40–500 nuclei per group. Reproducible peaks were called from grouped coverages with MACS2 (v2.2.9.1)^37^ using a reproducibility criterion of 2 and eight threads, and the resulting unified peak set was quantified to form the peak matrix.

#### Motifs, chromVAR deviations and footprinting

TF motif annotations were obtained from JASPAR2020 and were added to the chromatin accessibility assay using Signac(v1.8.0)^38^. TF-associated accessibility deviations were calculated using chromVAR (v1.16.0)^39^. Motif footprint profiles were computed by cell type with a smoothing window of five bases and displayed with subtractive, divisive, or unnormalised correction for Tn5 sequence bias, as indicated. Motif enrichment and deviation statistics were interpreted together with cognate TF expression.

### FB state and gene regulatory network (GRN) analyses

#### FB subclustering and trajectory inference

FBs were reanalysed in the scVI representation. A principal-graph trajectory was inferred with scFates (v1.1.1) using SimplePPT in the Palantir embedding^40,41^. The final branch used 60 principal points, seed 5, *λ* = 100, *σ* = 0.025, and 200 optimization steps on CPU. The root principal point was selected from FB_basal responsibilities (root node 29), and pseudotime was computed with one mapping, seed 40, and one process. Genes associated with pseudotime were tested with the scFates association framework.

#### SCENIC+ enhancer-driven regulatory networks (eRegulons)

Enhancer-driven regulatory networks (eRegulons) were inferred with SCENIC+(v1.0.2)^42^. For the accessibility input, MACS2 (v2.2.9.1)^43^ peaks were called on BEDPE fragments for the human genome with shift = 73, extension size = 146, duplicate retention set to all, *q* = 0.05, and 10 CPUs. Fragment-level doublets were scored with Scrublet using expected_doublet_rate = 0.10 and removed at a score threshold of 0.22. Candidate topic models were evaluated across 2, 5, 10, 15, 20, 25, 30, 35, 40, 45, and 50 topics with 500 iterations, seed 555, *α* = 50, topic-specific *α* enabled, and *η* = 0.1. SCENIC+ combined pycisTopic accessibility topics, motif enrichment, and paired expression to infer direct and extended eRegulons, region- and gene-level AUCell scores, regulon specificity scores, and TF–region–gene networks. eRegulon activity scores were represented by AUCell based on target gene expression and target region accessibility.

#### Cell–cell communication analysis

Ligand–receptor communication was assessed with CellPhoneDB (v5.0.1)^44,45^ and database v5.0.0. Separate analyses were run for the LAVI/rhythm groups. Statistical analysis used 1,000 label permutations, a 10% minimum expression threshold, five threads, random seed 42, three-digit output precision, and interaction scoring enabled; subsampling was disabled, and the nominal interaction threshold was *P* < 0.05.

### Primary human cardiac fibroblast (HCF) experiments

#### Cell culture, TGF-β stimulation and siRNA transfection

Primary human cardiac fibroblasts (Innoprot, P10454) were used at passages 4–6 and maintained in fibroblast medium (Innoprot, P60108) supplemented with 10% foetal bovine serum (FBS), 10% fibroblast growth supplement and 1% penicillin–streptomycin. Cells were exposed to recombinant TGF-β1 (50 ng/mL; PeproTech, 100-21-10UG) for 48 h. For JMJD1C knockdown, cells were transfected with a non-targeting control or JMJD1C-targeting siRNA (Thermo Fisher Scientific, 10446914) using Lipofectamine RNAiMAX (Invitrogen, 13778075) according to the manufacturer’s protocol. Specifically, 9.0 μL of RNAiMAX reagent and 3.0 μL of 10 μM siRNA were added per well of a 6-well plate (final concentration 25 pmol). After 48 h, media was exchanged and cells treated with TGF-β1 or vehicle for a further 48 h, as indicated.

### Bulk RNA sequencing of fibroblast perturbations

#### Library preparation and sequencing

Bulk RNA-seq was performed on five biological replicates for siRNA control and JMJD1C knockdown, each with and without TGF-β1 treatment. RNA was purified using the RNeasy Plus Mini Kit (Qiagen, Germany) according to the manufacturer’s protocol, and genomic DNA was removed using gDNA Eliminator columns. 500 ng RNA was submitted for library preparation. Poly(A)-selected libraries with UMIs were prepared and sequenced by Azenta (Germany) on an Illumina NovaSeq 6000 with 150-bp paired-end reads.

### Statistical analysis and reproducibility

Statistical tests were two-sided unless explicitly stated. For genome-wide analyses, multiple testing was controlled with the method native to the analysis: Bonferroni correction for Seurat/MAST outputs, spatial FDR for Milo, and Benjamini–Hochberg false-discovery-rate correction for DESeq2^46^, clusterProfiler^47^, gene-set enrichment^48^ and limma^49^ outputs. Exact thresholds and the unit of replication are given in the relevant subsection and figure legend. For cell culture and RT–qPCR experiments, comparisons between two experimental conditions were performed using two-tailed paired t-tests, whereas comparisons involving more than two groups were performed using one-way ANOVA followed by Tukey’s multiple-comparison test, as appropriate. For histological analyses, data points represent independent patient specimens and statistical significance was assessed using one-way ANOVA followed by Tukey’s multiple-comparison test.

### Data and code availability

Differential-analysis tables and analysis notebooks will be deposited in GitHub. Raw Multiome and RNA-seq sequencing data will be deposited in a controlled-access archive (GEO), and non-identifying processed objects, figure-generating code, environment files and parameterised workflows will be deposited in GitHub.

## Results

### A single-nucleus atlas resolves chamber-specific cellular remodelling in AF

To define cellular remodelling across the human atria, we generated snRNA profiles from LAA and RAA specimens, combining dedicated snRNA-seq libraries with the RNA component of paired Multiome libraries (**Fig. 1a)**. After quality control (QC) and donor demultiplexing, the atlas comprised 25,119 LAA nuclei from 24 chamber samples and 50,535 RAA nuclei from 36 chamber samples (Supplementary Table 1). Unsupervised clustering resolved atrial CM (aCM), FB, endothelial, mural, mesothelial, myeloid, lymphoid, adipocyte, mast-cell and neuronal populations in both chambers (**Fig. 1b,f**), and canonical marker expression supported these annotations (**Fig. 1c,g**). QC embeddings showed that retained nuclei were distributed across donors, rhythm groups, sexes, and 10x library types without a single technical variable defining the major clusters (Supplementary Figs. 1 and 2).

**Fig. 1.**
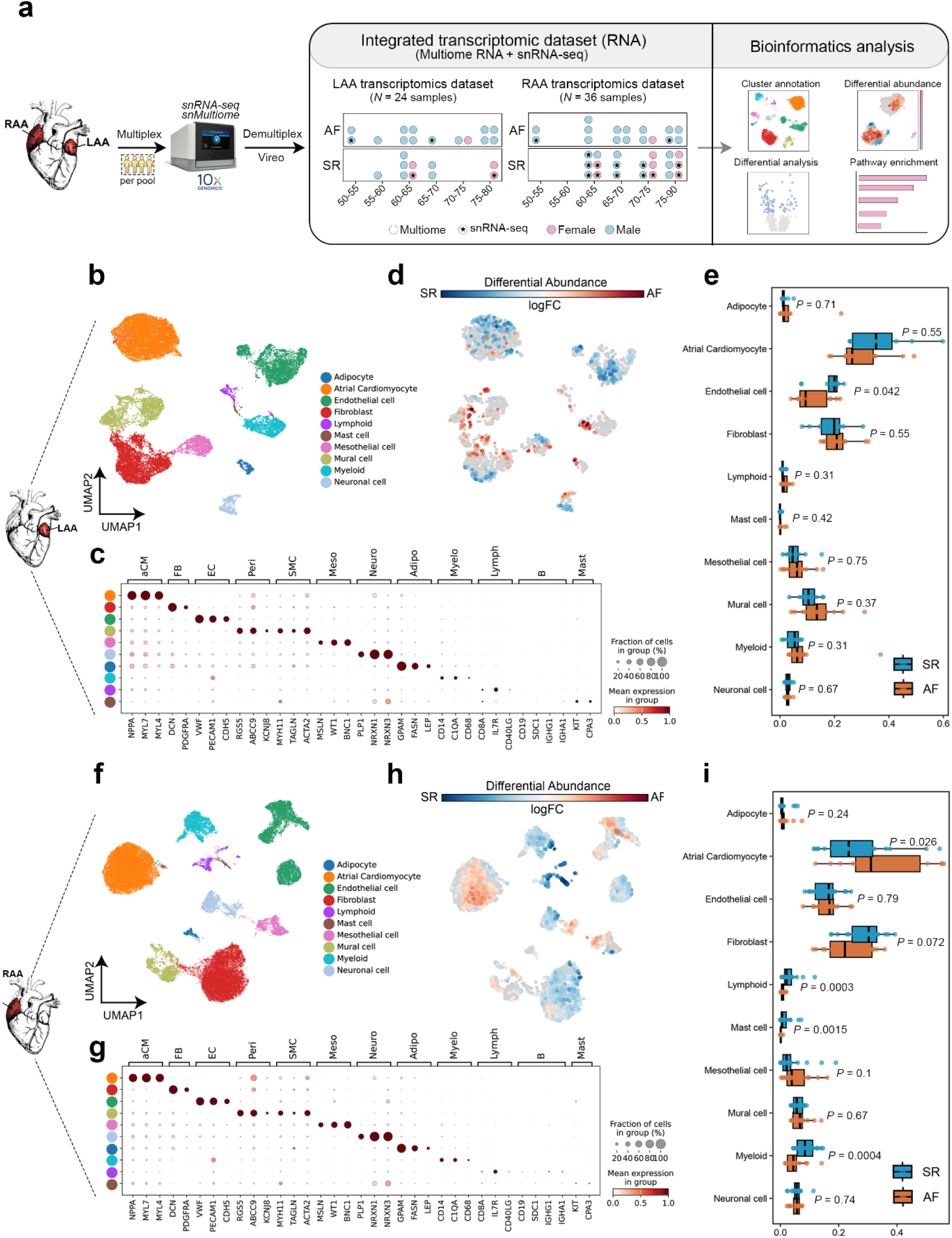
Single-nucleus transcriptomic atlases of the human left and right atrial appendages. **a**, Study overview. Left atrial appendage (LAA) and right atrial appendage (RAA) specimens were collected from donors stratified by rhythm (atrial fibrillation (AF) and sinus rhythm (SR)), sex, and age; nuclei were multiplexed before library preparation and assigned to donors with Vireo and integrated with the single-nucleus RNA (snRNA) component of the 10x Multiome dataset. Downstream analyses included cell type annotation, differential abundance (DA), differential expression (DE), and pathway enrichment. **b**, UMAP of 25,119 LAA nuclei from 24 samples, coloured by major cell identity. **c**, Canonical marker expression used to annotate major cell populations in the LAA; colour denotes scaled mean expression and point size denotes the fraction of expressing nuclei. **d**, Milo neighbourhood DA map comparing LAA samples from AF and SR donors. Red and blue neighbourhoods are relatively enriched in AF and SR, respectively; nonsignificant neighbourhoods (false discovery rate (FDR) > 5%) are grey. **e**, Donor-level cell-type proportions in AF and SR in the LAA; each point represents one donor and boxes summarize the distributions as drawn. **f**, UMAP of 50,535 RAA nuclei from 36 samples. **g**, Canonical marker expression used to annotate major cell populations in the RAA. **h**, Corresponding Milo neighbourhood map for RAA. **i**, Donor-level cell-type proportions in AF and SR in the RAA. Statistical significance was assessed using a two-sided Wilcoxon rank-sum test for each cell type.

We next asked whether AF was associated with chamber-specific changes in cellular abundance. Milo differential-abundance (DA) analysis tested overlapping neighbourhoods of transcriptionally similar nuclei defined on the K-nearest-neighbour (KNN) graph. Both AF- and SR-enriched neighbourhoods were detected across several cellular compartments in the LAA and RAA, indicating that the direction of AF-associated changes varied within major cell types (**Fig. 1d,h**). Donor-level analyses supported changes in several compartments, for example, a modest increase in the FB proportion in AF in LAA (**Fig. 1e**), but showed broad overlap between rhythm groups and substantial inter-individual variability (**Fig. 1e,i**; Supplementary Figs. 3 and 4). Thus, AF-associated abundance changes were heterogeneous and were often enriched within transcriptional neighbourhoods rather than uniformly affecting an entire major cell type.

To determine whether appendage tissue captured the principal cellular features of the corresponding atrial chamber, we compared our LAA and RAA atlases with left-atrial (HCA-LA) and right-atrial (HCA-RA) free-wall populations, respectively, from the Human Cardiac Cell Atlas^19^. LAA and RAA populations were transcriptionally similar to their respective HCA reference populations (Supplementary Fig. 5a,b,g,h), and matched cell-type proportions were correlated (Supplementary Fig. 5c–f,i–l). These relationships were observed both in the complete in-house sample collections, comprising AF and SR samples (Supplementary Fig. 5c,e,i,k), and in the analyses restricted to SR samples (Supplementary Fig. 5d,f,j,l), thereby reducing the potential influence of AF-associated remodelling. These findings support the use of LAA and RAA tissue to investigate the major cell populations of the left and right atria.

We then compared the overall transcriptomic differences between the two appendage datasets. The integrated embedding of the two datasets preserved the major cell identities (**Fig. 2a**) and showed mixing across rhythm, sex and library type while retaining chamber-associated structure (**Fig. 2b**). Bulk (all cell types combined) nuclei-level differential-expression (DE) analysis identified 761 genes with higher expression in LAA and 1,892 with higher expression in RAA (**Fig. 2c**). Genes elevated in LAA were enriched for heart-development, cell matrix adhesion, FB proliferation, and K^+^-transport processes (**Fig. 2d**). Despite these molecular differences, the aggregate cell-type compositions of LAA and RAA were closely correlated when AF and SR samples were analysed together (Spearman’s *R* = 0.964, *P* < 0.001) and when the comparison was restricted to SR samples (Spearman’s *R* = 0.903, *P* = 0.001) (**Fig. 2e,f**; Supplementary Fig. 6); matched cell types also showed chamber-dependent transcriptional similarity (**Fig. 2g**), with neuronal cells showing the highest similarity (*r* = 0.75). Sex-stratified embeddings likewise retained the major populations (**Fig. 2h,k**; Supplementary Table 2), and cell-type proportions (**Fig. 2i,l**; Supplementary Fig. 7). Transcriptional patterns were also highly correlated between male and female samples in both LAA and RAA (**Fig. 2j,m**). Together, these analyses establish a chamber-resolved human atrial atlas, revealing broadly conserved cellular architecture alongside distinct chamber-specific patterns of transcriptional remodelling.

**Fig. 2.**
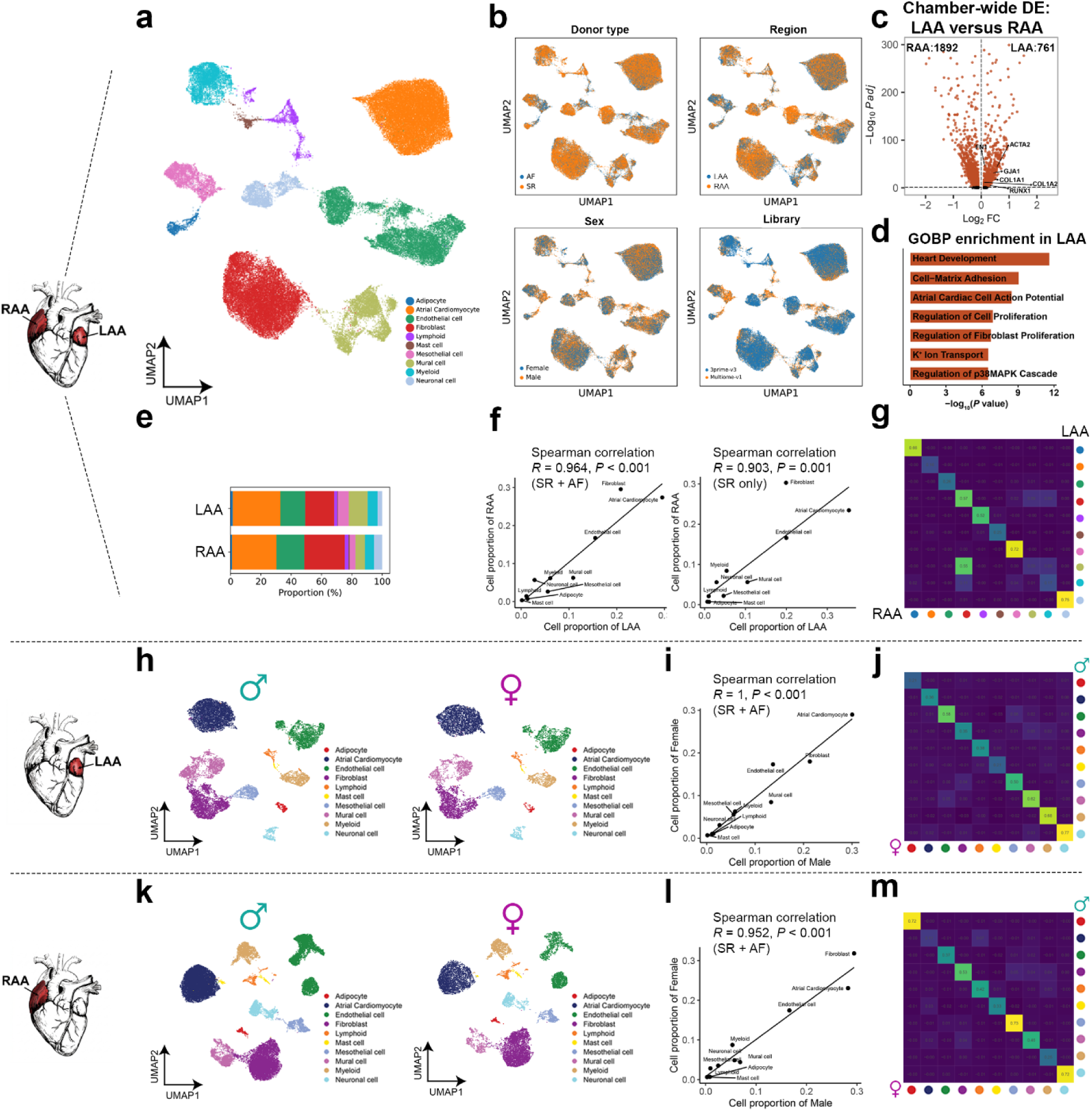
Chamber- and sex-associated features of the human atrial single-nucleus atlas. **a**, UMAP of the integrated LAA and RAA atlas, coloured by major cell identity. The schematic indicates the anatomical sampling sites. **b**, Integrated UMAPs coloured by donor rhythm status (top left), atrial region (top right), sex (bottom left), and 10x library (bottom right), respectively. **c**, Chamber-wide DE analysis comparing LAA with RAA using MAST. Volcano plots show the average log₂FC on the x-axis and −log₁₀ of the Bonferroni-adjusted *P* value on the y-axis. Positive log₂FC indicates higher expression in LAA and negative values indicate higher expression in RAA. Genes with adjusted *P* < 0.05 and |log₂ fold change| ≥ 0.25 are shown in red; selected extracellular matrix (ECM) and contractile genes are labelled. The numbers indicate the differentially expressed genes (DEGs) enriched in each chamber. **d**, Gene Ontology Biological Process (GOBP) enrichment among LAA-associated genes. Enrichr analysis was performed using the 200 genes with positive log₂FC that ranked highest by the MAST-adjusted *P* value. Bars show −log_10_ of the Enrichr enrichment *P* value for the displayed terms. **e**, Aggregate proportions of the major cell populations in the LAA and RAA datasets. **f**, Spearman correlations of aggregate cell-type proportions between LAA and RAA using samples from AF and SR donors together (left) or samples from SR donors only (right). Each point represents one major cell type; correlation coefficients and two-sided *P* values are shown. **g**, Cross-chamber cell-type similarity matrix comparing transcriptional identities between the LAA and RAA atlases. Rows and columns represent major cell types; colour and numerical values indicate the corresponding correlation coefficient. **h**, LAA UMAPs showing nuclei from male (left) and female (right) participants, coloured by major cell identity. **i**, Spearman correlation of aggregate cell-type proportions between male and female LAA samples. Each point represents one major cell type; the correlation coefficient and two-sided *P* value are shown. **j**, Cross-sex cell-type similarity matrix comparing the transcriptional identities of major LAA cell populations between male and female participants. **k**, RAA UMAPs showing nuclei from male (left) and female (right) participants, coloured by major cell identity. **l**, Spearman correlation of aggregate cell-type proportions between male and female RAA samples. Each point represents one major cell type; the correlation coefficient and two-sided *P* value are shown. **m**, Cross-sex cell-type similarity matrix comparing the transcriptional identities of major RAA cell populations between male and female participants. UMAP, uniform manifold approximation and projection; FC, fold change.

### AF is accompanied by chamber-dependent transcriptional and structural remodelling

Having established the broad chamber architecture, we next examined AF-associated transcriptional programmes. Cell-type-resolved analysis identified significant AF-associated DE across the major LAA and RAA populations (**Fig. 3a,h**; Supplementary Fig. 8; Supplementary Tables 3 and 4). Across evaluable LAA cell types, 2–375 genes were more highly expressed, and 2–341 were less expressed in AF; the corresponding ranges in RAA were 11–622 and 3–1,003 genes, respectively. Chamber-wide analysis identified 986 genes with higher expression and 1,051 lower in AF in LAA, compared with 1,224 higher and 1,926 lower in AF in RAA (**Fig. 3b,i**). In LAA, AF-associated genes included ECM and remodelling factors such as *COL1A1*, *COL1A2*, *COL3A1*, *FN1*, *AEBP1*, *RUNX1* and *LTBP2*, and enriched processes included sarcomere organisation, cardiac contraction, ECM organisation, TGF-β signalling and immune regulation (**Fig. 3b,c**). Cell-type-specific analyses further resolved altered contractile, Ca^2+^ handling, ion-transport and action potential (AP) programmes in aCMs (**Fig. 3d,e**), together with PDGF-, IL-1- and matrix-associated programmes in FBs (**Fig. 3f,g**). RAA showed widespread AF-associated changes dominated by contractile, sarcomeric, Ca^2+^-transport and inflammatory programmes, with comparatively less prominent ECM enrichment than LAA (**Fig. 3i–n**). Thus, AF was accompanied by extensive but chamber-dependent transcriptional remodelling.

**Fig. 3.**
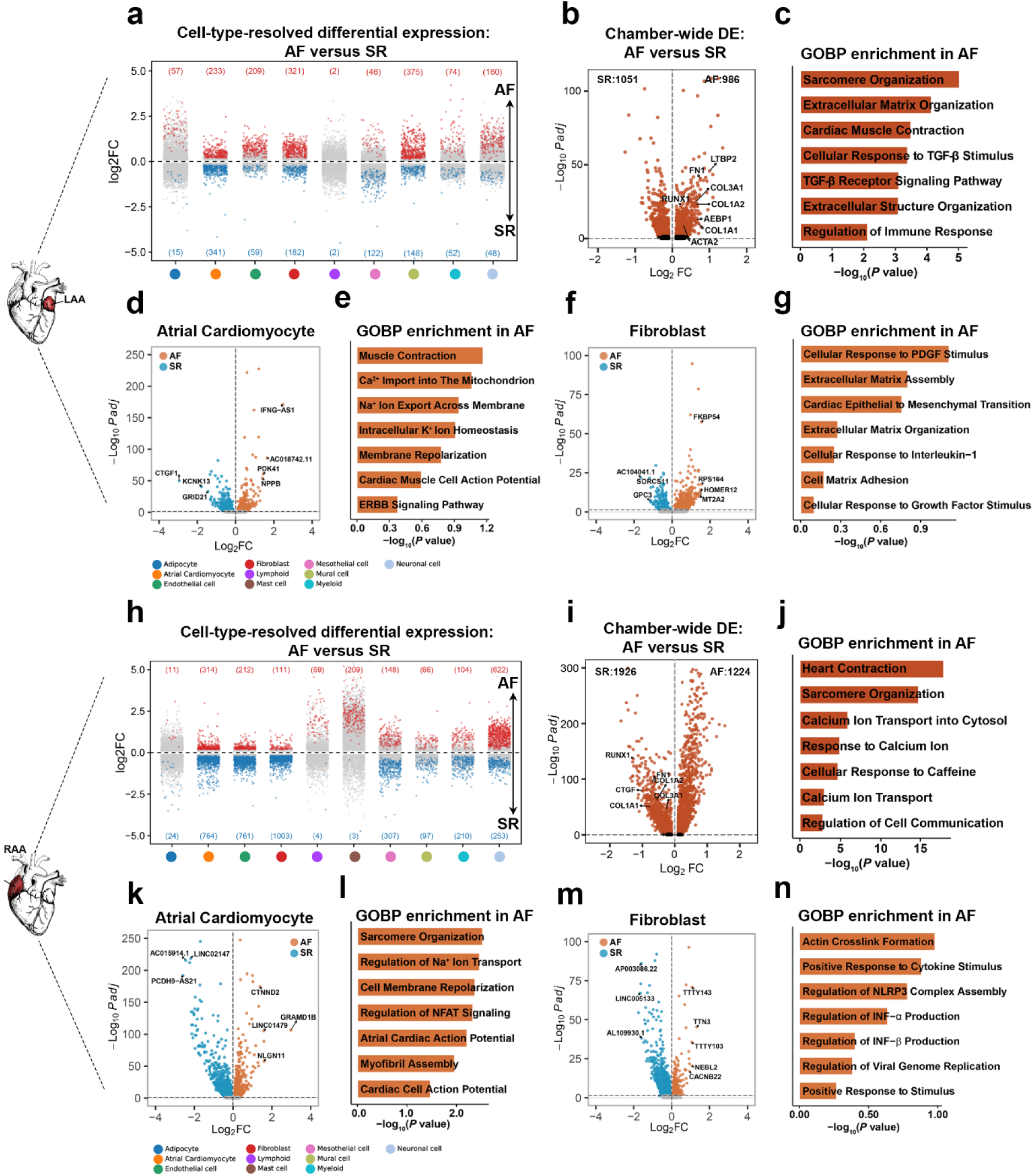
AF-associated transcriptional remodelling across atrial chambers and cell types. **a**, Cell-type-resolved DE analysis comparing AF with SR in the LAA. DE was evaluated separately within each major cell type using the MAST model. Genes were tested when detected in at least 10% of nuclei in either group and when the absolute average log₂FC was at least 0.10. Each point represents one tested gene. Genes with Bonferroni-adjusted *P* < 0.05 and absolute average log₂FC > 0.20 are shown in red when increased in AF and blue when increased in SR; other tested genes are shown in grey. Numbers indicate the corresponding numbers of highlighted genes. **b**, Chamber-wide DE analysis comparing all LAA nuclei between AF and SR using MAST. The volcano plot shows average log₂FC and −log₁₀ of the Bonferroni-adjusted *P* value. Coloured genes had adjusted *P* < 0.05; no additional effect-size threshold was applied for colouring beyond the gene-testing threshold of absolute average log₂FC ≥ 0.10. Selected ECM and remodelling-associated genes are labelled. **c**, GOBP enrichment among genes with higher expression in AF in the chamber-wide LAA analysis. **d,** Volcano plots comparing AF with SR within LAA aCM. Orange and blue denote genes with higher expression in AF and SR, respectively; selected top genes are labelled. **e,** GOBP enrichment among genes with higher expression in AF in LAA aCMs. **f**, Volcano plots comparing AF with SR within LAA FB. Orange and blue denote genes with higher expression in AF and SR, respectively; selected top genes are labelled. **g**, GOBP enrichment among genes with higher expression in AF in LAA FBs. **h**, Cell-type-resolved DE analysis comparing AF with SR in the RAA. **i**, Chamber-wide DE analysis comparing all RAA nuclei between AF and SR. **j**, GOBP enrichment among genes with higher expression in AF in the chamber-wide RAA analysis. **k,** Volcano plots comparing AF with SR within RAA aCM. Orange and blue denote genes with higher expression in AF and SR, respectively; selected top genes are labelled. **l,** GOBP enrichment among genes with higher expression in AF in RAA aCMs. **m**, Volcano plots comparing AF with SR within RAA FB. Orange and blue denote genes with higher expression in AF and SR, respectively; selected top genes are labelled. **n**, GOBP enrichment among genes with higher expression in AF in RAA FBs. For all enrichment analyses, Enrichr was applied to the 200 genes with positive log₂FC that ranked highest by the MAST-adjusted *P* value. Bars show −log₁₀ of the adjusted enrichment *P* value for the displayed terms. Positive log₂FC indicates higher expression in AF, whereas negative log₂FC indicates higher expression in SR. Schematics indicate the atrial appendage analysed in each section. DE, differential expression; FC, fold change; GOBP, Gene Ontology biological process. aCMs, atrial cardiomyocytes; FBs, fibroblasts.

Since atrial enlargement is considered a prequel to AF, and frequently coexists, to examine their relationship and whether atrial enlargement represented a predisposing step to AF, we next separated LAA samples along two crossed clinical dimensions: rhythm (AF or SR) and structural-remodelling burden indexed by LAVI (high or low; Hi-AF, Hi-SR, Lo-AF and Lo-SR) and compared transcriptional changes between them. Pairwise comparisons were organised around Lo-SR as the low-remodelling reference and around contrasts that isolated rhythm or LAVI within a stratum. Hi-AF versus Lo-SR showed the broadest combined phenotype, with enrichment of heart-contraction, TGF-β-receptor, extracellular-structure and immune-response programmes (**Fig. 4a,b**). By comparison, Lo-AF versus Lo-SR predominantly highlighted contractile, fatty-acid-oxidation, ion-homeostasis, stretch-response and MAPK programmes, with little evidence of a strong fibrotic signature (**Fig. 4d,e**). Comparing Hi-AF with Lo-AF, thereby holding AF status constant, identified TGF-β production and receptor signalling, collagen-fibril and ECM organisation, and IL-1-production programmes associated with high LAVI (**Fig. 4g,h**). Within SR, Hi-SR relative to Lo-SR showed enrichment of myofibril assembly, cardiac contraction, Ca^2+^ responses and ERK1/2 signalling (Supplementary Fig. 9a,b). Conversely, comparing Hi-AF with Hi-SR while holding high LAVI constant identified ECM, collagen-fibril and supramolecular-fibre organisation, TGF-β receptor signalling and histone lysine demethylation in Hi-AF (Supplementary Fig. 9d,e). Cell-type similarity matrices and donor-level composition correlations remained high across each comparison, indicating that these pathway differences occurred largely within a conserved major-cell-class architecture rather than through wholesale replacement of cell types (**Fig. 4a,c,d,f,g,i**; Supplementary Fig. 9c,f).

**Fig. 4.**
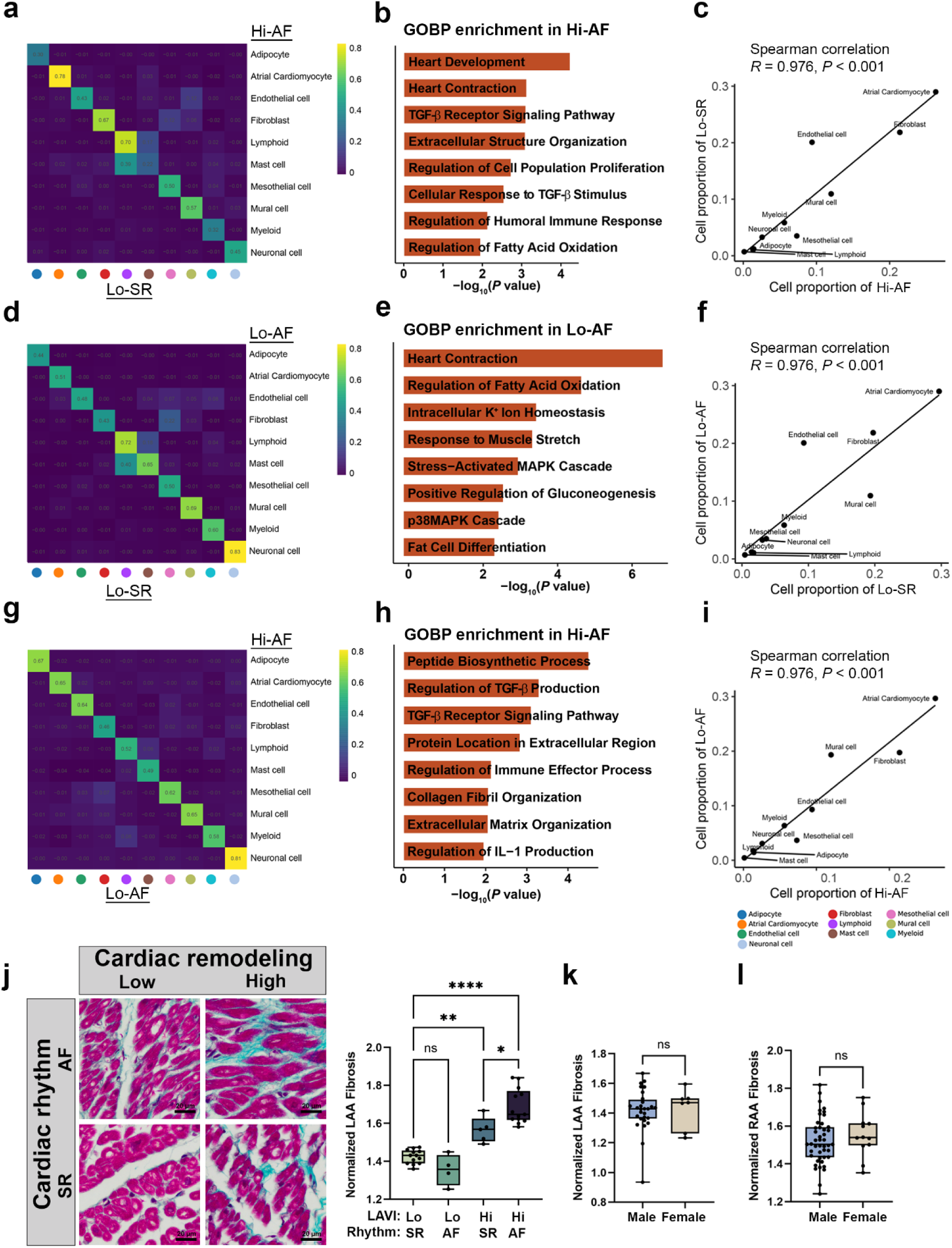
Pairwise rhythm–LAVI comparisons distinguish AF- and structural-remodelling-associated atrial signatures. **a**, Cell-type similarity matrix comparing LAA populations from participants with AF and high left atrial volume index (Hi-AF) with those from participants in SR with low LAVI (Lo-SR). Rows and columns represent major cell types; colour and numerical values indicate the corresponding correlation coefficient. **b**, Selected GOBP terms enriched among genes with higher expression in Hi-AF than in Lo-SR. Bar length represents −log₁₀(*P* value). **c**, Spearman correlation of aggregate cell-type proportions between the Hi-AF and Lo-SR groups. Each point represents one major cell type; the correlation coefficient and two-sided *P* value are shown. **d**, Cell-type similarity matrix comparing Lo-AF with Lo-SR. **e**, Selected GOBP terms enriched among genes with higher expression in Lo-AF than in Lo-SR. **f**, Spearman correlation of aggregate cell-type proportions between Lo-AF and Lo-SR. **g**, Cell-type similarity matrix comparing Hi-AF with Lo-AF. **h**, Selected GOBP terms enriched among genes with higher expression in Hi-AF than in Lo-AF. **i**, Spearman correlation of aggregate cell-type proportions between Hi-AF and Lo-AF. **j**, Left, representative Masson’s trichrome-stained LAA sections from Hi-AF, Hi-SR, Lo-AF and Lo-SR samples. Collagen-rich fibrotic tissue is stained blue and myocardium red. Scale bar = 20 µm. Right, donor-level quantification of normalized LAA fibrosis across the four rhythm–LAVI groups (Hi-AF, *n* = 13; Hi-SR, *n* = 5; Lo-AF, *n* = 4; and Lo-SR, *n* = 13). Differences were assessed using an ordinary one-way ANOVA followed by Tukey’s two-sided multiple-comparison test. **k**, Normalized histological fibrosis in LAA samples from male and female participants (male, *n* = 28; female, *n* = 7). **l**, Normalized histological fibrosis in RAA samples from male and female participants (male, *n* = 45; female, *n* = 12). In k,l, differences between sexes were assessed using an unpaired two-tailed *t*-test. In j–l, each point represents one biological sample; boxplots show the median and interquartile range, with whiskers extending to the most extreme observations within 1.5 times the interquartile range. ns, not significant; \*\**P* < 0.01; \*\*\*\**P* < 0.0001. Hi-AF, AF with high LAVI; Hi-SR, SR with high LAVI; Lo-AF, AF with low LAVI; Lo-SR, SR with low LAVI.

We next asked whether the matrix-remodelling transcriptomic signature was reflected in tissue-level fibrosis. Masson’s trichrome staining of 35 LAA samples showed group-dependent collagen deposition, with the greatest fibrosis in Hi-AF. Fibrosis was significantly higher in Hi-AF and Hi-SR than in Lo-SR, whereas Lo-AF did not differ significantly from Lo-SR (**Fig. 4j**). Sex-stratified analysis detected no significant difference in normalised fibrosis between male and female samples in either the LAA or RAA (**Fig. 4k,l**), although the unequal sex distribution limited statistical power. The histology therefore independently supported the association between high LAVI and fibrotic remodelling and placed the strongest fibrotic phenotype in samples with both high LAVI and AF.

### Paired Multiome profiling maps the human LAA regulome

The rhythm- and LAVI-associated transcriptional changes prompted us to investigate their candidate regulatory basis. Since matrix- and TGF-β-related signatures were most prominent in LAA, we investigated remodelling of this chamber further and subjected samples to paired snRNA-seq and chromatin-accessibility (ATAC-seq; 10x Multiome) profiling in 20 LAA samples. From this analysis, 12,189 high-quality nuclei were retained (**Fig. 5a**). Barcode-rank, fragment-length, transcription-start-site (TSS) enrichment, fraction-in-peaks and duplication metrics supported library quality (Supplementary Fig. 10a–j), while donor- and group-coloured embeddings showed integration across samples (Supplementary Fig. 10k,l).

**Fig. 5.**
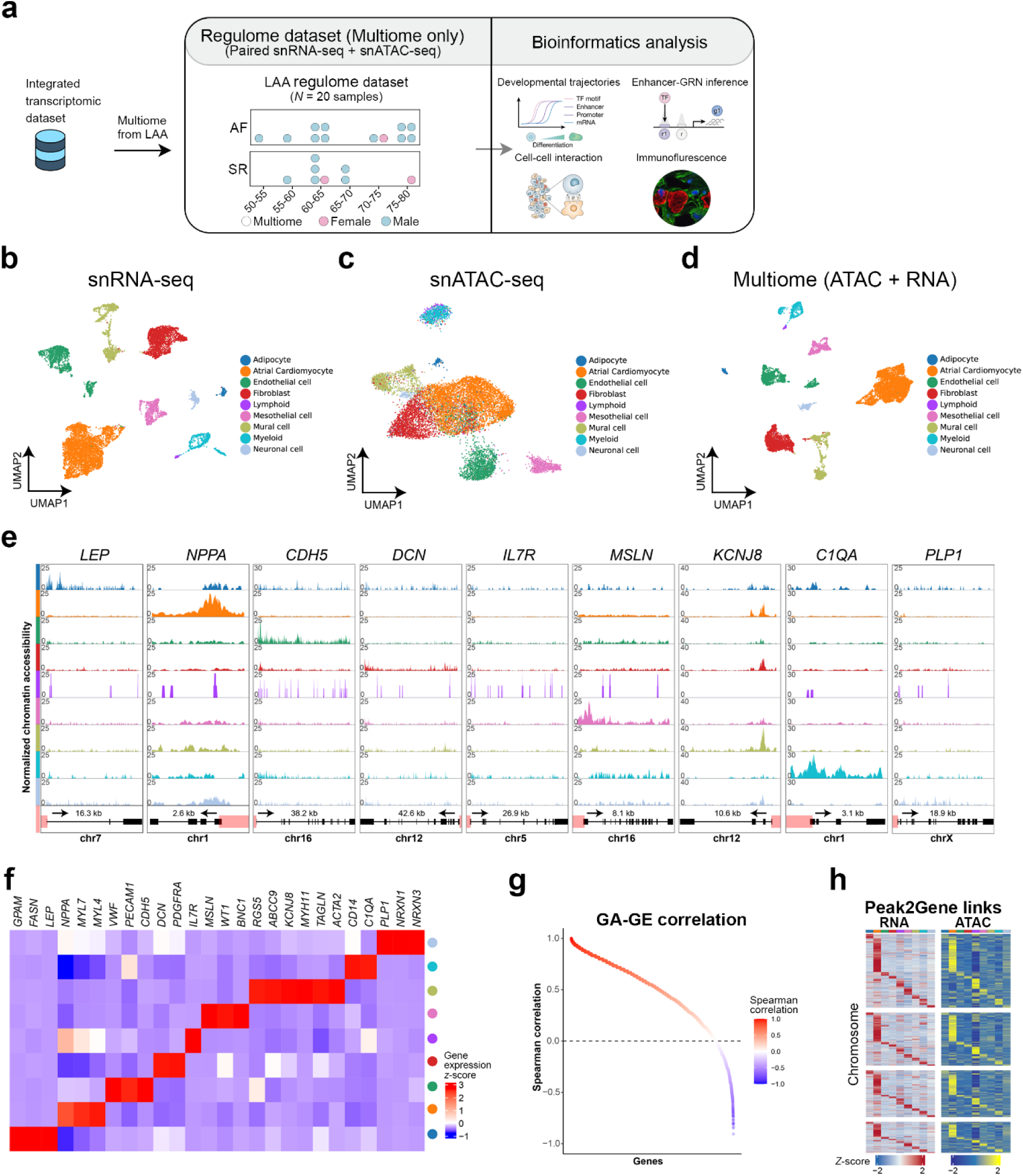
Paired single-nucleus transcriptome and chromatin-accessibility profiling defines the human left atrial regulome. **a**, Workflow for paired snRNA-seq and snATAC-seq profiling of 20 LAA samples by 10x Multiome, followed by modality-specific and integrated analysis. **b**–**d**, UMAPs of the RNA representation (**b**), ATAC representation (**c**), and integrated multiomic representation (**d**), coloured by major cell identity. **e**, Cell-type-specific chromatin-accessibility tracks at representative lineage-marker loci; For each gene, 1 kb upstream and downstream are shown. The promoter/transcription start site (TSS) is highlighted in pink. **f**, Heatmap of row-scaled expression of canonical lineage markers across cell types. **g**, Ranked Spearman correlations between gene activity and gene expression (GA-GE) within the same nucleus across genes. **h**, Chromosome-stratified heatmaps of linked RNA expression and ATAC accessibility for 18,793 peak-to-gene associations, ordered consistently between cell types; Chr1-4 are shown; see the full heatmap in Supplementary Fig. 11. Cell types in e, f, and h are coloured as in a. P2G, Peak2Gene.

RNA, ATAC and joint embeddings independently recovered the major atrial populations, except for the rare mast cell population (**Fig. 5b–d**). The RNA modality provided finer separation of several populations (**Fig. 5b**), whereas the lower-dimensional ATAC representation preserved the broad lineage structure (**Fig. 5c**); their joint embedding associated each epigenomic profile with its paired transcriptome (**Fig. 5d**). Aggregate accessibility at lineage-marker loci corresponded with cell-type-specific RNA expression (**Fig. 5e,f**), for example, *NPPA* in aCM, *DCN* in FB, and *C1QA* in myeloid cells. Cross-modality analysis showed that the dominant cell-identity programmes were concordant, although a minority of genes had high inferred accessibility but low expression (Supplementary Fig. 11a,b). Gene-activity and gene-expression values were positively correlated for most genes (**Fig. 5g**). Peak-to-gene (P2G) analysis identified 18,793 candidate accessibility–expression links distributed across chromosomes and cell types (**Fig. 5h**; Supplementary Fig. 11c). This paired regulome enabled disease-associated transcriptional states to be connected with candidate *cis*-regulatory elements (cCREs) and regulatory factors.

### A stressed CM state exhibits transcriptional and *cis*-regulatory remodelling

Subclustering of Multiome aCM nuclei resolved four states. One state, aCM4, co-expressed the established stress markers *NPPA* and *NPPB* and was therefore annotated as a stressed aCM state^50^ (**Fig. 6a,b**; Supplementary Fig. 12a–d). aCM states differed in Ca^2+^ channel, ion-channel, ryanodine receptor (RYR2) and stress-marker expression, including *KCNN2* and *KCNJ3/KCNJ5*-encoded GIRK-channel components (**Fig. 6c**; Supplementary Fig. 12e), together with their transcriptional-entropy profiles (**Fig. 6d**; Supplementary Fig. 12f). At the donor level, the aCM4 proportion was larger in Hi-AF than in Lo-SR samples (adjusted *P* = 0.0364; **Fig. 6e**), associating this state with the combined AF and high-LAVI phenotype.

**Fig. 6.**
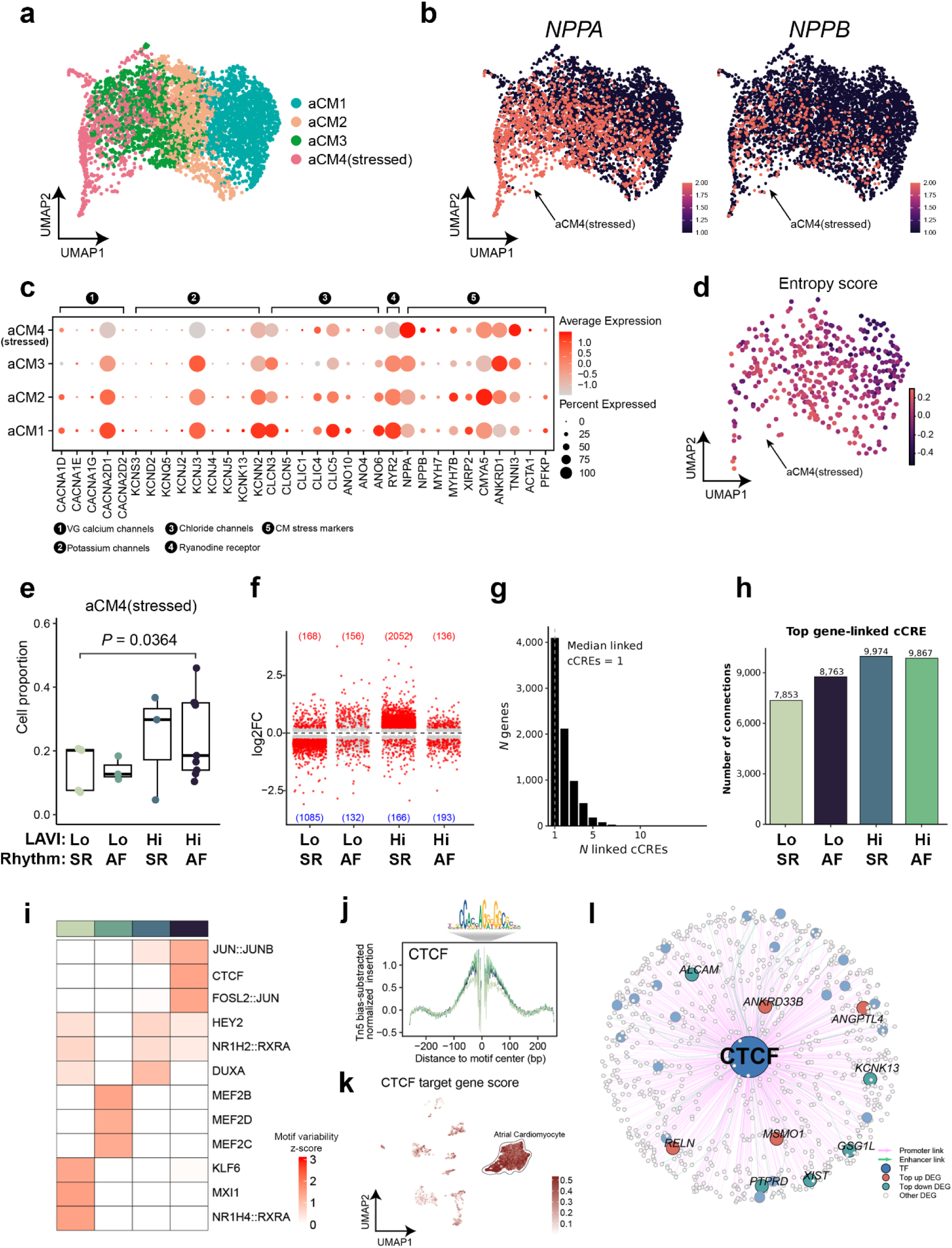
A stressed atrial cardiomyocyte state exhibits altered transcription and *cis*-regulatory elements. **a**, UMAP of atrial cardiomyocytes (aCMs) identifying four states (aCM1–aCM4). **b**, UMAP feature plots of *NPPA* and *NPPB* expression. **c**, Dot plot of selected Ca^2+^ channels, K^+^ channels, Cl^-^channels, RYR receptor, and CM stress genes across states; colour denotes scaled mean expression and point size the fraction of expressing nuclei. **d**, Neighbourhood-level chromatin entropy projected onto the aCM embedding after adjustment for neighbourhood size and latent transcriptional variance. **e**, Donor-level proportion of stressed aCM4 across groups; pairwise comparisons were performed using a two-sided Wilcoxon rank-sum test with Benjamini–Hochberg correction for multiple comparisons; adjusted *P* value is shown for the indicated comparison (Hi-AF: *n* = 9; Hi-SR: *n* = 3; Lo-AF: *n* = 3; Lo-SR: *n* = 5). **f**, DE summaries for the indicated group comparisons; colour denotes significance, and labels give the numbers of significant genes (*P* adj < 0.05 and log_2_FC ≥ 0.25). **g**, Distribution of the number of linked candidate *cis*-regulatory elements(cCREs) per gene (median, one). **h**, Numbers of gene–cCRE connections in Hi-AF, Hi-SR, Lo-AF and Lo-SR. **i**, Motif-variability z-scores for top transcription factors (TFs) across groups. Groups are coloured as in e. **j**, Bias-corrected aggregate *CTCF* footprint centred on the *CTCF* motif; TF binding motif shown as motif logo above. **k**, Integrated UMAP coloured by a *CTCF* target-gene score with aCM circled. **l**, *CTCF*-centred regulatory network linking promoter and enhancer cCREs to DEGs; top -up and -down DEGs labelled. FC, fold change; RYR2, ryanodine receptor 2; DE, differential expression.

Pairwise rhythm–LAVI comparisons identified extensive aCM transcriptional differences, with Hi-SR showing the greatest number of changes relative to Lo-SR among the displayed comparisons (**Fig. 6f**; Supplementary Table 6). cCRE connectivity also differed across groups. Although most linked genes were connected to one cCRE, the total number of gene–cCRE links ranged from 7,853 in Lo-SR to 8,763 in Hi-AF, 9,867 in Lo-AF and 9,974 in Hi-SR, with shared and group-selective linked genes and genomic annotations (**Fig. 6g,h**; Supplementary Fig. 13; Supplementary Table 10). Motif analysis identified group-dependent regulatory variation, including increased CTCF motif variability in Hi-AF (**Fig. 6i**). CTCF is a chromatin architectural protein that contributes to insulator boundaries and constrains enhancer– promoter communication; altered CTCF occupancy can therefore reorganise regulatory contacts^46,47^. Aggregate footprinting showed accessibility around the CTCF motif, and a CTCF target-gene score localised predominantly to aCMs (**Fig. 6j,k**). The inferred network connected promoter and enhancer cCREs with Hi-AF-associated genes, including *KCNK13*, *GSG1L*, *ANKRD33B*, *ALCAM*, *ANGPTL4*, *MSMO1*, *PTPRD*, *XIST* and *RELN* (**Fig. 6l**). *CTCF* expression (Supplementary Fig. 14a,b), motif variability (Supplementary Fig. 14c,d) and target scores (Supplementary Fig. 14e,f) also varied across rhythm–LAVI groups and aCM states (Supplementary Tables 9 and 10). Together, these analyses link the stressed aCM4 state to extensive reorganisation of a cCRE and nominate CTCF as a potential architectural regulator.

### PTGDS^+^ FBs show an inflammatory–fibrotic programme in remodelled atria

Given the central role of FBs in structural and electrical atrial remodelling in AF, we next investigated their remodelling. Sub-clustering analysis of Multiome data resolved six FB states: APOD^+^, APOE^+^, basal, FAP–POSTN^+^, myofibroblast (MYO) and PTGDS^+^ populations (**Fig. 7a**). Their marker profiles were consistent with established human cardiac FB states^28^ (**Fig. 7b,c**; Supplementary Table 5). Group-level composition and fold-change summaries suggested expansion of the PTGDS^+^ state in the high-LAVI groups (**Fig. 7d,e**). Donor-level analysis showed that the PTGDS^+^ proportion was larger in Hi-SR than in Lo-SR (adjusted *P* = 0.041), whereas the Hi-AF–Lo-SR comparison did not reach significance (adjusted *P* = 0.097), reflecting substantial inter-individual variability (**Fig. 7f**). PTGDS^+^ FBs expressed a distinct programme including *IL6*, *NR4A3* and *SEMA4A*, and their enriched processes included IL-6 signalling, IL-1β production, TGF-β responses, collagen-fibril and ECM organisation, cell migration and adhesion (**Fig. 7g,h**; Supplementary Tables 7 and 8). Thus, PTGDS^+^ FBs define an inflammatory–fibrotic state associated primarily with structural remodelling rather than uniformly with AF alone.

**Fig. 7.**
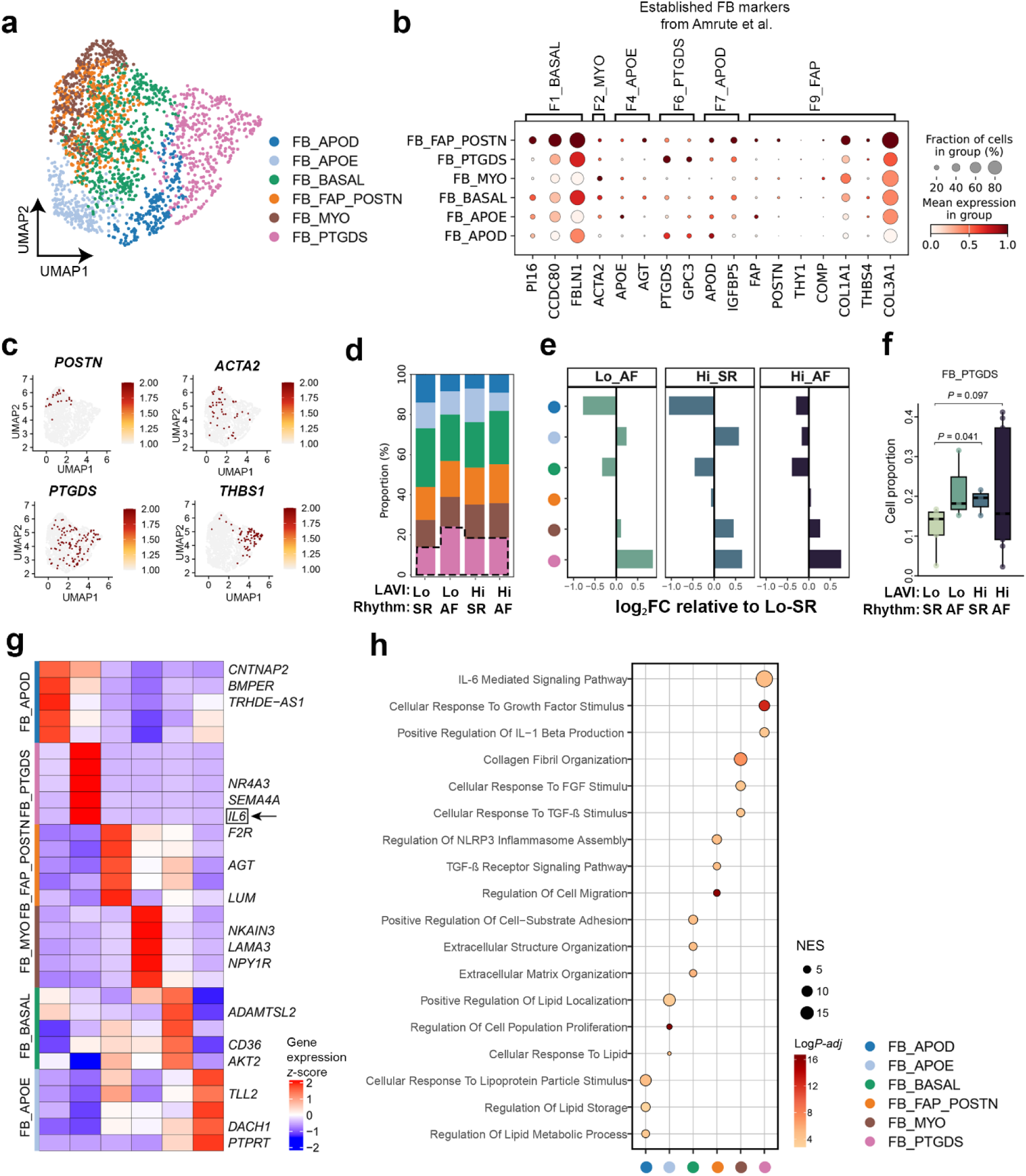
PTGDS^+^ FBs expand with atrial myopathy and show an inflammatory–fibrotic programme. **a**, UMAP of FB identifying six states: APOD^+^, APOE^+^, basal, FAP–POSTN^+^, myofibroblast (MYO) and PTGDS^+^ FB. **b**, Expression of established FB-state markers reported by Amrute *et al*^28^. across the six states. Point size denotes the fraction of nuclei expressing each gene, and colour denotes mean expression within the state. **c**, UMAP feature plots showing normalized expression of *POSTN*, *ACTA2*, *PTGDS* and *THBS1* in FBs. **d**, Relative composition of the six FB states across the Hi-AF, Hi-SR, Lo-AF and Lo-SR groups. The dashed outline highlights the PTGDS^+^ FB state. **e**, State-resolved differences in relative abundance for Hi-AF, Hi-SR and Lo-AF compared with Lo-SR. Values represent log₂(mean proportion in the indicated group/mean proportion in Lo-SR); positive and negative values indicate higher and lower average abundance, respectively. **f**, Donor-level proportions of PTGDS^+^ FB across the four groups. Each point represents one donor; Hi-AF, *n* = 9; Hi-SR, *n* = 3; Lo-AF, *n* = 3; and Lo-SR, *n* = 5. Differences were assessed using unpaired two-sided Wilcoxon rank-sum tests with Benjamini–Hochberg correction; adjusted *P* values are shown for the indicated comparisons. Boxplots show the median and interquartile range, with whiskers extending to the most extreme observations within 1.5 times the interquartile range. **g**, Heatmap of row-scaled expression of representative state-enriched genes across the six FB states; *IL6* is highlighted. **h**, Selected GOBP terms enriched in each FB state. Point size denotes the normalized enrichment score (NES), and colour denotes −log₁₀ of the adjusted *P-value*. FB-state colours are consistent across panels. FB, fibroblast; GOBP, Gene Ontology biological process.

### Enhancer-driven networks identify regulators of FB-state transitions

To identify candidate regulators of FB-state remodelling, we integrated TF expression with SCENIC+-inferred enhancer-driven regulatory networks (eRegulons), in which concordant target-gene expression and target-region accessibility provide an estimate of regulatory programme activity. Gene-based eRegulon activity and accessibility-based region activity delineated lineage-selective regulatory programmes across the major atrial cell types (**Fig. 8a**). The corresponding TF-expression profiles provided an additional cell-type-resolved view of these candidate regulators (**Fig. 8b**). Ranking the inferred networks by FB eRegulon specificity prioritised *GLIS3*, *TWIST2* and *TBX20* as the three most FB-selective eRegulons (**Fig. 8c**; Supplementary Table 11). Their expression and accessibility varied across FB states and rhythm–LAVI groups (Supplementary Fig. 15a–i). At the FB-state level, *TWIST2* expression was concentrated predominantly in PTGDS^+^ FBs (Supplementary Fig. 15a), whereas *TBX20* and *GLIS3* showed broader but state-dependent expression, with *TBX20* highest in PTGDS^+^ FBs (Supplementary Fig. 15d,g). Across rhythm–LAVI groups, expression of all three TFs was higher in Hi-AF than in Lo-SR (Supplementary Fig. 15b,e,h). Aggregate ATAC-seq profiles further revealed higher chromatin accessibility at the *TWIST2*, *TBX20,* and *GLIS3* loci in Hi-AF than in Lo-SR (Supplementary Fig. 15c,f,i). A trajectory rooted in basal FBs extended towards the PTGDS^+^ state, with *GLIS3* and *TBX20* increasing with pseudotime (**Fig. 8d**; Supplementary Fig. 15j,k). *NR4A1* also increased along this inferred trajectory, consistent with an independently described activated atrial FB programme^51^. Modelling of TF loss of function using CellOracle predicted that loss of each candidate TF redirected the local vector field towards the basal region, whereas simulated overexpression favoured movement towards the PTGDS^+^ region (**Fig. 8e**). The gene regulatory network (GRN) connected the three TFs to partly overlapping networks (**Fig. 8f**) whose predicted targets were enriched for ECM, TGF-β and fibrosis-related processes (Supplementary Fig. 15l).

**Fig. 8.**
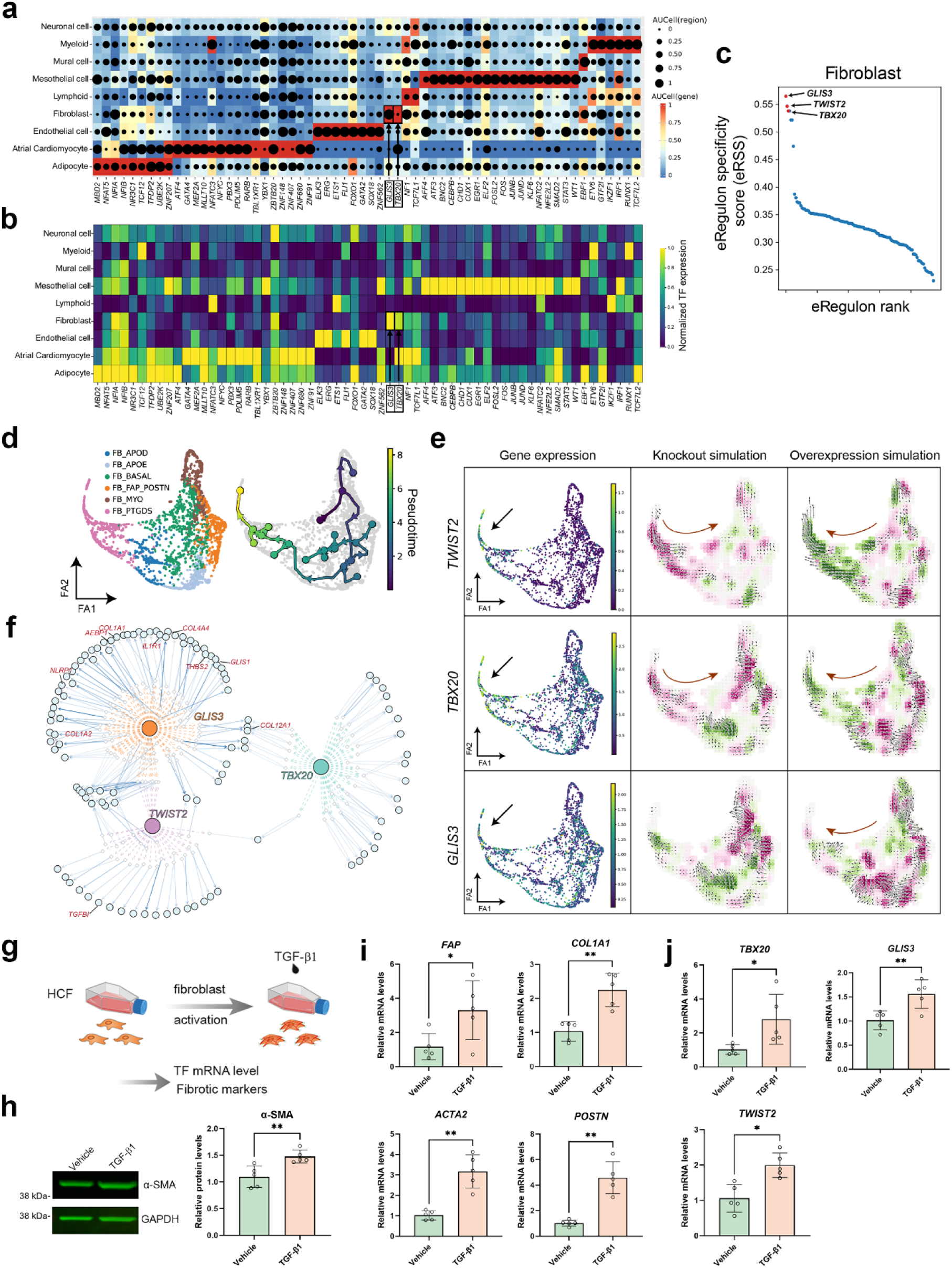
Multiomic regulon analysis identifies *TBX20*, *GLIS3* and *TWIST2* as FB-state regulators. **a**, SCENIC+ enhancer-driven regulon (eRegulon) activity (computed with AUCell based on target gene expression and target region accessibility) across major cell types. Point size denotes region-based AUCell score and colour denotes gene-based AUCell score. **b**, Expression of the corresponding TFs across cell types. FB-associated TF highlighted in a, b. **c**, FB eRegulons ranked by eRegulon specificity score (eRSS), highlighting the top three TFs: *GLIS3*, *TWIST2* and *TBX20*. **d**, Force-directed (FA) FB embedding coloured by cell state (left) and scFates pseudotime (right). **e**, Observed expression (left) and CellOracle-predicted vector fields following in silico knockout (middle) or overexpression (right) of *TWIST2*, *TBX20* or *GLIS3*. **f**, SCENIC+-inferred eRegulon networks for the three TFs. Circles denote genes and diamonds regulatory regions; TF–region edges are coloured by regulator and region–gene edges encode peak-to-gene importance; the coloured circles (orange, green, purple) indicate regulators; fibrosis-associated targets are labelled in red. **g**, Experimental design for validation in primary human cardiac fibroblasts (HCFs). Cells were treated with vehicle or TGF-β1 (50 ng/mL, 48 h), followed by measurement of candidate TFs and FB-activation markers. **h**, Representative immunoblots of α-smooth-muscle actin (α-SMA) and GAPDH in vehicle- and TGF-β1-treated HCFs (left) and quantification of α-SMA protein abundance (right). **i**, Relative transcript abundance of *FAP*, *COL1A1*, *ACTA2* and *POSTN* after TGF-β1 stimulation. **j**, Relative transcript abundance of *TBX20*, *GLIS3*, and *TWIST2*. For h–j, points represent independent experiments (*n* = 5), bars show the mean, and error bars indicate s.d. Statistical significance was assessed using two-sided paired *t*-tests. α-SMA abundance was normalized to GAPDH. \**P* < 0.05; \*\**P* < 0.01. TF, transcription factor.

We next tested whether these candidate regulators responded to profibrotic stimulation in primary human cardiac fibroblasts (HCFs). HCFs were treated with TGF-β1, followed by measurement of FB activation markers and candidate TF expression (**Fig. 8g**). Immunoblotting showed increased α-SMA protein following TGF-β1 stimulation (**Fig. 8h**). Consistent with FB activation, TGF-β1 increased the transcript levels of *FAP*, *COL1A1*, *ACTA2* and *POSTN* (**Fig. 8i**). TGF-β1 also induced *TBX20*, *GLIS3* and *TWIST2* expression (**Fig. 8j**), experimentally supporting the responsiveness of the three regulators prioritised by Multiome and SCENIC+ analyses.

### JMJD1C is a required epigenetic regulator of the FB TGF-β1 response

Since epigenetic regulators play a central role in translating extracellular signals into transcriptional programmes and provide a context for TF activity, we screened chromatin-modifying genes expressed across atrial FB states and rhythm–LAVI groups (**Fig. 9a**; Supplementary Fig. 16). The H3K9me2 demethylase *JMJD1C* was prominent and showed its highest expression in PTGDS^+^ FBs (**Fig. 9b**). *JMJD1C* expression was also higher in Hi-AF than in Lo-SR (*P* = 0.038; **Fig. 9c**). In cultured HCFs, TGF-β1 increased *JMJD1C* mRNA, protein abundance and nuclear fluorescence (Supplementary Fig. 17), linking JMJD1C to the FB response to profibrotic stimulation.

**Fig. 9.**
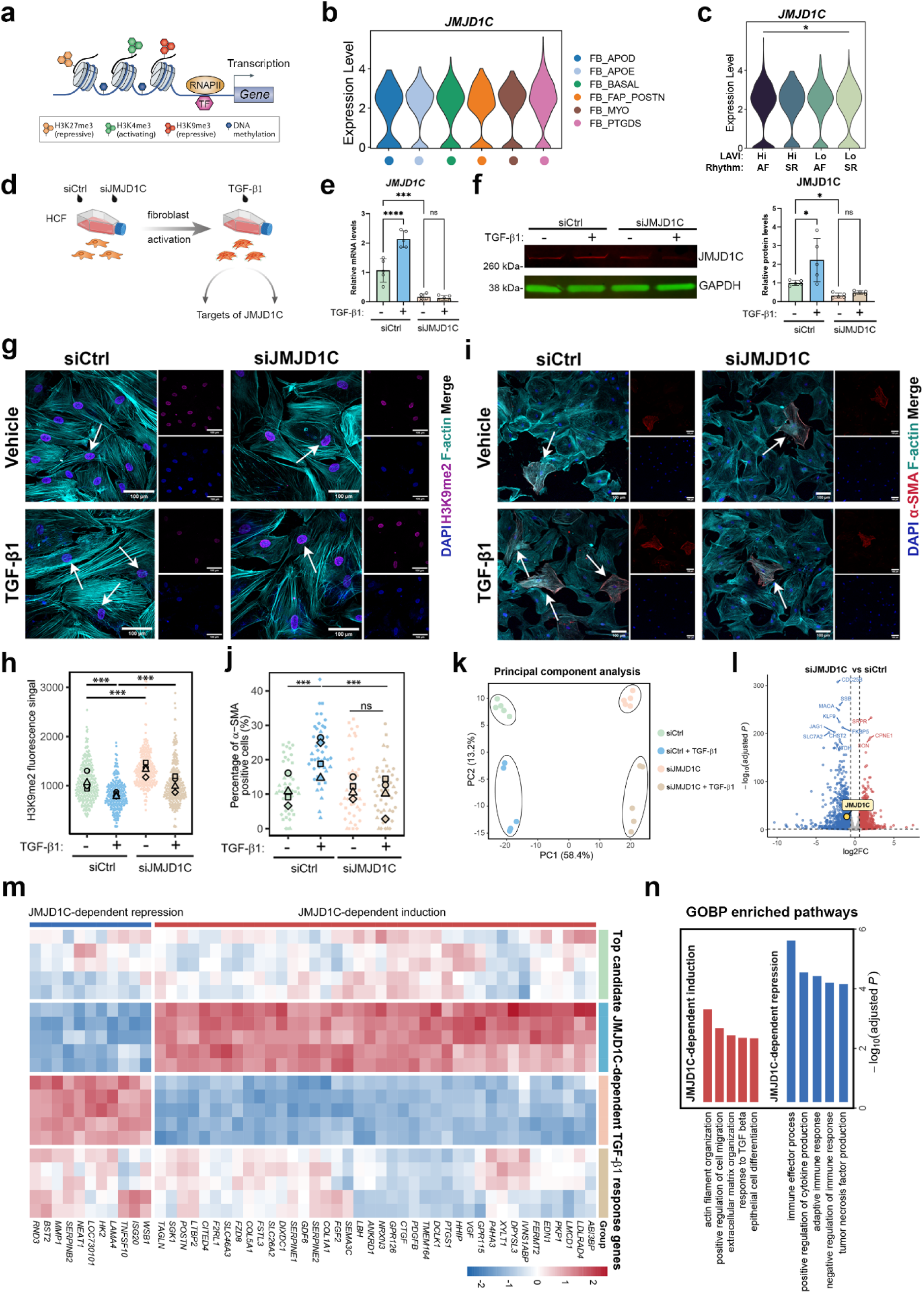
JMJD1C regulates the TGF-β1-responsive FB programme. **a**, Schematic of chromatin modifications and enzymes that regulate transcription. **b**, *JMJD1C* expression across FB states, showing higher expression in PTGDS^+^ FB. **c**, *JMJD1C* expression across disease groups; a two-sided Wilcoxon rank-sum test was used. **d**, Experimental design for *JMJD1C* knockdown in primary HCFs with or without TGF-β1 (50 ng/mL, 48 hours), followed by identification of targets of JMJD1C. **e**,**f**, Knockdown efficiency measured by RT-qPCR (**e**) and immunoblotting (**f**) (*n* = 5; ordinary one-way ANOVA with Tukey multiple testing correction). **g**, Representative DAPI (blue), H3K9me2 (purple) and F-actin (cyan) images after *JMJD1C* knockdown and TGF-β1 stimulation. **h**, Swarm plot of nuclear H3K9me2 intensity in g. Each small symbol represents one nucleus, whereas larger symbols indicate the mean of an independent experiment. **i**, Representative images of cells after *JMJD1C* knockdown and TGF-β1 immunostained for α-smooth-muscle actin (α-SMA) (red) and F-actin (cyan). Nuclei are indicated in blue by DAPI staining. **j**, Swarm plot of quantitation of α-SMA-positive cells in i. Each small symbol represents one image, whereas larger symbols indicate the mean of an independent experiment. Statistical analysis in h,j was performed using a linear mixed-effects model with condition as a fixed effect and technical measurements nested within biological replicate as random effects. Pairwise comparisons of estimated marginal means were adjusted using Tukey’s method. Scale bars, 100 μm. **k**, Principal-component analysis (PCA) of bulk RNA-seq profiles from four conditions (*n* = 5). **l**, Volcano plot showing genes differentially expressed (DESeq2) between siJMJD1C and control, with significant DEG defined by adjusted *P* < 0.05 and |log_2_FC| ≥ 0.58; top genes are labelled and JMJD1C is highlighted. **m**, Row-scaled expression of genes whose induction or repression by TGF-β1 depends on JMJD1C. **n**, GOBP enrichment among JMJD1C-dependent induced and repressed genes. ns, not significant; \**P* ≤ 0.05, \*\**P* ≤ 0.01, \*\*\**P* ≤ 0.001, \*\*\*\**P* ≤ 0.0001. HCF, human cardiac fibroblast; DEG, differentially expressed genes; FC, fold change; GOBP, gene ontology biological process.

We then tested whether *JMJD1C* contributed functionally to this response by depleting it with siRNA prior to vehicle or TGF-β1 exposure (**Fig. 9d**). RT–qPCR and immunoblotting confirmed efficient knockdown in both control and TGF-β1-stimulated conditions (**Fig. 9e,f**). Consistent with JMJD1C-dependent regulation of H3K9 methylation, TGF-β1 reduced global H3K9me2 fluorescence in control cells, whereas this reduction was significantly impaired by JMJD1C depletion (**Fig. 9g,h**). JMJD1C depletion also reduced the proportion of α-SMA-positive activated FB after TGF-β1 stimulation (**Fig. 9i,j**).

To gain deeper insights into the role of *JMJD1C* knockdown on the response of FB to TGF-β1 exposure, we performed bulk RNA sequencing of FB ± JMJD1C siRNA ± TGF-β1 (*n* = 5). Principal-component analysis (PCA) separated the four groups along the first two components (**Fig. 9k**). Pairwise contrasts identified baseline effects of *JMJD1C* depletion and TGF-β1-responsive genes whose induction or repression depended on JMJD1C (**Fig. 9l,m**). JMJD1C-dependent induced genes were enriched for actin-filament organisation, cell migration, ECM organisation and responses to TGF-β, whereas JMJD1C-dependent repressed genes were enriched for immune-effector and cytokine-regulatory processes (**Fig. 9n**). Expanded DE (Supplementary Fig. 18a–d), Hallmark (Supplementary Fig. 18e–h), GOBP (Supplementary Fig. 18i,j) and gene-program analyses (Supplementary Fig. 18k,l) confirmed that JMJD1C depletion reshaped epithelial–mesenchymal-transition (EMT), TGF-β, myogenesis, unfolded-protein-response and IFN-related programmes.

Finally, to identify candidate upstream regulators of *JMJD1C*, FB-subtype SCENIC+ analysis nominated *ELF2* and *KLF12* (Supplementary Fig. 19a,b). Their predicted eRegulons showed state-dependent activity; both factors were connected to candidate regulatory regions linked to *JMJD1C*, and their expression varied across rhythm–LAVI groups and FB states (Supplementary Fig. 19c,d,f,g; Supplementary Table 11). Validating this pathway, TGF-β1 stimulation increased *ELF2* and *KLF12* transcript abundance in HCF (Supplementary Fig. 19e,h). These findings provide candidate upstream regulators of *JMJD1C* and, together with the results of the knockdown experiments, identify *JMJD1C* as a functional modulator of a substantial subset of the FB TGF-β1 response.

### Myeloid–FB analysis predicts macrophage-derived TGF-β signalling

Owing to its established role in cardiac FB activation and its recurrent presence as a pro-fibrotic signal in our analysis of the LAA FB-state and HCF experiments, we next investigated cellular sources of TGF-β1. Myeloid subclustering identified CD14^+^ monocytes, intermediate macrophages, three LYVE1^+^ macrophage states, and dendritic cells (DCs), with group-dependent differences in their relative abundance (**Fig. 10a,b**). Cell-cell interaction analysis using CellPhoneDB predicted TGF-β signalling from macrophage states to TGF-β-receptor complexes on PTGDS^+^ FBs in each rhythm–LAVI group (**Fig. 10c**). Broader analysis of the TGF-β superfamily also predicted interactions with APOD^+^, APOE^+^, basal, FAP–POSTN^+^ and MYO states, including basal and FAP–POSTN populations positioned earlier than PTGDS^+^ FBs on the inferred trajectory (Supplementary Fig. 20). This alignment is compatible with macrophage-derived TGF-β-family signals contributing both to entry of FB into a remodelling trajectory and to maintenance of a terminal inflammatory–fibrotic state. Together, these results connect atrial macrophages with the experimentally supported TGF-β–JMJD1C FB programme and define testable models of immune–stromal signalling in atrial remodelling.

**Fig. 10.**
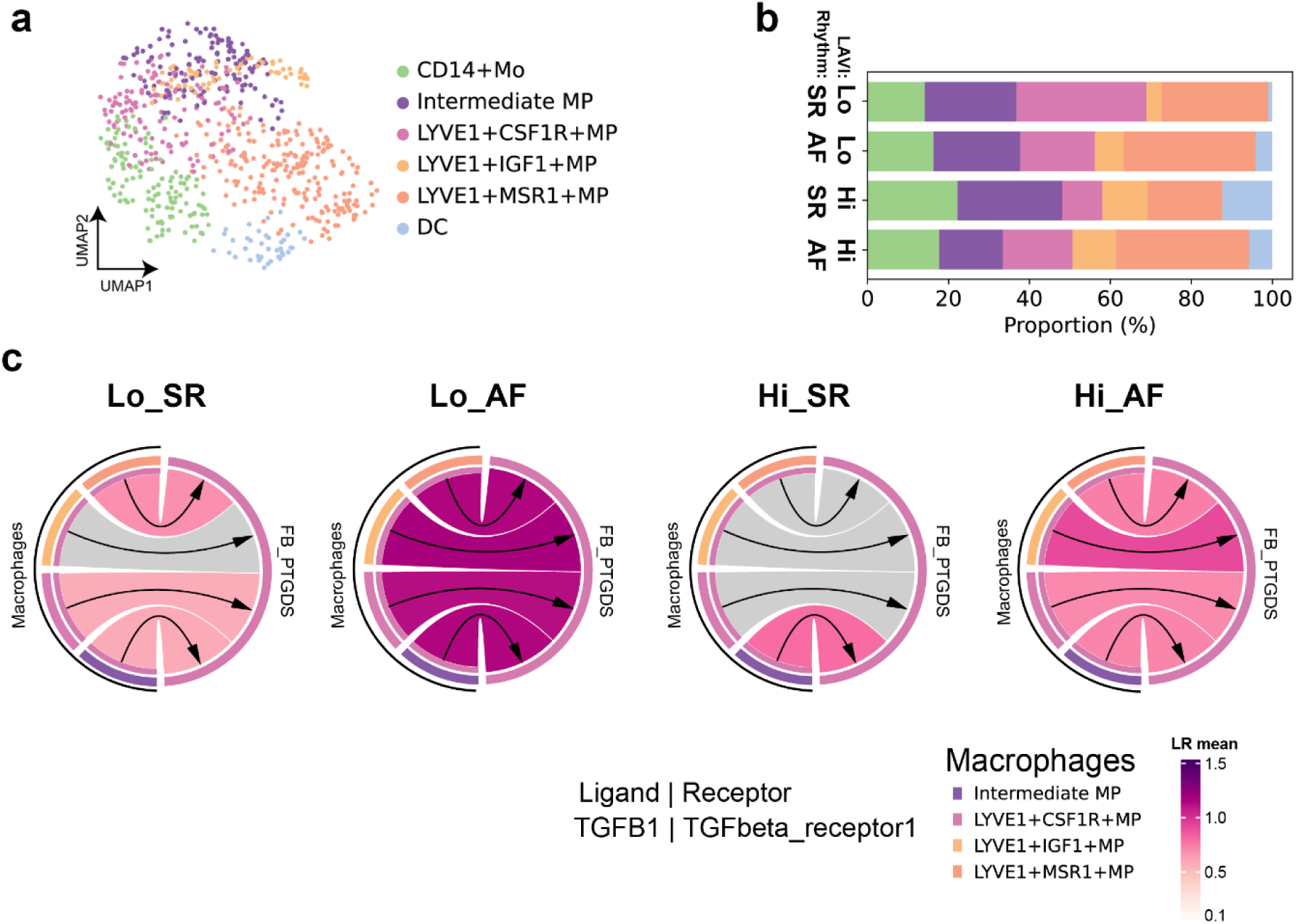
Macrophages communicate with PTGDS^+^ FBs through predicted TGF-β signalling. **a**, UMAP of myeloid nuclei resolving CD14^+^ monocytes, intermediate macrophages, three LYVE1^+^ macrophage states and dendritic cells (DCs). **b**, Relative abundance of these states across groups. **c**, CellPhoneDB-predicted TGFB1–TGF-β receptor signalling from macrophage states to PTGDS^+^ FB in each group. Ribbon colour encodes the mean expression of the ligand–receptor pair as indicated; arrows denote the macrophage-to-FB direction.

## Discussion

By integrating chamber-resolved snRNA-seq, paired RNA–ATAC profiling, histology, regulatory-network inference and functional validation, this study delineates cellular programmes associated with AF and structural atrial remodelling. AF was accompanied by changes in cell type/cell state proportions and gene expression in both atria, although LAA and RAA exhibited different remodelling profiles. Within LAA, the combined single-nucleus gene expression/epigenome analysis resolved a stressed aCM4 and an inflammatory PTGDS^+^ FB state with distinct candidate *cis*-regulatory architectures. Functional experiments identified that *JMJD1C* lies downstream of the TGF-β1 response in human cardiac FB, while ligand–receptor analysis proposed atrial macrophages as a potential source of TGF-β signalling. Together, our data link rhythm, structural remodelling, and cell-state diversity with epigenetic regulation of gene expression across CM, stromal and immune interactions underlying atrial remodelling.

The rhythm–LAVI design addresses a central feature of atrial remodelling: AF and AM are closely related but are also unique states. The 2024 EHRA/HRS/APHRS/LAHRS consensus emphasizes that an atrial myopathic substrate may precede AF, whereas AF can accelerate electrical, contractile and structural remodelling^9^. Accordingly, in our study, Hi-AF is considered a combined AF/high-remodelling phenotype, Hi-SR as structural remodelling in the absence of AF, Lo-AF as AF without overt atrial enlargement, and Lo-SR as a low-remodelling SR reference phenotype. However, these groups should not be equated directly with the EHRAS histopathological classes or proposed clinical stages, because LAVI is a measurement of atrial geometry rather than a diagnosis of AM. Within this design, the pairwise molecular comparisons and increased fibrosis in both high-LAVI groups support separable rhythm-and structure-associated processes. Additionally, the chamber-resolved atlas further showed the complexity of atrial remodelling across both atria, where common disease exposure did not produce identical responses: LAA preferentially engaged ECM, immune, and TGF-β programmes. These observations significantly extend insights provided by chamber-resolved human heart atlases^19^ by analysing anatomical identity together with rhythm and structural-remodelling burden.

The findings from the analysis of CMs connect acquired atrial remodelling to the cell type most strongly implicated by AF genetics. Previous cell-type-resolved chromatin studies have localized significant common AF risk to CM regulatory elements^21^, and large-scale atrial single-nucleus studies have identified CM stress-response genes in AF^20^. Multiomic analysis has also implicated androgen receptor (AR) signalling in AF CMs^25^. Here, an NPPA^+^ and NPPB^+^ stressed aCM4 state was increased in Hi-AF relative to Lo-SR and showed altered regulatory connectivity across rhythm–LAVI phenotypes. The inferred CTCF-centred network is compelling since this TF functions to organize enhancer–promoter communication and chromatin-domain boundaries^52^. Moreover, its depletion in CMs alters chromatin organisation and enhancer–gene interactions^53,54^. At the AF-associated *PITX2* locus, deletion of CTCF-binding boundary elements disrupts chromatin-domain organisation and *PITX2* regulation, linking altered genome architecture to sinoatrial-node dysfunction and atrial arrhythmias^55^.

FB diversification and increased heterogeneity are a prominent feature of cardiac remodelling during pathology and HF^28,56^. Human HF atlases have shown that CMs often converge on shared stress states while FB and myeloid populations diversify^57^. Further, chronically remodelled ventricles are populated by activated *POSTN*- and *AEBP1*-associated FBs^58^. PTGDS^+^ FBs have also been resolved in ventricular HF, where a PTGDS*/*GPC3 state was distinct from the FAP–POSTN and MYO populations that preferentially expanded with disease^28^. In atria, an inflammatory NR4A1⁺ FB state has likewise been reported^51^. The atrial PTGDS⁺ state described here shares a non-canonical inflammatory identity but is distinguished by concurrent IL6, ECM and TGF-β programmes, enrichment of *JMJD1C*, and preferential representation in high-LAVI atria. Its abundance was greater in both high-LAVI groups than in Lo-SR, reaching statistical significance in Hi-SR. Rather than resembling a conventional contractile MYO, its combined inflammatory and matrix remodelling programme is consistent with a state that may help sustain a pro-fibrotic atrial niche. Its presence in Hi-SR further suggests that FB-state remodelling can accompany atrial enlargement before clinically manifest AF.

The identification of *JMJD1C* as a state-enriched, TGF-β-responsive transcriptional regulator provides a mechanistic bridge between the PTGDS⁺ transcriptional programme and epigenetic remodelling. *JMJD1C* expression was highest in PTGDS⁺ FBs and increased after TGF-β1 stimulation, placing it downstream of receptor activation. *JMJD1C* depletion modified the TGF-β1-associated change in global H3K9me2, reduced α-SMA^+^ cells, and selectively rewired both induced and repressed transcriptional responses, including ECM and immune programmes. These convergent chromatin, immunofluorescence, and RNA-seq readouts support a role for *JMJD1C* in setting the amplitude and composition of FB activation rather than simply marking the PTGDS⁺ state.

Together, these findings position H3K9me2 regulation as a potential link between extracellular profibrotic signals and persistent transcriptional remodelling. In vCMs, pathological hypertrophy is associated with miR-217-dependent loss of EHMT1/2 and H3K9me2; genetic or pharmacological EHMT1/2 inactivation is sufficient to reactivate foetal gene programmes and induce hypertrophy, whereas preservation of this pathway is protective^59^. Conversely, JMJD1C removes repressive H3K9 methylation, and previous cardiac studies showed that *JMJD1C* silencing or treatment with the broad *Jmjd1c* demethylase inhibitor JIB-04 attenuated Ang-II-induced hypertrophy, fibrotic-gene expression, and *TIMP1* transcription^60^. The therapeutic potential of epigenetic intervention in the atrium is further supported by inhibition of the H3K27 methyltransferase EZH2, which reduced atrial FB differentiation, fibrosis, and AF vulnerability in experimental models^61^. Moreover, the development of the orally active JMJD1C inhibitor 193D7 for experimental cancer immunotherapy demonstrates that JMJD1C is pharmacologically tractable^62^. Collectively, these findings support selective, ideally FB-targeted, JMJD1C inhibition as a testable strategy for limiting TGF-β-driven atrial fibrosis. However, the lineage-specific functions of H3K9me2 regulators necessitate careful assessment of target selectivity, cardiac electrophysiology, immune effects, and systemic toxicity.

The predicted intercellular signalling from macrophages to FB place this activity within a broader inflammatory niche. Recruited SPP1⁺ macrophages promote atrial fibrosis and AF susceptibility in an experimental environment^23^, whereas human HF studies show that CCR2⁺ macrophage-derived IL-1β can induce FAP–POSTN FB emergence and that interrupting this axis reduces ventricular fibrosis *in vivo*^28^. In our snRNA-Seq analysis, multiple macrophage states expressed TGF-β, and PTGDS⁺ FBs expressed compatible TGF-β receptor complexes across rhythm–LAVI groups. Since PTGDS⁺ FBs locate at the distal end of the inferred trajectory, this interaction could be plausibly interpreted as either maintaining the established inflammatory–matrix state or initiating the basal-to-PTGDS⁺ transition. The convergence of this predicted interaction with TGF-β-dependent *JMJD1C* induction in primary human cardiac FBs supports a coherent macrophage–TGF-β–*JMJD1C* axis, while remaining an inference from ligand–receptor expression rather than direct evidence of signal transfer.

The principal findings of this study were made possible through the generation of the first human atrial dataset to integrate chamber identity, rhythm status and structural remodelling with paired transcriptomic–epigenomic profiling and functional testing. This study design enabled isolation of molecular features associated with AF from those accompanying atrial enlargement and to link a high-LAVI-associated PTGDS⁺ FB state to candidate enhancer networks, a TGF-β-responsive *JMJD1C* programme and macrophage-derived signals. Together with the stressed aCM4 regulatory landscape, these findings support a multicellular model in which CM stress, FB-state remodelling and immune–stromal signalling jointly shape the atrial substrate. *JMJD1C* emerges as an epigenetic regulatory target, providing a focused basis for future evaluation of cell-state-selective antifibrotic strategies.

## Author contributions statement

Conceptualization and study design: H.L.R., V.F.M.S., G.W.D.K. and H.H. Investigation: M.R.L.T., G.W.D.K., H.H. and V.F.M.S. collected the human biopsies; Y.F. and A.V. performed nuclear isolation; Y.F. and L.L. performed cell culture, qPCR, immunoblotting and immunofluorescence; and J.T. performed qPCR. Formal analysis: Y.F. analysed the snRNA-seq and snMultiome data. Interpretation: B.T. and S.T. contributed to interpretation of the snRNA-seq analyses. Writing—original draft: Y.F. and H.L.R. Writing—review and editing: all authors. All authors read and approved the final manuscript.

## Acknowledgements

The authors thank Roxane Menten for expert lab management and Genomics Core Leuven for 10x snMultiome sequencing and the Flow Cytometry Core Facility for technical assistance with nuclei sorting.

## References

1. Ko, D., Chung, M. K., Evans, P. T., Benjamin, E. J. & Helm, R. H. Atrial Fibrillation: A Review. JAMA 333, 329–342 (2025).

2. Elsheikh, S., Hill, A., Irving, G., Lip, G. Y. H. & Abdul-Rahim, A. H. Atrial fibrillation and stroke: State-of-the-art and future directions. Current Problems in Cardiology 49, 102181 (2024).

3. Anter, E., Jessup, M. & Callans, D. J. Atrial Fibrillation and Heart Failure. Circulation 119, 2516–2525 (2009).

4. Rivard, L. et al. Atrial Fibrillation and Dementia: A Report From the AF-SCREEN International Collaboration. Circulation 145, 392–409 (2022).

5. Van Gelder, I. C. et al. 2024 ESC Guidelines for the management of atrial fibrillation developed in collaboration with the European Association for Cardio-Thoracic Surgery (EACTS). European Heart Journal 45, 3314–3414 (2024).

6. Cheng, S. et al. Global burden of atrial fibrillation/atrial flutter and its attributable risk factors from 1990 to 2021. Europace 26, euae195 (2024).

7. Nattel, S., Heijman, J., Zhou, L. & Dobrev, D. Molecular basis of atrial fibrillation pathophysiology and therapy: a translational perspective. Circulation Research 127, 51–72 (2020).

8. Williams, D. O., Jones, E. L., Nagle, R. E. & Smith, B. S. Familial Atrial Cardiomyopathy with Heart Block. QJM 41, 491–508 (1972).

9. Goette, A. et al. Atrial cardiomyopathy revisited—evolution of a concept: a clinical consensus statement. Europace 26, euae204 (2024).

10. Tubeeckx, M. R. L., De Keulenaer, G. W., Heidbuchel, H. & Segers, V. F. M. Pathophysiology and clinical relevance of atrial myopathy. Basic Res Cardiol 119, 215–242 (2024).

11. Maguy, A., Mahendran, Y., Tardif, J.-C., Busseuil, D. & Li, J. Autoimmune Atrial Fibrillation. Circulation 148, 487–498 (2023).

12. Yamazoe, M. et al. B cells promote atrial fibrillation via autoantibodies. Nat Cardiovasc Res 4, 1381–1396 (2025).

13. Nattel, S. Molecular and Cellular Mechanisms of Atrial Fibrosis in Atrial Fibrillation. JACC: Clinical Electrophysiology 3, 425–435 (2017).

14. Raniga, D. et al. Left atrial volume index: A predictor of atrial fibrillation recurrence following direct current cardioversion—a systematic review and meta-analysis. International Journal of Cardiology: Heart & Vasculature 51, 101364 (2024).

15. Miranda, A. M. A. et al. Single-cell transcriptomics for the assessment of cardiac disease. Nat Rev Cardiol 20, 289–308 (2023).

16. Gardner, R. S., Tucker, N. R. & Amancherla, K. Leveraging Single-Cell Technologies to Advance Understanding of Myocardial Disease. Circulation Research 138, e326002 (2026).

17. Tang, Z. et al. The future of cardiology: Integrating single-cell transcriptomics with multi-omics for enhanced cardiac disease insights. Current Problems in Cardiology 50, 103005 (2025).

18. Abplanalp, W. T., Tucker, N. & Dimmeler, S. Single-cell technologies to decipher cardiovascular diseases. Eur Heart J 43, 4536–4547 (2022).

19. Kanemaru, K. et al. Spatially resolved multiomics of human cardiac niches. Nature 619, 801–810 (2023).

20. Hill, M. C. et al. Large-scale single-nuclei profiling identifies role for ATRNL1 in atrial fibrillation. Nat Commun 15, 10002 (2024).

21. Selewa, A. et al. Single-cell genomics improves the discovery of risk variants and genes of atrial fibrillation. Nature Communications 14, 4999 (2023).

22. Sheng, Y. et al. Deciphering mechanisms of cardiomyocytes and non-cardiomyocyte transformation in myocardial remodeling of permanent atrial fibrillation. Journal of Advanced Research 61, 101–117 (2024).

23. Hulsmans, M. et al. Recruited macrophages elicit atrial fibrillation. Science 381, 231–239 (2023).

24. Suzuki, Y. et al. Left atrial single-cell transcriptomics reveals amphiregulin as a surrogate marker for atrial fibrillation. Commun Biol 7, 1601 (2024).

25. Leblanc, F. J. A. et al. Single-nucleus multi-omics implicates androgen receptor signaling in cardiomyocytes and NR4A1 regulation in fibroblasts during atrial fibrillation. Nat Cardiovasc Res 4, 433–444 (2025).

26. Preissl, S., Gaulton, K. J. & Ren, B. Characterizing cis-regulatory elements using single-cell epigenomics. Nature Reviews Genetics 24, 21–43 (2023).

27. Hocker, J. D. et al. Cardiac cell type–specific gene regulatory programs and disease risk association. Science Advances 7, eabf1444 (2021).

28. Amrute, J. M. et al. Targeting immune–fibroblast cell communication in heart failure. Nature 635, 423–433 (2024).

29. Xie, Y. et al. Single-cell multiomics and chromatin structure reveal gene-regulatory dynamics in heart failure. Science 393, eady6893 (2026).

30. Fu, Y. et al. Benchmarking of computational demultiplexing methods for single-nucleus RNA sequencing data. Brief Bioinform 26, bbaf371 (2025).

31. Heumos, L. et al. Best practices for single-cell analysis across modalities. Nat Rev Genet 1–23 (2023) doi:10.1038/s41576-023-00586-w.

32. Lopez, R., Regier, J., Cole, M. B., Jordan, M. I. & Yosef, N. Deep generative modeling for single-cell transcriptomics. Nat Methods 15, 1053–1058 (2018).

33. Wolf, F. A., Angerer, P. & Theis, F. J. SCANPY: large-scale single-cell gene expression data analysis. Genome Biol 19, 15 (2018).

34. Dann, E., Henderson, N. C., Teichmann, S. A., Morgan, M. D. & Marioni, J. C. Differential abundance testing on single-cell data using k-nearest neighbor graphs. Nat Biotechnol 40, 245–253 (2022).

35. Hao, Y. et al. Integrated analysis of multimodal single-cell data. Cell 184, 3573–3587.e29 (2021).

36. Granja, J. M. et al. ArchR is a scalable software package for integrative single-cell chromatin accessibility analysis. Nat Genet 53, 403–411 (2021).

37. Gaspar, J. M. Improved peak-calling with MACS2. 496521 Preprint at 10.1101/496521 (2018).

38. Stuart, T., Srivastava, A., Madad, S., Lareau, C. A. & Satija, R. Single-cell chromatin state analysis with Signac. Nat Methods 18, 1333–1341 (2021).

39. Schep, A. N., Wu, B., Buenrostro, J. D. & Greenleaf, W. J. chromVAR: inferring transcription-factor-associated accessibility from single-cell epigenomic data. Nat Methods 14, 975–978 (2017).

40. Faure, L., Mollet, B., Liebermeister, W. & Faulon, J.-L. A neural-mechanistic hybrid approach improving the predictive power of genome-scale metabolic models. Nat Commun 14, 4669 (2023).

41. Setty, M. et al. Characterization of cell fate probabilities in single-cell data with Palantir. Nat Biotechnol 37, 451–460 (2019).

42. Bravo González-Blas, C., et al. SCENIC+: single-cell multiomic inference of enhancers and gene regulatory networks. Nat Methods 20, 1355–1367 (2023).

43. Feng, J., Liu, T., Qin, B., Zhang, Y. & Liu, X. S. Identifying ChIP-seq enrichment using MACS. Nat Protoc 7, 1728–1740 (2012).

44. Efremova, M., Vento-Tormo, M., Teichmann, S. A. & Vento-Tormo, R. CellPhoneDB: inferring cell–cell communication from combined expression of multi-subunit ligand–receptor complexes. Nat Protoc 15, 1484–1506 (2020).

45. Troulé, K. et al. CellPhoneDB v5: inferring cell–cell communication from single-cell multiomics data. Nat Protoc 20, 3412–3440 (2025).

46. Love, M. I., Huber, W. & Anders, S. Moderated estimation of fold change and dispersion for RNA-seq data with DESeq2. Genome Biology 15, 550 (2014).

47. Wu, T. et al. clusterProfiler 4.0: A universal enrichment tool for interpreting omics data. The Innovation 2, 100141 (2021).

48. Subramanian, A. et al. Gene set enrichment analysis: A knowledge-based approach for interpreting genome-wide expression profiles. Proceedings of the National Academy of Sciences 102, 15545–15550 (2005).

49. Ritchie, M. E. et al. limma powers differential expression analyses for RNA-sequencing and microarray studies. Nucleic Acids Res 43, e47 (2015).

50. Kuppe, C. et al. Spatial multi-omic map of human myocardial infarction. Nature 608, 766–777 (2022).

51. Leblanc, F. J. A. et al. Single-nucleus multi-omics implicates androgen receptor signaling in cardiomyocytes and NR4A1 regulation in fibroblasts during atrial fibrillation. Nature Cardiovascular Research 4, 433–444 (2025).

52. Preissl, S., Gaulton, K. J. & Ren, B. Characterizing cis-regulatory elements using single-cell epigenomics. Nat Rev Genet 24, 21–43 (2023).

53. Nora, E. P. et al. Targeted Degradation of CTCF Decouples Local Insulation of Chromosome Domains from Genomic Compartmentalization. Cell 169, 930–944.e22 (2017).

54. Rosa-Garrido, M. et al. High-Resolution Mapping of Chromatin Conformation in Cardiac Myocytes Reveals Structural Remodeling of the Epigenome in Heart Failure. Circulation 136, 1613–1625 (2017).

55. Baudic, M. et al. TAD boundary deletion causes PITX2-related cardiac electrical and structural defects. Nat Commun 15, 3380 (2024).

56. Youness, M. et al. Location and aetiology are determinants of fibroblast activation and heterogeneity in the failing human heart. Genome Med 17, 155 (2025).

57. Koenig, A. L. et al. Single-cell transcriptomics reveals cell-type-specific diversification in human heart failure. Nat Cardiovasc Res 1, 263–280 (2022).

58. Kattih, B. et al. Single-nuclear transcriptome profiling identifies persistent fibroblast activation in hypertrophic and failing human hearts of patients with longstanding disease. Cardiovascular Research 119, 2550–2562 (2023).

59. Thienpont, B. et al. The H3K9 dimethyltransferases EHMT1/2 protect against pathological cardiac hypertrophy. J Clin Invest 127, 335–348 (2017).

60. Zhang, S., Lu, Y. & Jiang, C. Inhibition of histone demethylase JMJD1C attenuates cardiac hypertrophy and fibrosis induced by angiotensin II. Journal of Receptors and Signal Transduction 40, 339–347 (2020).

61. Zhang, S., Lu, Y. & Jiang, C. Inhibition of histone demethylase JMJD1C attenuates cardiac hypertrophy and fibrosis induced by angiotensin II. Journal of Receptors and Signal Transduction 40, 339–347 (2020).

62. Long, X. et al. Targeting JMJD1C to selectively disrupt tumor Treg cell fitness enhances antitumor immunity. Nat Immunol 25, 525–536 (2024).

