## Supplementary Materials for "Identification of JMJD1C as a key mediator of fibroblast remodelling in human AF by single-nucleus multi-omics"

### Supplementary methods

#### Primary processing, demultiplexing and quality control

##### Read processing and count matrices

Raw base calls were demultiplexed to FASTQ files and aligned to the 10x Genomics GRCh38-2020 reference. Gene-expression libraries were processed with Cell Ranger (v7.1.0)[1] and paired Multiome libraries with Cell Ranger ARC (v2.0.2)[2]. Gene-expression background counts were removed with CellBender (v0.3.0) using the default model settings[3]. Donor genotypes were called from aligned reads with cellsnp-lite (v1.2.3)[4] and pooled libraries were demultiplexed with Vireo (v0.5.8)[5]. Nuclei assigned as genotype doublets or with ambiguous donor identity were excluded.

##### Gene-expression quality control

Quality control was performed per library in Scanpy[6]. Genes detected in fewer than three nuclei were removed. Nuclei were retained when they contained at least 200 detected genes and at least 500 total counts, with mitochondrial and ribosomal transcript fractions each below 5%[7]. Scrublet (v0.2.3) was run separately for each library with its default expected doublet rate; nuclei with a Scrublet score of 0.30 or greater were removed[8]. Additional genotype doublets identified by Vireo were excluded.

#### Single-nucleus RNA-sequencing (snRNA-seq) analysis

##### Differential gene expression and pathway analysis

Differential expression (DE) in whole-chamber and cell-type summaries was performed in Seurat (v5.1.0) using FindMarkers on the SCT assay with MAST, two-sided testing, min.pct = 0.10, and LogFC. = 0.10[9,10]. AF was contrasted with SR unless otherwise stated. Seurat-adjusted *P* values correspond to Bonferroni correction across all tested features. In the cell-type panels of Fig. 3, highlighted genes in red and blue additionally satisfied adjusted $P < 0.05$ and |average log2 fold change| > 0.20; whole-chamber volcano panels used adjusted $P < 0.05$ without an additional display fold-change threshold. Pairwise comparisons among the four LAVI/rhythm groups used the same MAST settings. For Enrichr analysis, the top 200 positively ranked genes for each contrast direction were selected by adjusted MAST *P*-value and submitted to GO Biological Process (GOBP) 2023[11].

#### Single-nucleus chromatin-accessibility (snATAC-seq) analysis

##### Signac gene activity and peak-to-gene links

Paired Multiome data were additionally analysed with Signac (v1.8.0)[12]. Gene activity scores were calculated from accessibility surrounding annotated gene bodies and promoters, then normalised with LogNormalize using the median GeneActivity library size as the scale factor for the relevant analysis object. Cell-type-pseudobulk averages were used for cross-modality comparisons, and gene-wise RNA–gene-activity concordance was assessed with Spearman correlation across cell types. Genes were ranked by concordance; the upper and lower tails were used to illustrate loci with concordant or discordant accessibility and expression.

Peak-to-gene (P2G) associations in the paired left atrial appendage (LAA) analysis were inferred with Signac LinkPeaks using the peak-count assay and RNA data. Pearson correlation was evaluated for peaks within 500 kb of a gene, requiring at least 20 nuclei, pvalue_cutoff = 0.05, and score_cutoff = 0.05. For the final displayed set, links were retained when score ≥ 0.01, $P < 0.05$, and the linked feature was variable in both modalities, yielding 18,793 links.

##### Cardiomyocyte (CM) multimodal entropy

To quantify local RNA–chromatin concordance among atrial cardiomyocytes (aCM), separate k-nearest-neighbour (KNN) graphs were constructed from the scVI[13] gene-expression and PeakVI[14] accessibility embeddings with k = 50 and neighbourhood sampling proportion 0.10. For each RNA neighbourhood, memberships of its nuclei across ATAC neighbourhoods were converted to probabilities, and Shannon entropy was calculated in base 2. Entropy was regressed on RNA-neighbourhood size and the total within-neighbourhood variance of the scVI representation by ordinary least squares; the residual was used as adjusted entropy. Higher adjusted entropy therefore indicates that transcriptionally similar nuclei are dispersed over a larger number of accessibility neighbourhoods, consistent with lower local epigenetic coherence. Neighbourhood-average residuals were projected onto the CM UMAP for visualization.

#### Fibroblast-state and gene-regulatory analyses

##### CellOracle network inference and in silico perturbation

Fibroblast (FB) gene-regulatory networks (GRN) were analysed with CellOracle (v0.20.0)[15]. Raw counts, FB state labels, and the force directed (FA) embedding were imported; a promoter/enhancer base network derived from motif-scanned accessible regions was used as the regulatory prior. Cluster-specific links were fitted with ridge parameter α = 10 and filtered at $P < 0.001$, retaining the 2,000 largest absolute coefficients. Perturbation models were fitted with α = 10. Knockout (KO) simulations set the target-gene expression to zero and propagated regulatory effects for three iterations. Overexpression (OE) simulations set *TBX20* to 5, *GLIS3* to 4, or *TWIST2* to 2 in the model's expression scale. Transition vectors used 200 neighbours, randomised KNN sampling, and sampled_fraction = 1; embedding shifts used sigma_corr = 0.05 and were visualised on a 40 × 40 grid with smooth = 0.8 and 200 neighbours. Pseudotime-grid interpolation used 50 neighbours.

#### Histology and fibrosis quantification

LAA specimens were fixed in 4% paraformaldehyde (PFA), paraffin-embedded, and sectioned at 5 µm as previously described[16]. In brief, collagen distribution was visualized by Masson's trichrome staining. Images were acquired using either an Olympus BX43 microscope (Olympus Stream Motion Software) or a CELENA® S Digital Imaging System, using identical exposure and colour settings across groups. Images were analysed in ImageJ (v2.14.0) software[17]. Collagen-positive areas (blue) were segmented from each image and normalised to the analysed tissue area to obtain fibrosis fraction. Fibrosis fractions were log10-transformed before statistical analysis. For the four-group comparison, log10-transformed fibrosis fractions were analysed by one-way analysis of variance (ANOVA), followed by Tukey's multiple-comparison test.

#### Primary human cardiac fibroblast (HCF) experiments

##### RNA extraction, reverse transcription, and quantitative PCR

Total RNA was isolated from cultured FBs using the RNeasy Plus Mini Kit (Qiagen, Germany) according to the manufacturer's protocol. For each reaction, 100 ng RNA was combined with 1 µL random primers (200 ng/µL; Invitrogen, 48190011) and 1 µL 10 mM dNTPs (Thermo Scientific, R0192), heated at 65 °C for 5 min and cooled. Reverse-transcription mix was added to a final volume of 20 µL containing 4 µL 5× first-strand buffer, 1 µL 0.1 M DTT, 1 µL RNaseOUT, and 1 µL SuperScript III reverse transcriptase (200 U/µL; Invitrogen, 18080044). Reactions were incubated at 25 °C for 10 min, 50 °C for 60 min, and 70 °C for 15 min, then held at 4 °C. cDNA was diluted before quantitative PCR (qPCR).

qPCR was performed in 6 µL reactions on a Bio-Rad CFX384 instrument using GoTaq qPCR Master Mix (Promega, A6002). Each well contained 1.5 µL diluted cDNA and 4.5 µL master mix (3 µL 2× GoTaq mix, 0.75 µL primer mix to a final primer concentration of 0.2 µM, and 0.75 µL nuclease-free water). Cycling consisted of 95 °C for 3 min, 45 cycles of 95 °C for 15 s and 60 °C for 1 min, followed by melt-curve acquisition from 65 to 95 °C in 0.5 °C increments with 5 s per increment. Relative expression was calculated by the $2^{-\Delta\Delta Cq}$ method after normalization to the geometric mean of *18S*, *36B4*, and *RPL32* reference-gene Cq values and re-centring the control group to 1. Primer sequences are provided in Supplementary Table 12.

**Immunoblotting**

Cells were washed with PBS on ice and lysed in a modified RIPA buffer containing 150 mM NaCl, 20 mM Tris-HCl, pH 7.4, 10 mM EDTA, 1 mM DTT, 1% each of sodium deoxycholate and TRITON X-100, and 0.1 % SDS. The buffer was supplemented with protease and phosphatase inhibitor cocktails (Sigma-Aldrich, 11836170001) and with 0.25 U/µL of Benzonase (Novagen) to solubilize DNA. After 30 min incubation on ice, cell debris was subsequently removed by centrifugation at 10,000 x *g* for 10 mins at 4°C. Protein concentration of the lysate was determined by the BCA assay (Thermo, 23225). 20 µg of protein was resolved by SDS-PAGE on NuPAGE 4-12% Bis-Tris gradient gels (Invitrogen, NP0335BOX). Resolved proteins were electrophoretically transferred to 0.20 µm low fluorescence PVDF (Thermo, IB24002). Following transfer, non-specific protein binding sites were blocked by incubation for 1 h at room temperature in 0.5× Odyssey blocking buffer (LiCOR) diluted in TBS. Primary antibodies were diluted in 0.5x Odyssey blocking buffer/TBS and incubated overnight at 4°C. The following primary antibodies and dilutions were used: Histone 3 Rabbit mAb (Sigma Aldrich, #H0164; 1:10,000), GAPDH Mouse mAb (Sigma Aldrich, #G8795; 1:10000), JMJD1C Rabbit mAb (Abcam, #ab316320; 1:1000), H3K9me2 Mouse mAb (Abcam, #ab1220; 1:1000), α-SMA Mouse mAb (Sigma Aldrich, #A2547; 1:1000). Excess antibodies were removed by exchange with 3 washes of TBST after which membranes were incubated for 1 h in TBST containing secondary antibodies conjugated with either Alexa680 (Life Technologies) or IRDye 800 (Licor) diluted at 1:20,000. Secondary antibodies were removed by washing 3 times in TBST, after which immunoreactive bands were detected using an Odyssey CLx Infrared Imaging System (LI-COR Biosciences). Bands were analysed using Image Studio software (Licor) by creating a boundary around individual bands for quantification, performing background subtraction as determined for each lane individually, and calculating the signal/area within the boundary for each band. The signal/area of bands for the protein of interest to be quantified was divided by the signal/area of the housekeeping protein in the same lane.

**Immunofluorescence microscopy and image quantification**

FBs were seeded in µ-Slide 8 Well chambers (Ibidi, 80821). After treatment, cells were washed three times with PBS, fixed for 15 min at room temperature in 2% PFA in PBS, and washed three further times with PBS. Cells were permeabilised for 15 min with 0.2% Triton X-100 in PBS and blocked for 1 h in PBS containing 0.1% Triton X-100 and 3% bovine serum albumin (BSA). Primary antibodies against JMJD1C (Proteintech, 26637-1-AP; 1:1,000), H3K9me2 (Abcam, ab1220; 1:1,000), and α-smooth-muscle actin (α-SMA; Sigma–Aldrich, A2547; 1:500) were diluted in blocking buffer and incubated overnight at 4 °C. Cells were then washed four times for 15 min in PBS containing 0.1% Triton X-100 and incubated for 1 h with Alexa Fluor

488- and 568-conjugated secondary antibodies (Life Technologies) and, where indicated, phalloidin–iFluor 647 (Abcam, ab176759; 1:1,000) in blocking buffer. After four further 15-min washes, samples were mounted in Vectashield antifade mounting medium with DAPI (Vector Laboratories, H-1200-10) diluted 1:1 in PBS.

Fluorescence images were acquired on a Nikon A1R confocal microscope configured on an Eclipse Ti platform. JMJD1C imaging used a 40×/1.3-NA oil-immersion objective, whereas α-SMA imaging used a 20×/0.4-NA Ph2 objective. Six randomly selected fields were analysed per sample, with at least 100 cells evaluated in aggregate. Field-level measurements were averaged to obtain one descriptive value per biological replicate. Acquisition settings were held constant within each experiment. Representative channel and merged images were prepared only after quantification; the Fiji macros applied fixed display ranges within each experiment and added 100-μm scale bars.

For nuclear segmentation, the DAPI channel was smoothed with a Gaussian filter ($\sigma = 2.0$), thresholder with the Otsu dark-background method, converted to a binary mask, hole-filled, and separated by watershed[18]. Objects with an area of 30–1,500 μm² and circularity of 0.50–1.00 were retained as nuclear regions of interest (ROIs). Nuclear JMJD1C was quantified in each DAPI-defined ROI as mean fluorescence intensity. The quantification plotted individual-nucleus measurements together with biological-replicate means and analysed normalized nuclear JMJD1C intensity using a linear mixed-effects model with condition as a fixed effect and technical measurements nested within biological replicate as random effects, followed by a type III test and Tukey-adjusted pairwise comparisons.

For α-SMA quantification, the α-SMA channel underwent rolling-ball background subtraction with a radius of 50 pixels, Otsu thresholding, and binary conversion; the fraction of foreground pixels was recorded as the global α-SMA-positive area. To classify individual cells, each DAPI-defined nuclear ROI was enlarged by 12 pixels as an approximation of the perinuclear cell body, and mean α-SMA intensity was measured in the expanded ROI. Cells with a mean intensity of at least 400 arbitrary fluorescence units were classified as α-SMA positive, and the percentage of positive cells was calculated per field as 100 × positive cells/segmented nuclei. Field-level percentages were analysed with a linear mixed-effects model with condition as a fixed effect and technical measurements nested within biological replicate as random effects, followed by a type III test and Tukey-adjusted pairwise comparisons.

#### Bulk RNA-seq of FB perturbations

##### Transcript quantification and gene-level counts

Primary RNA-seq processing was performed with nf-core/rnaseq (v3.24.0) under Nextflow (v25.10.4)[19] using the Singularity[20] profile and the pipeline's GRCh38 reference. Paired-end reads were aligned with STAR[21] in two-pass mode, and transcriptome-coordinate BAM files were generated with --quantMode TranscriptomeSAM. Transcript abundance was then quantified in alignment-based mode with Salmon (v1.10.3)[22]. Salmon was run with six threads, the filtered reference annotation supplied through --geneMap genes.filtered.gtf, transcript sequences in genome.transcripts.fa, and the STAR Aligned.toTranscriptome.out.bam file as input and library type IU (paired-end, inward-facing, unstranded). No --seqBias, --gcBias or --validateMappings options were supplied in the executed command.

StringTie (v2.2.3)[23] was run separately on each coordinate-sorted, duplicate-marked BAM using genes.filtered.gtf as the reference annotation, and the options -e, -A, -C and -b; thus, expression estimation was restricted to transcripts in the supplied annotation while gene-abundance, covered-transcript and Ballgown-format outputs were generated. For downstream gene-level analysis, a transcript-to-gene map was constructed from the union StringTie annotation generated by the analysis workflow, and Salmon transcript estimates were imported into R with tximport (v1.30.0)[24] using countsFromAbundance = "no". The downstream reports were generated in R 4.3.2.

##### Differential expression (DE) analysis

Gene-level DE was analysed with DESeq2 (v1.42.1)[25]. Genes with at least 10 counts in at least 5 samples were retained. Variance-stabilising transformation used blind = FALSE for principal-component analysis (PCA) and heat maps. For coefficient-based contrasts, $\log_2$FC was shrunk with apeglm (v1.24.0)[26]; composite contrasts used maximum-likelihood estimates. Benjamini–Hochberg adjusted $P < 0.05$ and $|\log_2\mathrm{FC}| \geq 0.58$ were used for the current main DE display.

##### Functional enrichment of bulk RNA-seq results

GOBP over-representation analysis used clusterProfiler (v4.10.1)[27] and org.Hs.eg.db (v3.18.0) with all tested genes as the background universe, gene-set sizes of 10–500 and Benjamini–Hochberg adjusted $P < 0.05$. Preranked gene-set enrichment used the DESeq2 Wald statistic, seed-controlled analysis, minimum and maximum gene-set sizes of 10 and 1,000 and Benjamini–Hochberg correction[28]. Sample-level pathway scores were calculated from variance-stabilised expression with GSVA using a Gaussian kernel and gene-set sizes of 10–1,000[29]. Differential pathway activity was tested with limma using the design ~ batch +

knockdown * treatment and empirical Bayes moderation with trend = TRUE[30]. Gene sets were obtained with msigdbr (v25.1.1) where indicated.

#### References


1. Zheng, G. X. Y. *et al.* Massively parallel digital transcriptional profiling of single cells. *Nat Commun* **8**, 14049 (2017).
2. Satpathy, A. T. *et al.* Massively parallel single-cell chromatin landscapes of human immune cell development and intratumoral T cell exhaustion. *Nat Biotechnol* **37**, 925–936 (2019).
3. Fleming, S. J. *et al.* Unsupervised removal of systematic background noise from droplet-based single-cell experiments using CellBender. *Nat Methods* **20**, 1323–1335 (2023).
4. Huang, X. & Huang, Y. Cellsnp-lite: an efficient tool for genotyping single cells. *Bioinformatics* **37**, 4569–4571 (2021).
5. Huang, Y., McCarthy, D. J. & Stegle, O. Vireo: Bayesian demultiplexing of pooled single-cell RNA-seq data without genotype reference. *Genome Biology* **20**, 273 (2019).
6. Wolf, F. A., Angerer, P. & Theis, F. J. SCANPY: large-scale single-cell gene expression data analysis. *Genome Biol* **19**, 15 (2018).
7. Kanemaru, K. *et al.* Spatially resolved multiomics of human cardiac niches. *Nature* **619**, 801–810 (2023).
8. Wolock, S. L., Lopez, R. & Klein, A. M. Scrublet: Computational Identification of Cell Doublets in Single-Cell Transcriptomic Data. *Cell Systems* **8**, 281-291.e9 (2019).
9. Hao, Y. *et al.* Integrated analysis of multimodal single-cell data. *Cell* **184**, 3573-3587.e29 (2021).
10. Finak, G. *et al.* MAST: a flexible statistical framework for assessing transcriptional changes and characterizing heterogeneity in single-cell RNA sequencing data. *Genome Biol* **16**, 278 (2015).
11. Kuleshov, M. V. *et al.* Enrichr: a comprehensive gene set enrichment analysis web server 2016 update. *Nucleic Acids Res* **44**, W90–W97 (2016).
12. Stuart, T., Srivastava, A., Madad, S., Lareau, C. A. & Satija, R. Single-cell chromatin state analysis with Signac. *Nat Methods* **18**, 1333–1341 (2021).
13. Lopez, R., Regier, J., Cole, M. B., Jordan, M. I. & Yosef, N. Deep generative modeling for single-cell transcriptomics. *Nat Methods* **15**, 1053–1058 (2018).
14. Ashuach, T., Reidenbach, D. A., Gayoso, A. & Yosef, N. PeakVI: A deep generative model for single-cell chromatin accessibility analysis. *Cell Reports Methods* **2**, 100182 (2022).
15. Kamimoto, K. *et al.* Dissecting cell identity via network inference and in silico gene perturbation. *Nature* **614**, 742–751 (2023).
16. Cools, J. M. T. *et al.* Small-molecule-induced ERBB4 activation to treat heart failure. *Nat Commun* **16**, 576 (2025).

17. Schroeder, A. B. *et al.* The ImageJ ecosystem: Open-source software for image visualization, processing, and analysis. *Protein Science* **30**, 234–249 (2021).
18. Otsu, N. A Threshold Selection Method from Gray-Level Histograms. *IEEE Transactions on Systems, Man, and Cybernetics* **9**, 62–66 (1979).
19. Langer, B. E. *et al.* Empowering bioinformatics communities with Nextflow and nf-core. *Genome Biol* **26**, 228 (2025).
20. Kurtzer, G. M., Sochat, V. & Bauer, M. W. Singularity: Scientific containers for mobility of compute. *PLOS ONE* **12**, e0177459 (2017).
21. Dobin, A. *et al.* STAR: ultrafast universal RNA-seq aligner. *Bioinformatics* **29**, 15–21 (2013).
22. Patro, R., Duggal, G., Love, M. I., Irizarry, R. A. & Kingsford, C. Salmon provides fast and bias-aware quantification of transcript expression. *Nat Methods* **14**, 417–419 (2017).
23. Pertea, M. *et al.* StringTie enables improved reconstruction of a transcriptome from RNA-seq reads. *Nat Biotechnol* **33**, 290–295 (2015).
24. Soneson, C., Love, M. I. & Robinson, M. D. Differential analyses for RNA-seq: transcript-level estimates improve gene-level inferences. Preprint at https://doi.org/10.12688/f1000research.7563.2 (2016).
25. Love, M. I., Huber, W. & Anders, S. Moderated estimation of fold change and dispersion for RNA-seq data with DESeq2. *Genome Biology* **15**, 550 (2014).
26. Zhu, A., Ibrahim, J. G. & Love, M. I. Heavy-tailed prior distributions for sequence count data: removing the noise and preserving large differences. *Bioinformatics* **35**, 2084–2092 (2019).
27. Wu, T. *et al.* clusterProfiler 4.0: A universal enrichment tool for interpreting omics data. *The Innovation* **2**, 100141 (2021).
28. Subramanian, A. *et al.* Gene set enrichment analysis: A knowledge-based approach for interpreting genome-wide expression profiles. *Proceedings of the National Academy of Sciences* **102**, 15545–15550 (2005).
29. Hänzelmann, S., Castelo, R. & Guinney, J. GSVA: gene set variation analysis for microarray and RNA-Seq data. *BMC Bioinformatics* **14**, 7 (2013).
30. Ritchie, M. E. *et al.* limma powers differential expression analyses for RNA-sequencing and microarray studies. *Nucleic Acids Res* **43**, e47 (2015).

#### Supplementary figures

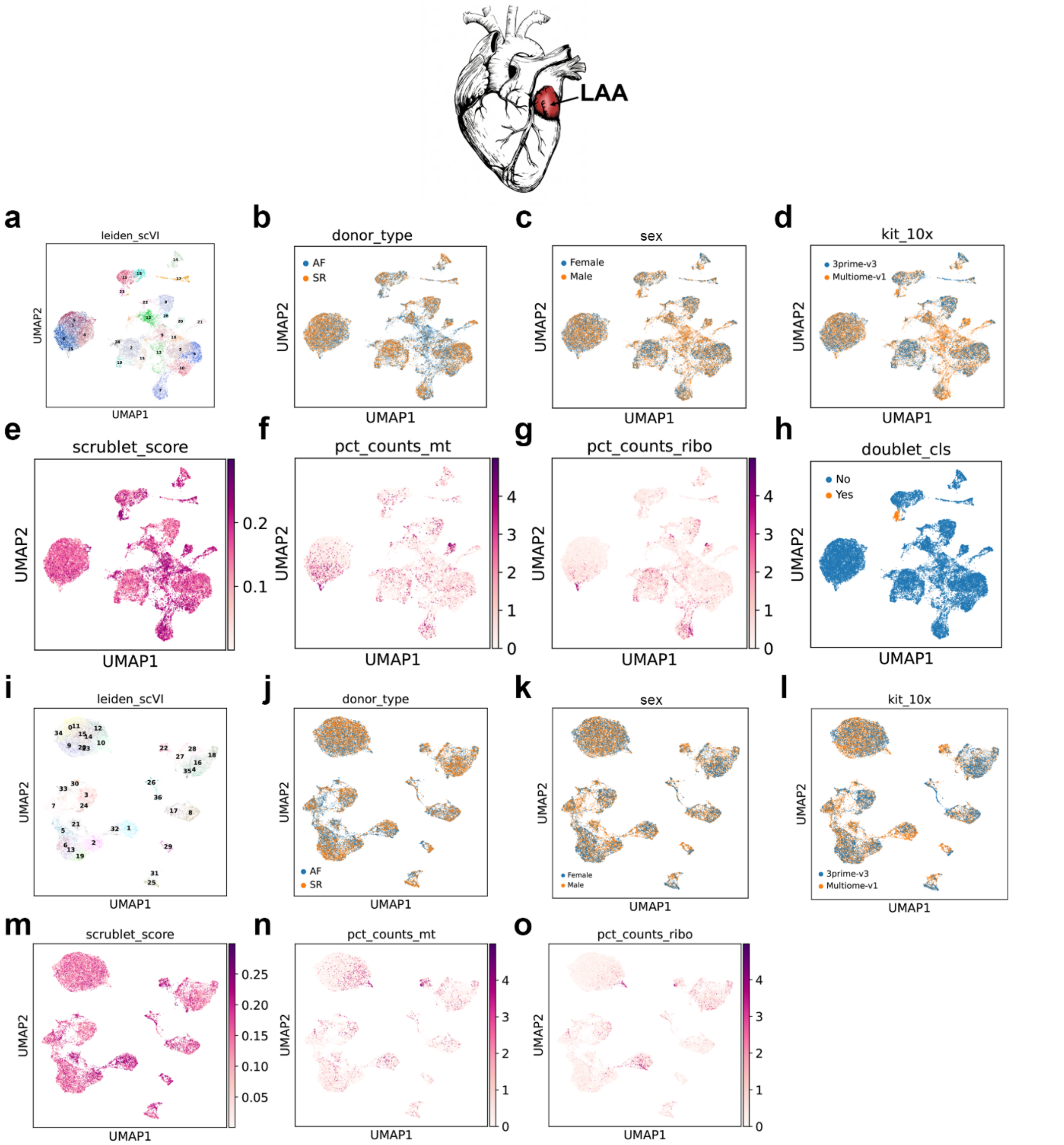


**Supplementary Fig. 1 | Quality control (QC) of the left atrial appendage (LAA) single-nucleus RNA-sequencing (snRNA-seq) dataset. a–h**, Pre-filtering UMAPs and quality metrics showing initial clusters, donor rhythm, sex, 10x chemistry, Scrublet doublet score, mitochondrial-read fraction, ribosomal-read fraction, and cluster of doublets. **i–o**, Corresponding post-filtering embeddings and covariates for retained nuclei. Nuclei with fewer than 200 detected genes or 500 counts, nuclei exceeding the applied mitochondrial and ribosomal-read threshold (5%), Scrublet-predicted doublets, and Vireo doublet assignments were removed. UMAP, uniform manifold approximation and projection. For details on filtering, see Methods.

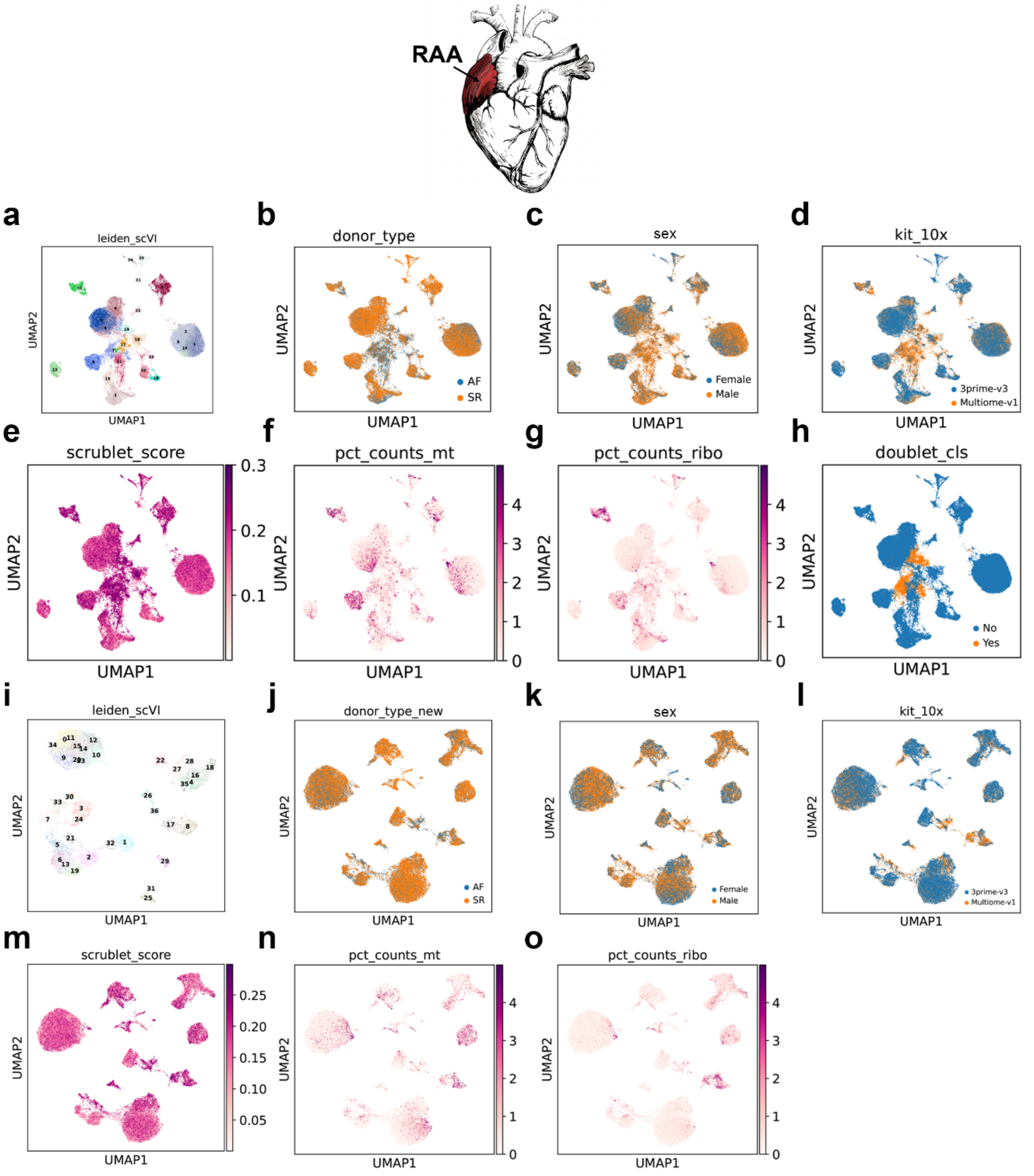


**Supplementary Fig. 2 | QC of the right atrial appendage (RAA) snRNA-seq dataset. a–h**, Pre-filtering UMAPs and quality metrics showing initial clusters, donor rhythm, sex, 10x chemistry, Scrublet doublet score, mitochondrial-read fraction, ribosomal-read fraction, and cluster of doublets. **i–o**, Corresponding post-filtering embedding and covariates for retained nuclei. UMAP, uniform manifold approximation and projection. For details on filtering, see Methods.

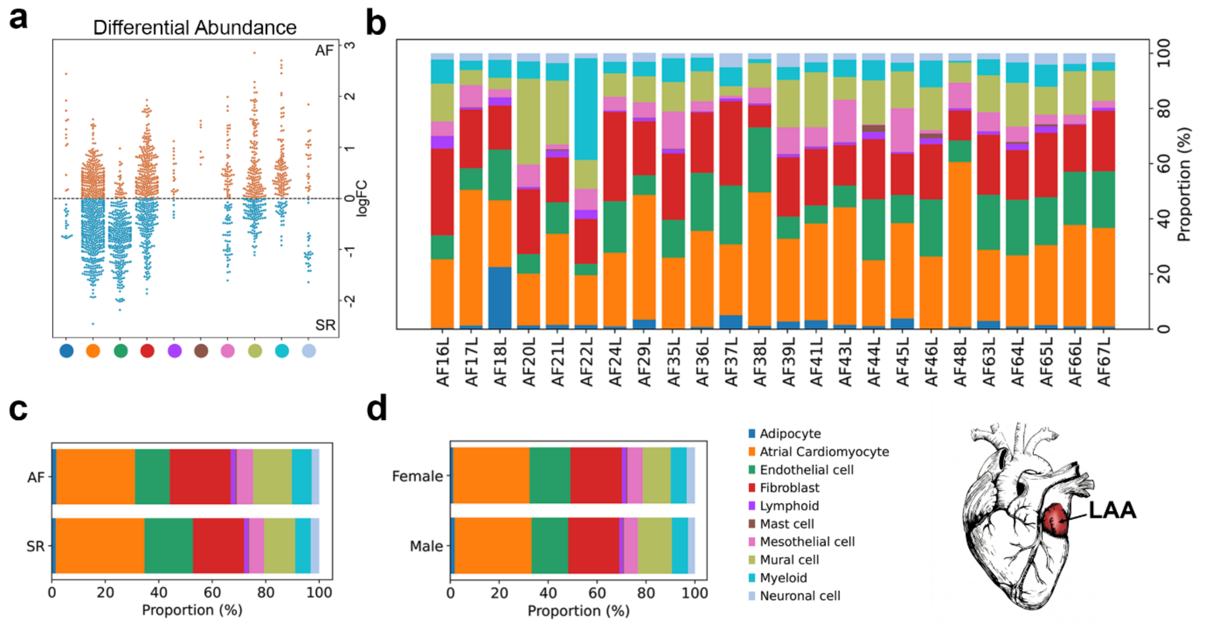


**Supplementary Fig. 3 | Cell-type composition of the LAA atlas. a**, Distribution of Milo log fold changes across neighbourhoods assigned to each LAA cell type. **b**, Cell-type composition for each donor. **c**,**d**, Aggregate composition by rhythm (**c**) and sex (**d**).

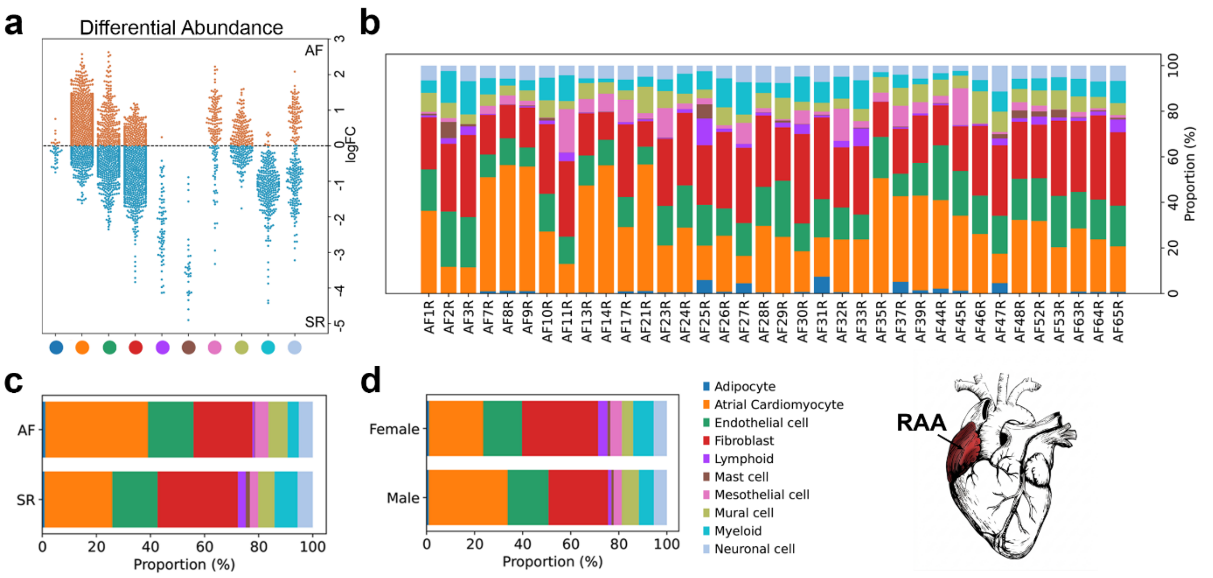


**Supplementary Fig. 4 | Cell-type composition of the RAA atlas. a**, Distribution of Milo log fold changes across neighbourhoods assigned to each RAA cell type. **b**, Per-donor cell-type composition. **c**,**d**, Aggregate composition by rhythm (**c**) and sex (**d**).

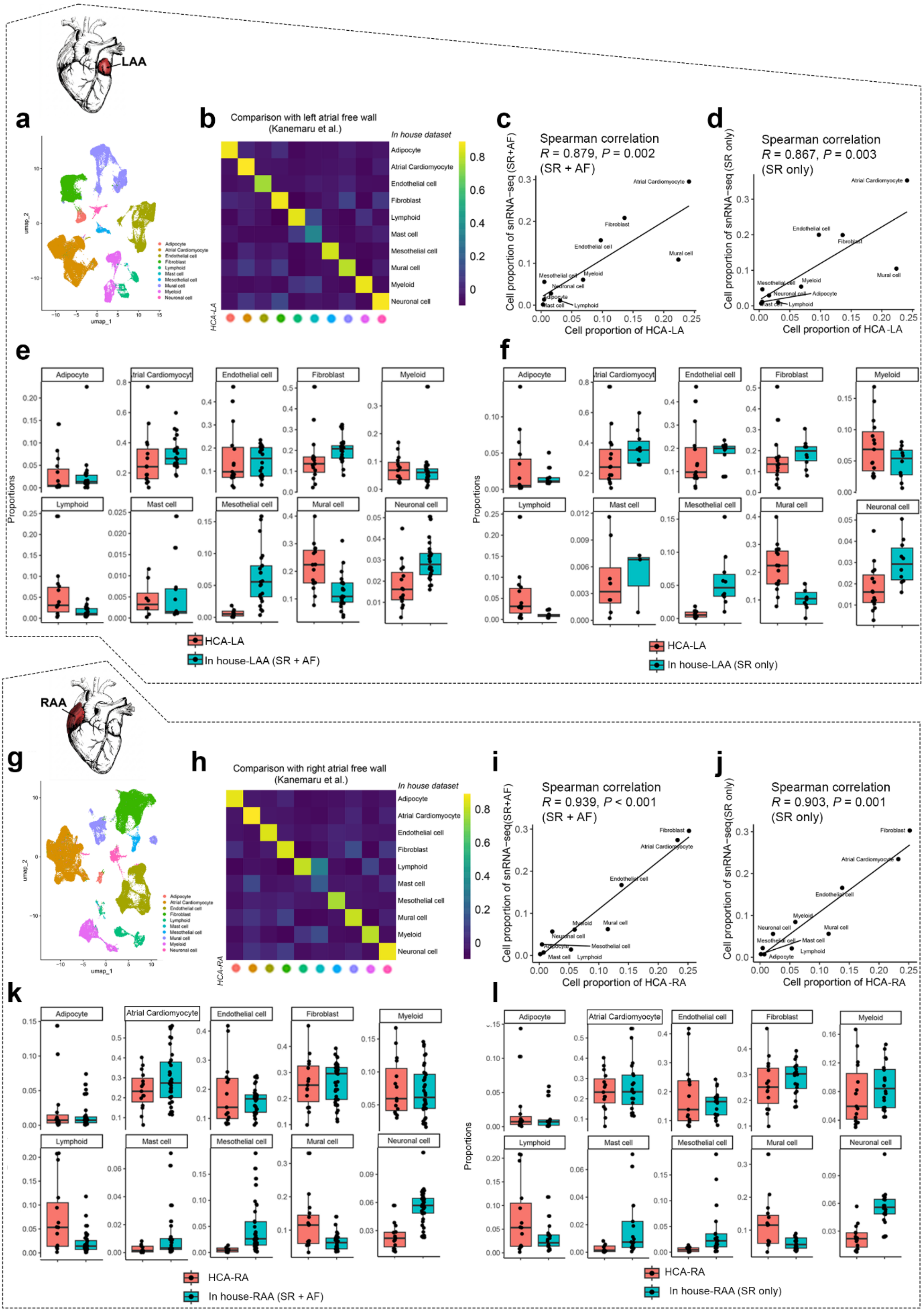


**Supplementary Fig. 5 | Concordance of the atrial appendage atlases with reference atrial free-wall populations. a**, UMAP of left-atrial nuclei from Human Cardiac Cell Atlas (HCA-LA) by Kanemaru et al[7], coloured by major cell identity. **b**, Cell-type similarity matrix comparing the in-house LAA atlas with HCA-LA. **c,d**, Spearman correlations of cell-type proportions between HCA-LA and the in-house LAA atlas using AF and SR samples together (**c**) or SR samples only (**d**). **e,f**, Donor-level cell-type proportions in HCA-LA and the corresponding in-house LAA datasets using AF and SR samples together (**e**) or SR samples only (**f**). **g–l**, Corresponding comparison of the in-house RAA atlas with right-atrial free-wall populations (HCA-RA) from Kanemaru *et al*. In c,d,i,j, each point represents one major cell type; Spearman correlation coefficients and two-sided *P* values are shown. In e,f,k,l, points represent individual samples, centre lines indicate medians, boxes indicate interquartile ranges, and whiskers extend to 1.5 times the interquartile range. AF, atrial fibrillation; SR, sinus rhythm; HCA, Human Cardiac Atlas.

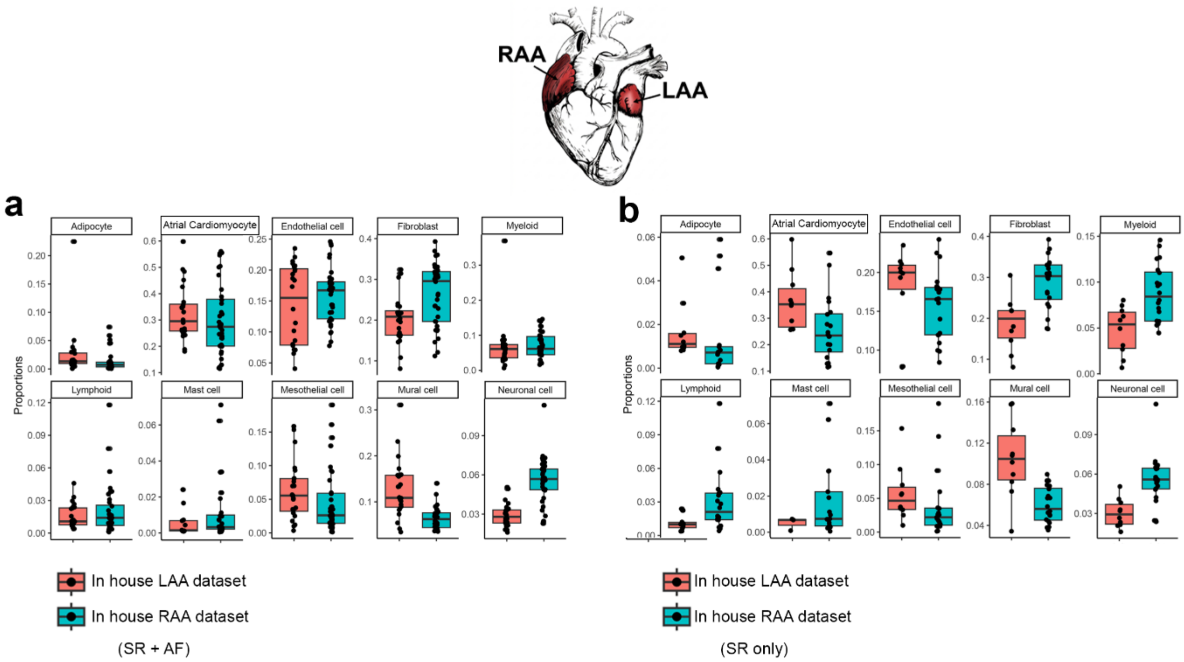


**Supplementary Fig. 6 | Integrated comparison of the human RAA and LAA atlases. a**,**b**,

Donor-level cell-type proportions in LAA and RAA using AF and SR samples (**a**) or SR samples only (**b**).

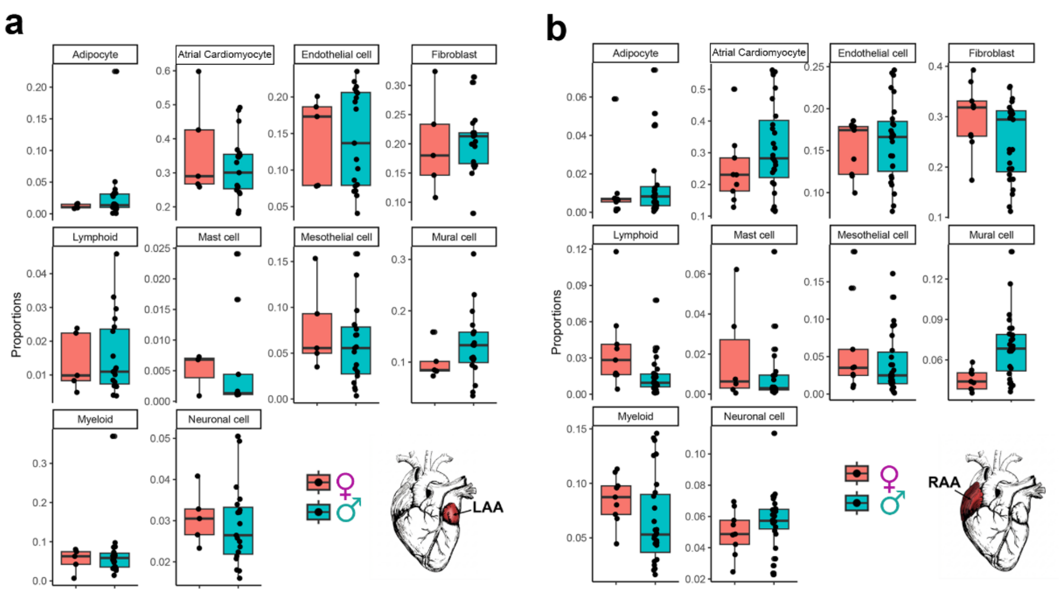


**Supplementary Fig. 7 | Sex-stratified analyses of cell composition. a**,**b**, Donor-level cell-type proportion between male and female in LAA (**a**) and RAA (**b**).

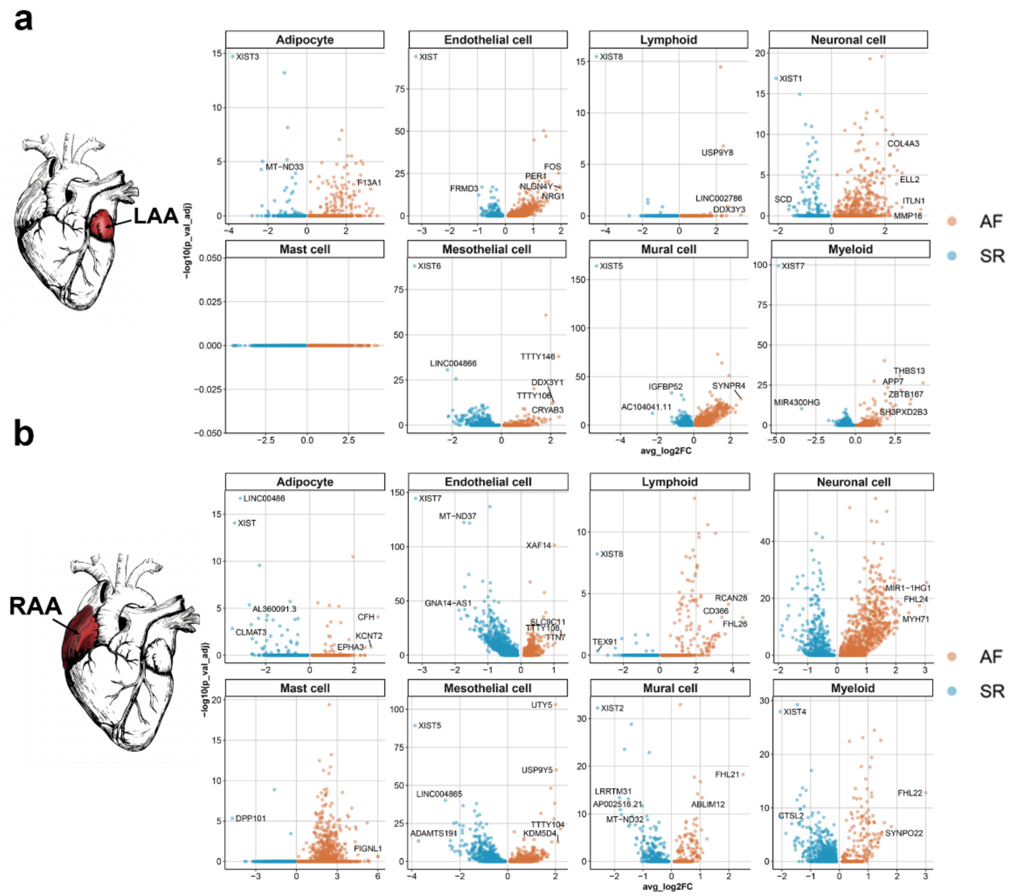


**Supplementary Fig. 8 | Cell-type-specific AF transcriptional signatures. a**,**b**, Volcano plots of AF versus SR differential expression (DE) within each major cell type in LAA (**a**) and RAA (**b**). Positive fold changes in orange denote higher expression in AF; negative fold changes in blue denote higher expression in SR. Significance is set at $P$ adj < 0.05 and |log2 fold change| ≥ 0; up to five selected genes are labelled in each comparison. AF, atrial fibrillation; SR, sinus rhythm.

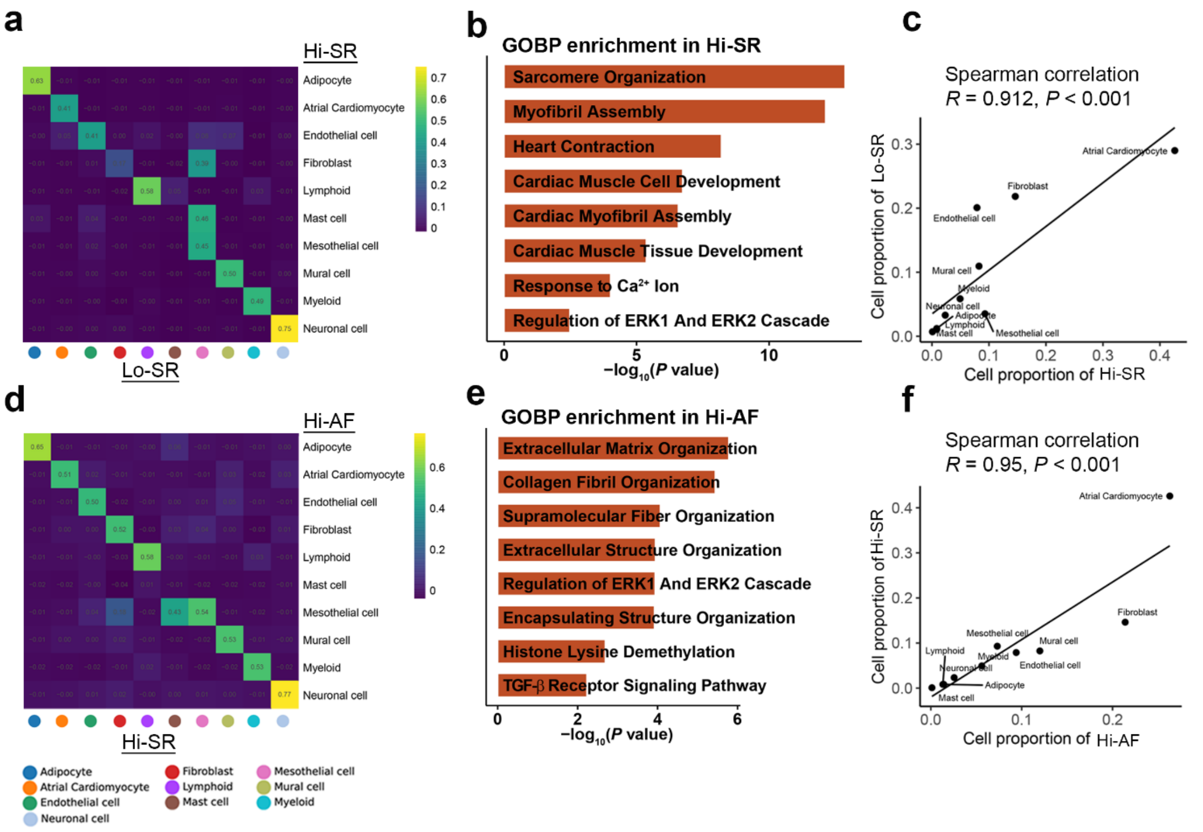


**Supplementary Fig. 9 | Pairwise analyses separate rhythm- and structural remodelling-associated atrial signatures. a–c**, Hi-SR versus Lo-SR. **d–f**, Hi-AF versus Hi-SR. For each comparison, cell-type similarity is followed by selected enriched biological process (BP) and concordance of donor-group cell-type proportions. In c,f, Exact correlation coefficients and *P* values are shown in each panel.

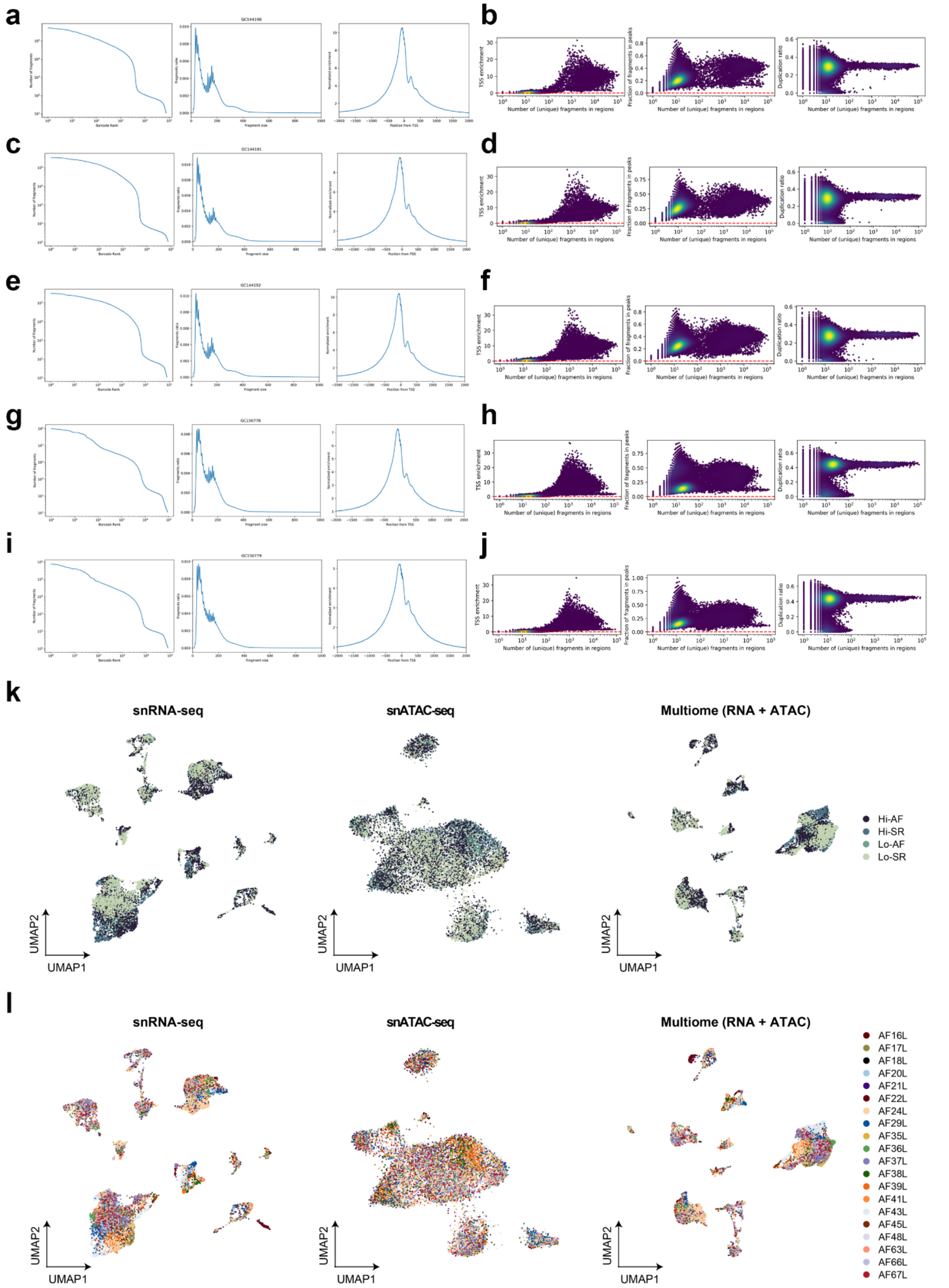


**Supplementary Fig. 10 | QC of the paired LAA Multiome dataset. a,c,e,g,i**, Barcode rank curves (left), fragment length distributions (middle) and transcription start site (TSS) enrichment profiles (right) for five Multiome libraries (before demultiplexing). **b**,**d**,**f**,**h**,**j**, Per-nucleus relationships among TSS enrichment (left), fraction of fragments in peaks (middle) and duplication ratio (right) for the corresponding libraries. **k**,**l**, RNA (left), ATAC (middle) and integrated UMAPs (right) coloured by group (**k**) or donor (**l**). Hi-AF, AF with high left atrial volume index (LAVI); Hi-SR, sinus rhythm with high LAVI; Lo-AF, AF with low LAVI; Lo-SR, SR with low LAVI.

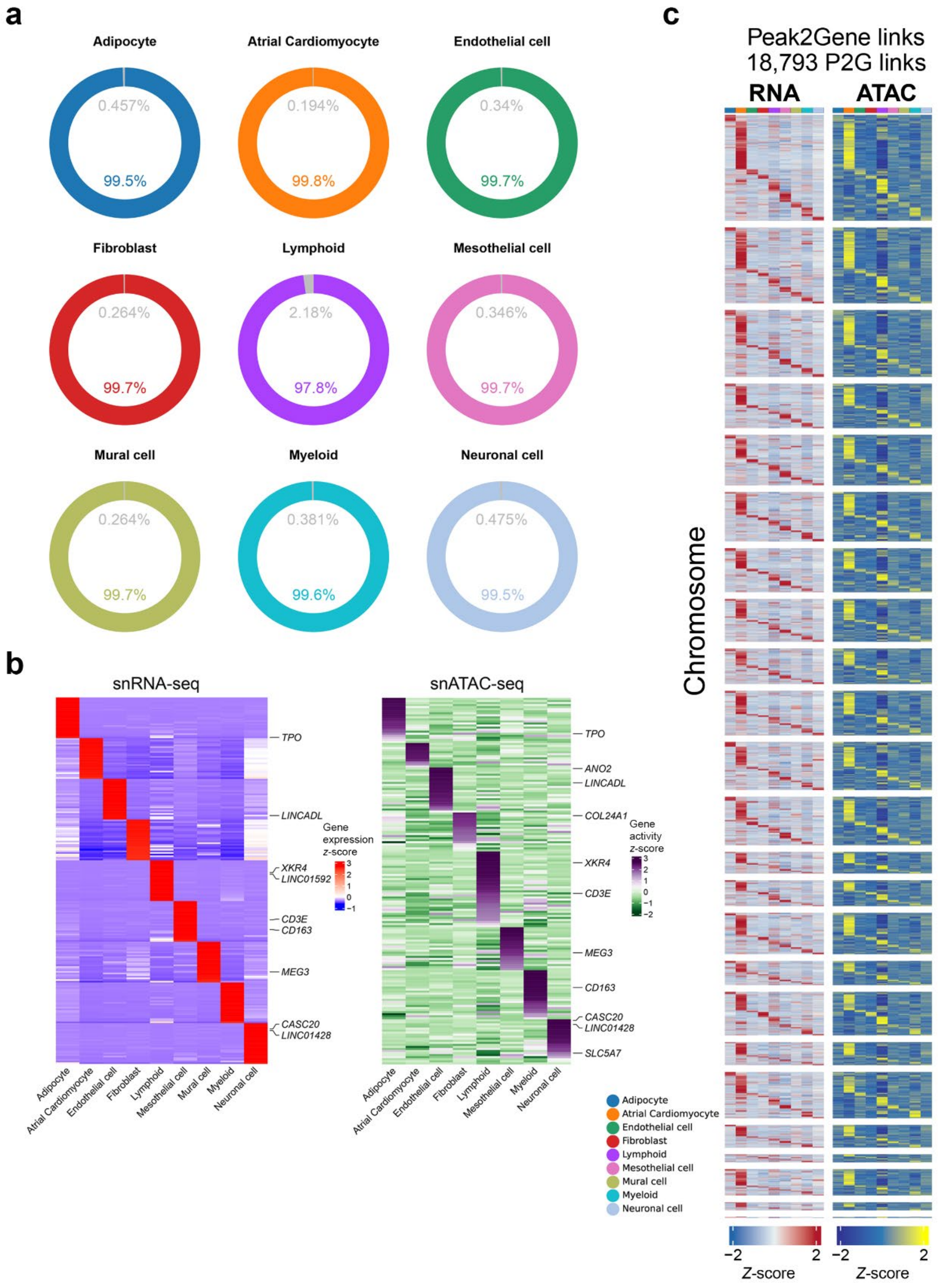


**Supplementary Fig. 11 | Cross-modality validation of Multiome cell identities and peak-to-gene links. a**, Donut chart showing the percent of genes with high chromatin accessibility and low gene expression in grey for each major cell type. High chromatin accessibility was defined as genes in the top 20% of gene activity, while low gene expression was defined as genes in the bottom 20% of gene expression. Percent of all other genes are colored by the cell type. **b**, Row-normalized RNA expression (left) and ATAC-derived gene activity (right) for the 25 most upregulated genes in each cell type, ranked by average log fold change. **c**, Chromosome-stratified RNA and ATAC heatmaps for 18,793 peak-to-gene (P2G) links.

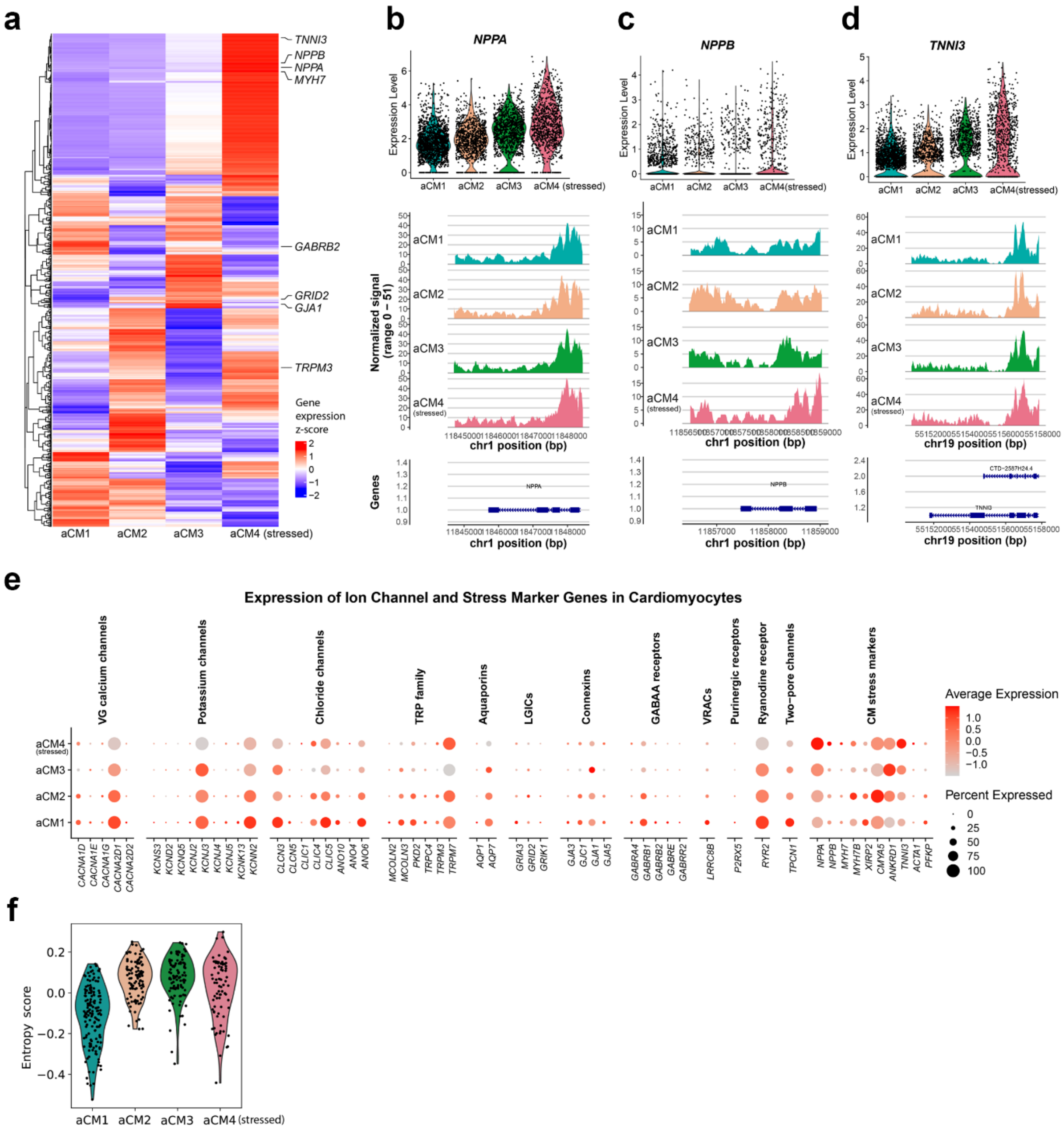


**Supplementary Fig. 12 | Molecular features of atrial cardiomyocyte (aCM) states. a**, Heatmap of genes differentially expressed among aCM1–aCM4. **b–d**, Violin plots of *NPPA*, *NPPB* and *TNNI3* expression across states (top) with corresponding aggregate chromatin-accessibility tracks (bottom). **e**, Dot plots of calcium-handling, ion-channel and stress-response genes; colour denotes scaled mean expression and point size the expressing fraction. **f**, Adjusted chromatin-entropy score across aCM states.

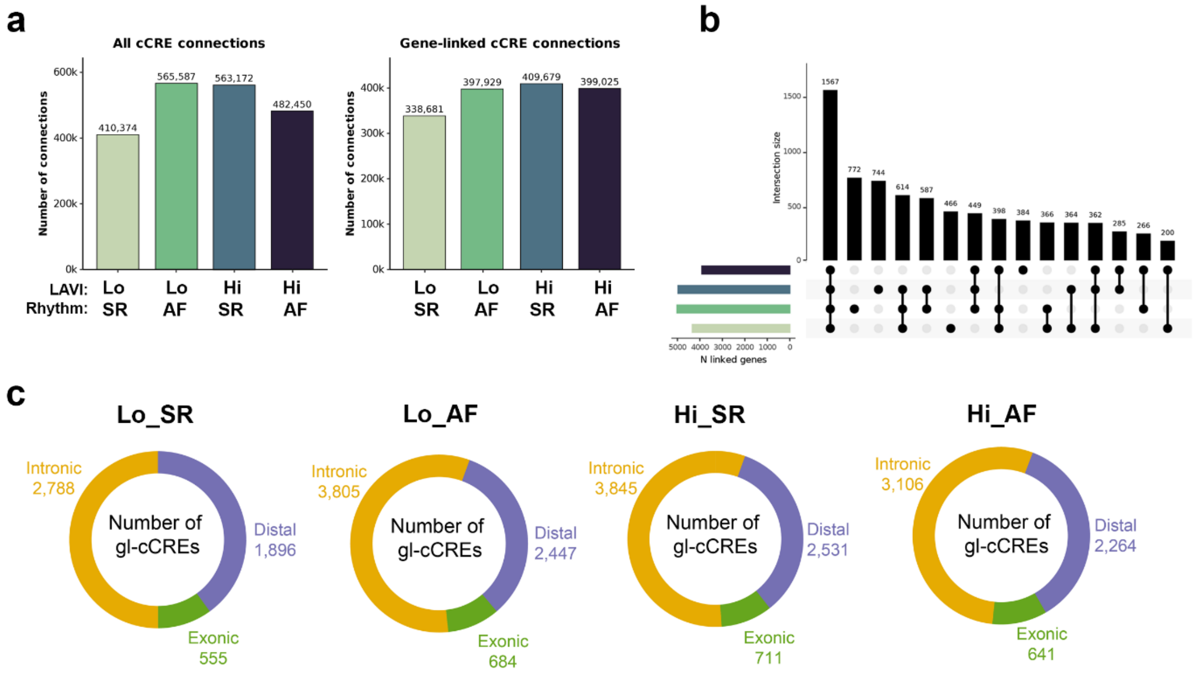


**Supplementary Fig. 13 | *Cis*-regulatory connectivity differs between aCM groups. a**, Total candidate *cis*-regulatory element (cCRE) connections (left) and gene-linked cCRE (gl-cCRE)

connections (right) in each group. **b**, UpSet plot showing group-specific and shared linked-gene sets. **c**, Genomic annotation of gl-cCRE as intronic, exonic or distal for each group.

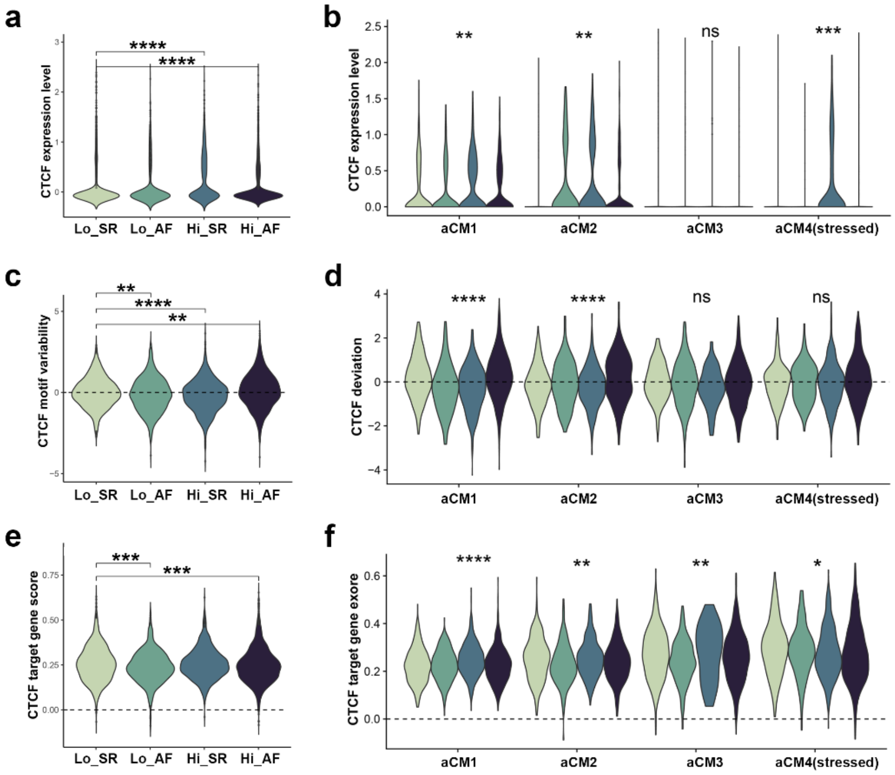


**Supplementary Fig. 14 | *CTCF*-associated activity varies across groups and CM states.** **a**,**c**,**e**, *CTCF* expression (**a**), *CTCF* motif variability (**c**), and *CTCF* target-gene score (**e**) across rhythm-LAVI groups. **b**,**d**,**f**, Corresponding measurements across aCM1–aCM4, stratified by rhythm-LAVI group. Violins show nucleus-level distributions; a two-sided Wilcoxon test was used; ns, not significant; *$P \leq 0.05$, **$P \leq 0.01$, ***$P \leq 0.001$, ****$P \leq 0.0001$.

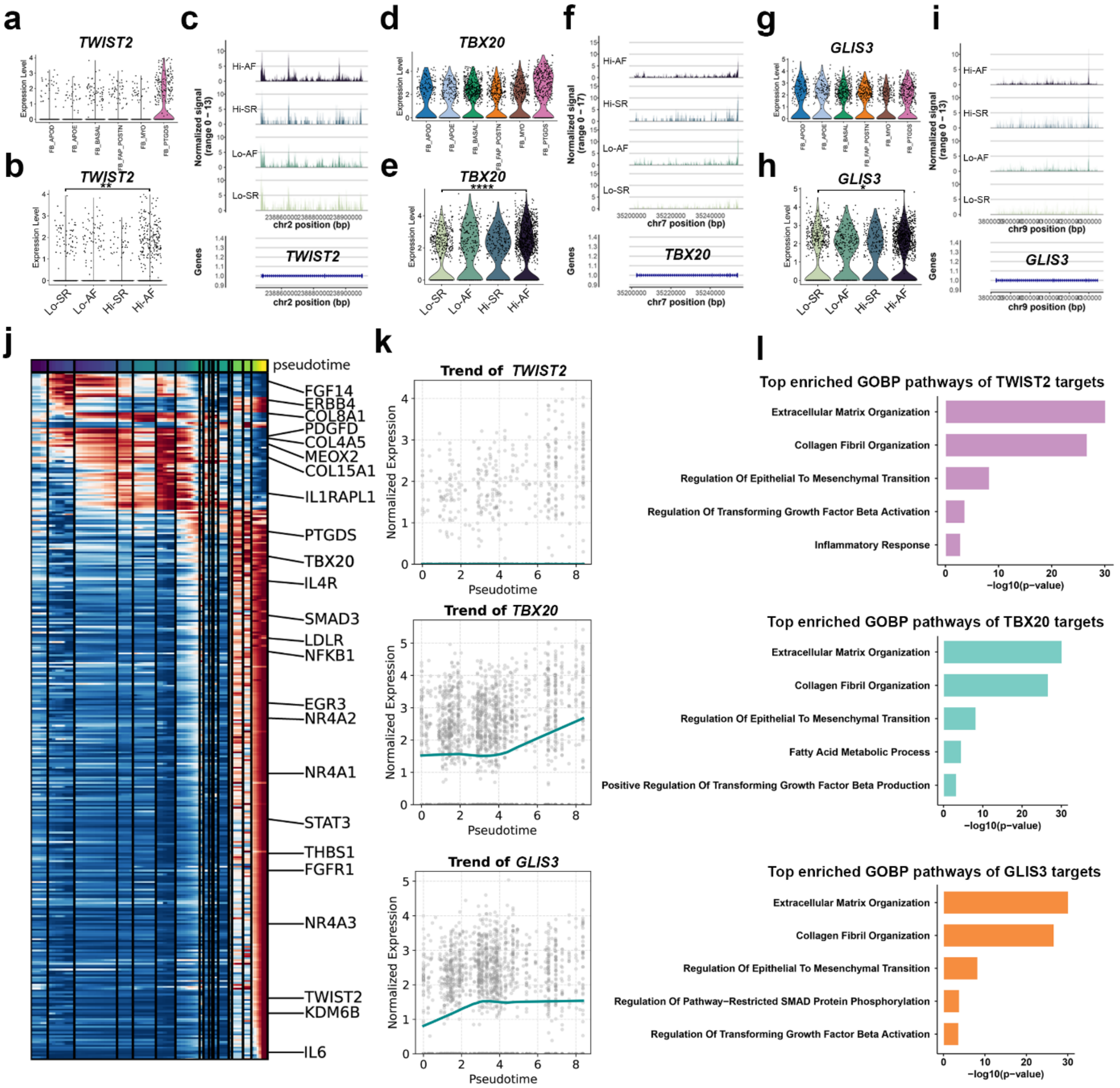


**Supplementary Fig. 15 | Fibroblast (FB) regulons and transcriptional dynamics along pseudotime. a–c**, *TWIST2* expression across FB states (**a**) and rhythm-LAVI groups (**b**), with aggregate accessibility tracks by group (**c**). **d–f**, Corresponding analyses for *TBX20*. **g–i**, Corresponding analyses for *GLIS3*. **j**, Heatmap of top differentially expressed genes (DEGs) varying along the basal-to-PTGDS$^{+}$ FB trajectory. **k**, Smoothed *TWIST2*, *TBX20* and *GLIS3* expression along scFates pseudotime. **l**, Selected GOBP terms enriched among SCENIC+-predicted targets of each TF, *TWIST2* (top), *TBX20* (middle), *GLIS3* (bottom). A two-sided

Wilcoxon test was used; *$P \leq 0.05$, **$P \leq 0.01$, ***$P \leq 0.001$, ****$P \leq 0.0001$. GOBP, Gene Ontology Biological Process.

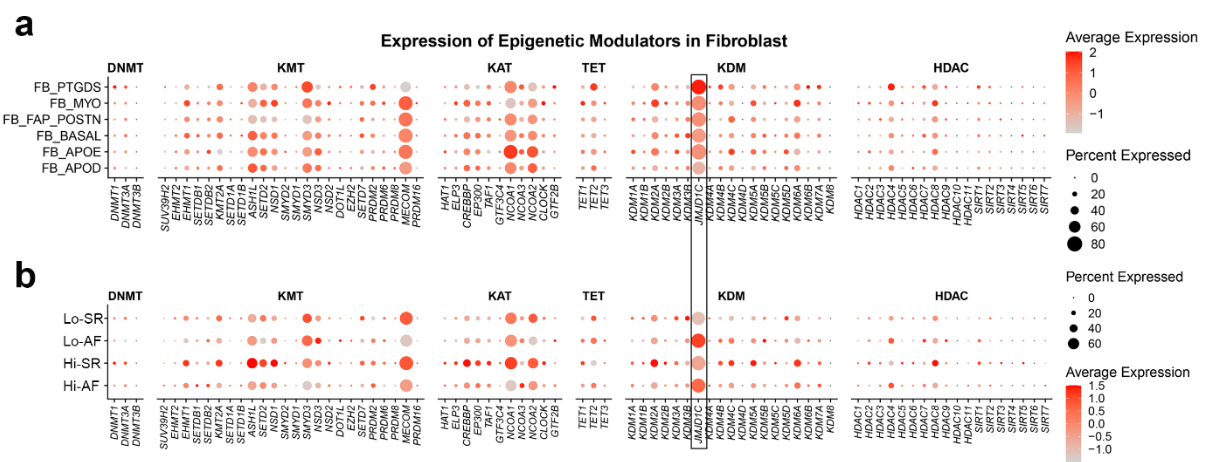


**Supplementary Fig. 16 | *JMJD1C* is prominent among FB epigenetic regulators. a**,**b**, Expression of DNA methyltransferases (DMT), lysine methyltransferases (KMT), lysine acetyltransferases (KAT), TET enzymes, lysine demethylases (KDM) and histone deacetylases (HDAC) across FB states (**a**) and rhythm-LAVI groups (**b**). *JMJD1C* is boxed. Colour denotes scaled mean expression and point size denotes the percentage of expressing nuclei.

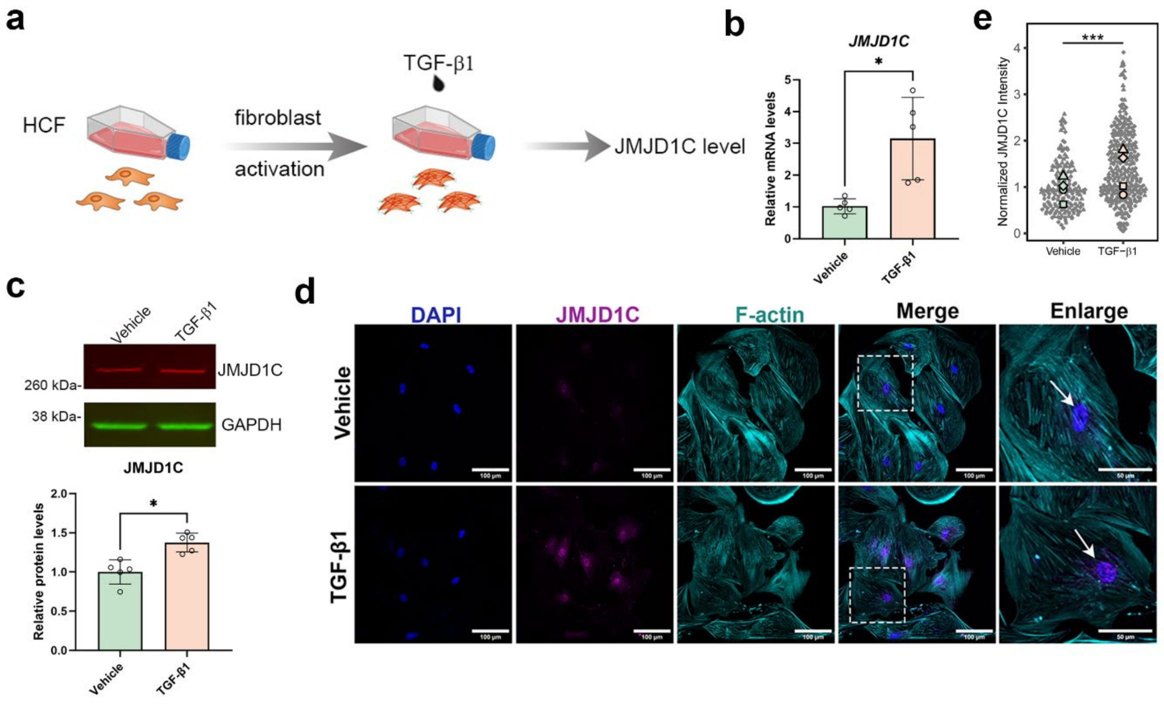


**Supplementary Fig. 17 | TGF-β1 increases *JMJD1C* expression in human cardiac fibroblasts (HCFs). a**, Experimental design. Primary HCFs were treated with vehicle or TGF-β1 (50 ng/mL, 48 h). **b**,**c**, *JMJD1C* transcript abundance measured by quantitative PCR (**b**)

and protein abundance measured by immunoblotting with GAPDH as the loading control (**c**). Points represent independent experiments ($n = 5$); bars show mean ± s.d. **d**, Representative immunofluorescence images showing DAPI (blue), JMJD1C (magenta), and F-actin (cyan); arrows indicate nuclear JMJD1C signal. Scale bars, 100 µm; enlargements, 50 µm. **e**, Swarm plot of quantification of normalized nuclear JMJD1C intensity. Small grey symbols represent individual cells and larger coloured symbols represent experiment-level means. Cells were nested within independent experiments and analysed using a linear mixed-effects model with treatment as a fixed effect and experiment as a random effect, followed by Tukey-adjusted comparisons. *$P < 0.05$; ***$P < 0.001$.

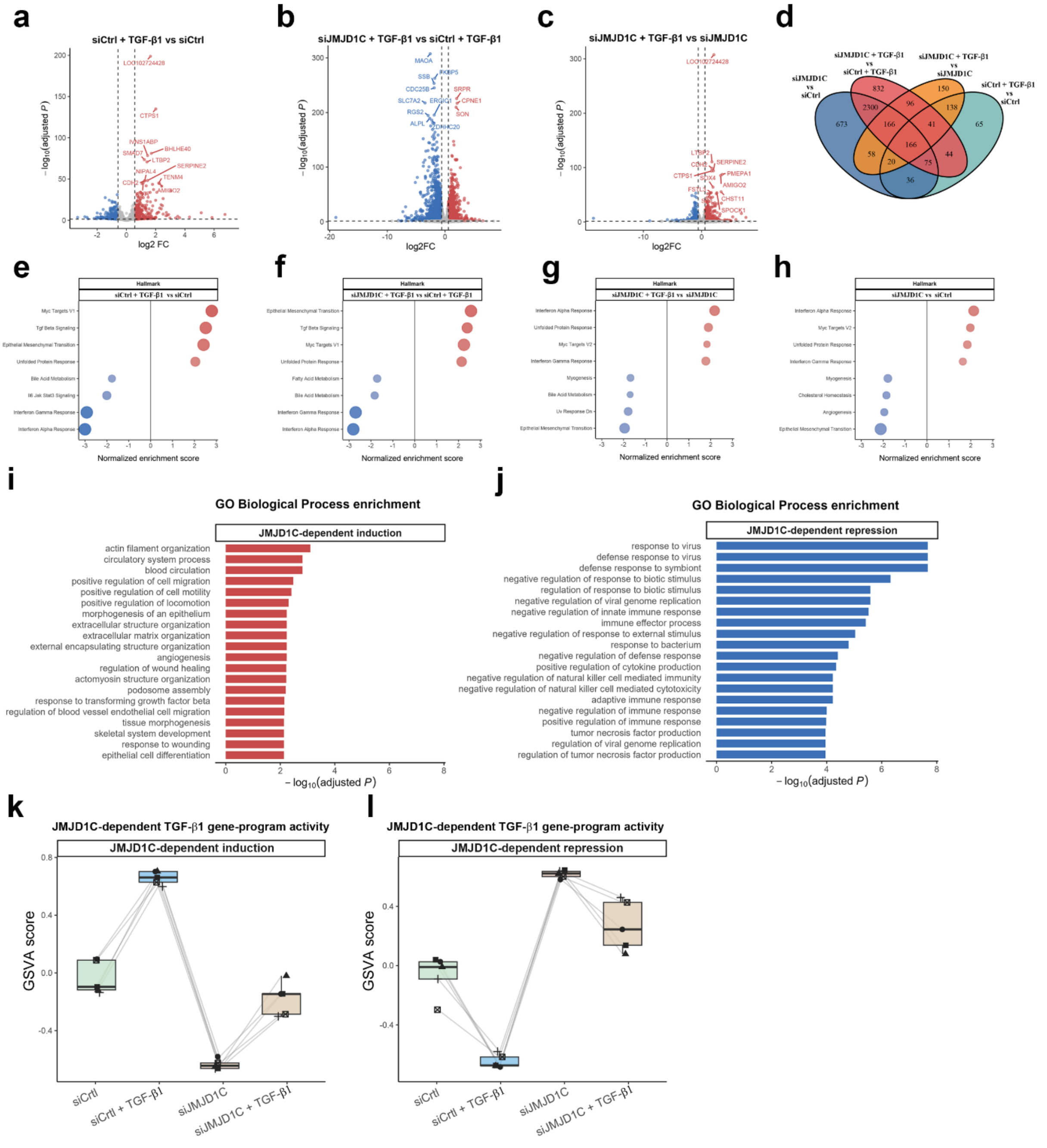


**Supplementary Fig. 18 | *JMJD1C* knockdown reshapes TGF-β1-responsive transcriptional programmes. a–c**, DESeq2 volcano plots for TGF-β1 versus vehicle in control cells (**a**), siJMJD1C versus control under TGF-β1 (**b**), and TGF-β1 versus vehicle after JMJD1C knockdown (**c**). Significant genes have adjusted $P$ < 0.05 and |log2 fold change| ≥ 0.58. **d**, Venn diagram of overlap of the corresponding DEG sets. **e–h**, Hallmark gene-set enrichment analysis (GSEA) for the indicated contrasts; red and blue denote positive and negative normalized enrichment scores. **i**,**j**, GOBP enrichment among *JMJD1C*-dependent induced (**i**) and repressed (**j**) genes. **k**,**l**, Gene-set variation analysis (GSVA) scores for these programmes in i and j across four conditions, respectively. DEGs, differentially expressed genes. GOBP, Gene Ontology Biological Process.

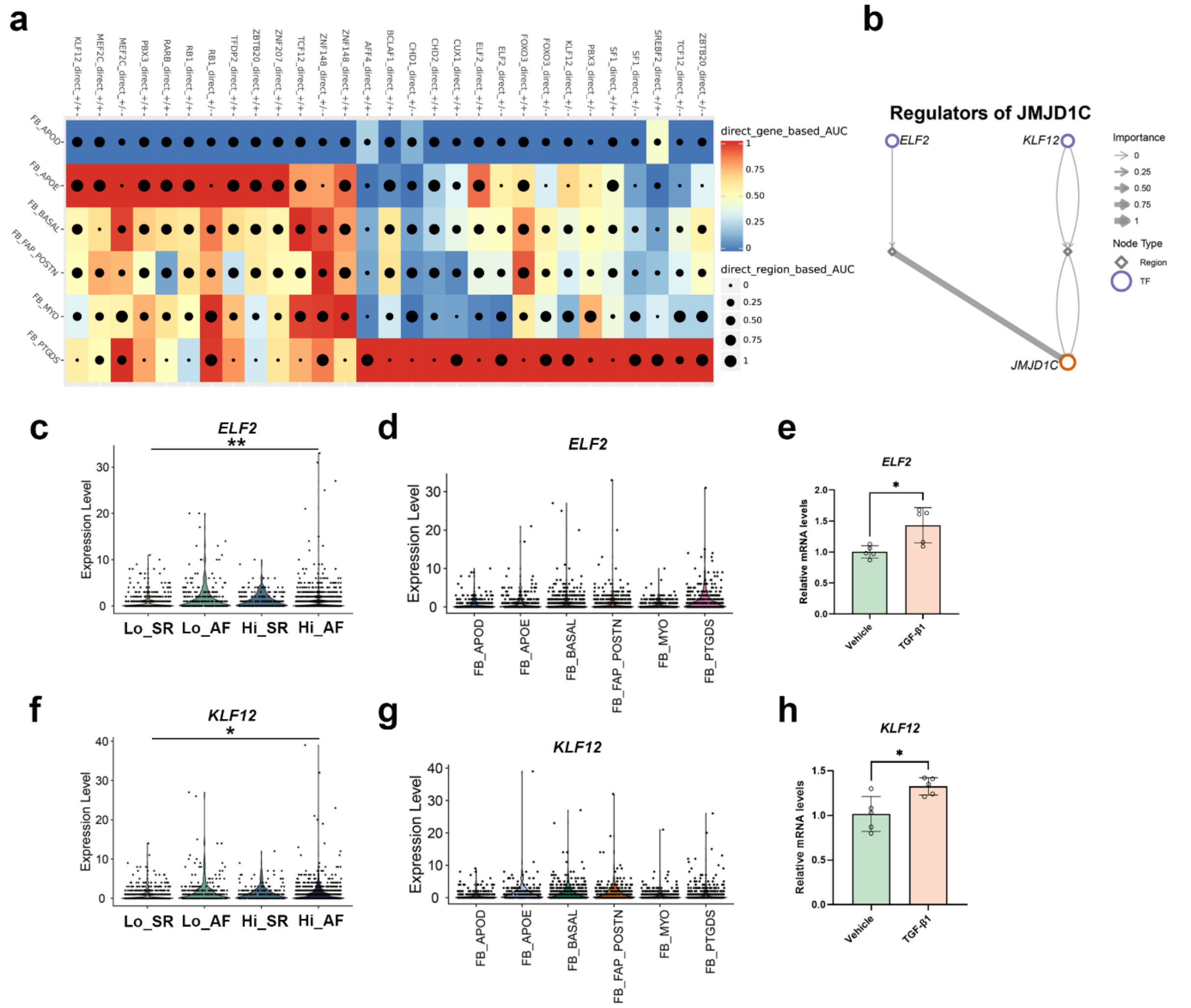


**Supplementary Fig. 19 | *ELF2* and *KLF12* are candidate upstream regulators of *JMJD1C*.** **a**, FB eRegulon activity (computed with AUCell based on target gene expression and target region accessibility) for candidate regulators across FB states; colour denotes gene-based AUCell score and point size denotes region-based AUCell score. **b**, SCENIC+-inferred *ELF2*- and *KLF12*-to-JMJD1C regulatory network. **c**,**d**, *ELF2* expression across rhythm-LAVI groups (**c**) and FB states (**d**). **e**, *ELF2* transcript abundance after vehicle or TGF-β1 ($n$ = 5). **f**–**h**, corresponding *KLF12* analyses. A two-sided Wilcoxon test was used in c,f and a paired *t*-test was used in e,h. *$P$ < 0.05; **$P$ ≤ 0.01.

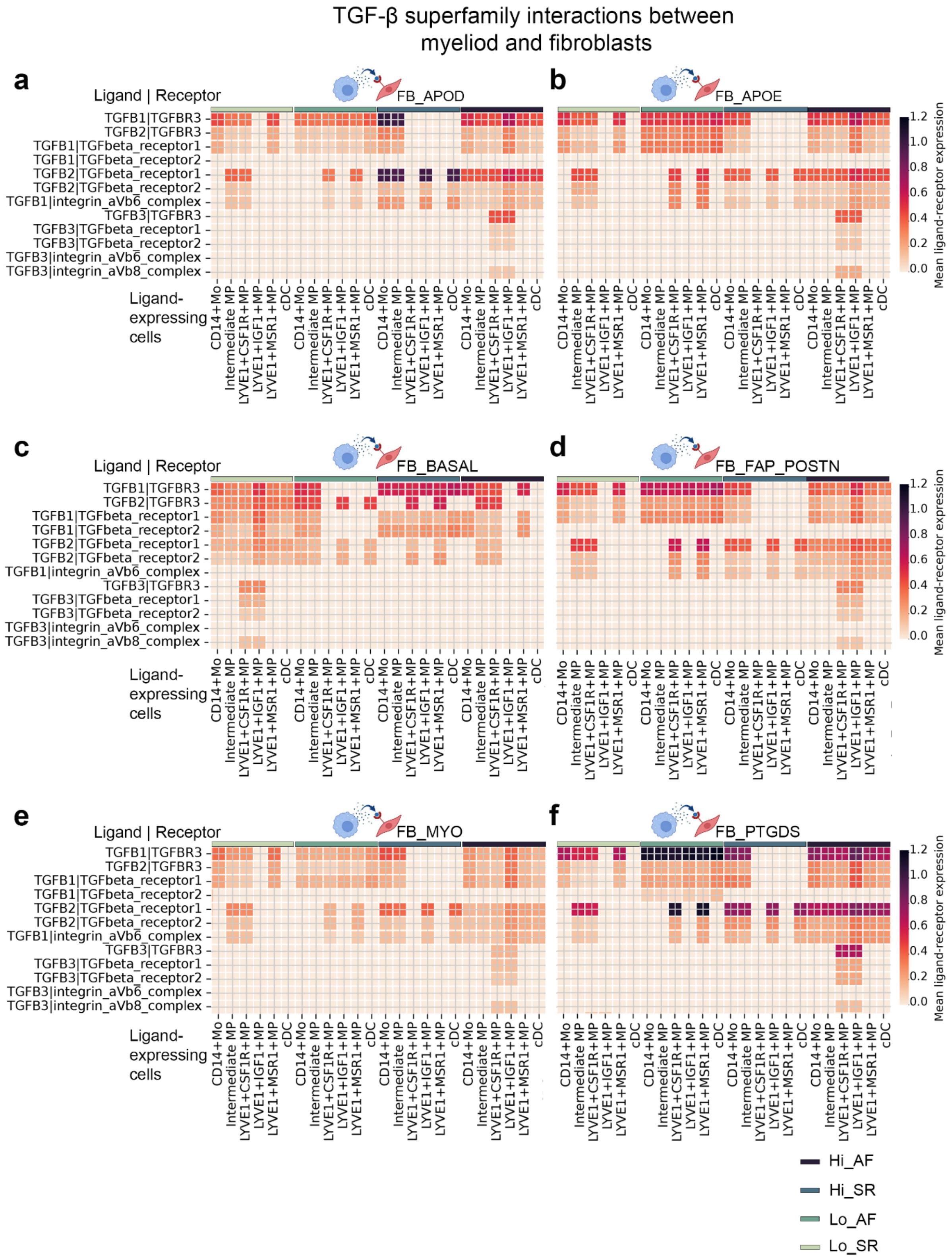


**Supplementary Fig. 20 | Predicted TGF-β signalling links myeloid states to FBs.** Heatmap of CellPhoneDB interaction means for TGF-β-superfamily ligand–receptor pairs between macrophage or dendritic-cell (DC) states and APOD$^{+}$ (**a**), APOE$^{+}$ (**b**), basal (**c**), FAP–POSTN$^{+}$ (**d**), myofibroblast (MYO) (**e**) and PTGDS$^{+}$ (**f**) states, stratified by Hi-AF, Hi-SR, Lo-AF and Lo-SR. Rows denote ligand–receptor pairs and columns sender–receiver combinations; higher colour intensity denotes a larger interaction mean.

#### Supplementary tables

**Supplementary table 1**. Patient demographics and clinical information

| | | LAA sequencing (n = 24) | RAA sequencing (n = 36) | Sequencing overlap (n = 14) |
|---|---|---|---|---|
| Characteristic | Statistic | Summary | Summary | Statistic |
| Age, years | Mean ± SD | 66.0 ± 8.0 | 69.5 ± 8.0 | 65.6 ± 8.6 |
| Female sex | n (%) | 5 (20.8%) | 9 (25.0%) | 4 (28.6%) |
| Atrial fibrillation | n (%) | 14 (58.3%) | 16 (44.4%) | 9 (64.3%) |
| LAVI, ml/m² | Mean ± SD | 36.9 ± 13.7 | 34.5 ± 10.9 | 33.4 ± 11.6 |
| BMI, kg/m² | Mean ± SD | 29.2 ± 5.3 | 29.6 ± 4.8 | 30.9 ± 5.6 |
| LVEF, % | Mean ± SD | 58.3 ± 9.5 | 56.7 ± 11.9 | 59.5 ± 9.5 |
| Arterial hypertension | n (%) | 16 (66.7%) | 25 (69.4%) | 10 (71.4%) |
| Diabetes mellitus | n (%) | 5 (20.8%) | 8 (22.2%) | 4 (28.6%) |
| Previous stroke | n (%) | 1 (4.2%) | 4 (11.1%) | 0 (0.0%) |
| Vascular disease | n (%) | 23 (95.8%) | 33 (91.7%) | 13 (92.9%) |
| HFrEF | n (%) | 1 (4.2%) | 5 (13.9%) | 1 (7.1%) |
| HFpEF | n (%) | 0 (0.0%) | 0 (0.0%) | 0 (0.0%) |
| CHF | n (%) | 1 (4.2%) | 5 (13.9%) | 1 (7.1%) |

LAA, left atrial appendage; RAA, right atrial appendage; HFrEF, heart failure with reduced ejection fraction; HFpEF, heart failure with preserved ejection fraction; CHF, chronic heart failure.
